# Organ on chip model of respiratory vascular interactions under COPD relevant oxidative stress

**DOI:** 10.64898/2026.06.04.730087

**Authors:** Maike Haensel, Rosie Millns, Harry Whitwell, Alexander J. Ainscough, Joseph van Batenburg-Sherwood, Lucas Breuil, Daria Kostuynina, Clare Lloyd, Beata Wojciak-Stothard

## Abstract

Oxidative stress-induced airway injury contributes to chronic obstructive pulmonary disease (COPD). Cardiovascular complications increase COPD morbidity and mortality, but mechanistic links between airway injury and vascular dysfunction remain unclear, largely due to limitations of *in vitro* models that fail to replicate the multicellular lung environment.

We developed REVAS, a modular organ-on-chip platform to study human respiratory-vascular cell-cell interactions at baseline and under oxidative stress conditions. REVAS consists of two respiratory chips hosting airway epithelium and microvascular endothelium, and a vascular chip hosting pulmonary artery endothelial cells co-cultured with vascular support cells, including smooth muscle cells, pericytes and fibroblasts. We studied effects of vascular support and respiratory cells on vascular endothelial phenotype at baseline and under H_2_O_2_-induced epithelial oxidative stress using functional assays, proteomic and transcriptomic analyses.

Multicellular environment enhanced vascular endothelial barrier function and promoted respiratory and vascular cell differentiation at baseline. Mural cells altered endothelial cell-matrix interactions, metabolism and cytoskeletal remodelling, while respiratory cells promoted endothelial aerobic respiration and quiescent phenotype.

Epithelial oxidative stress triggered inflammatory gene expression across all respiratory and vascular cells alongside apoptotic, reparative and pro-angiogenic signalling in endothelial and mural cells, accompanied by increased release of COPD-relevant cytokines and chemokines, including IL-6, TNF-α/β, IL-8, CCL5, CXCL9, PDGF, TGF-β. Comparative analyses with COPD endothelial datasets confirmed that REVAS recapitulates key features of disease-associated endothelial dysfunction.

These findings demonstrate that airway epithelial injury drives downstream vascular responses linked to inflammation and vascular remodelling, establishing REVAS as a human-relevant platform for mechanistic and therapeutic evaluation of cell-cell interactions in COPD and related lung diseases.

**Graphical Abstract:** 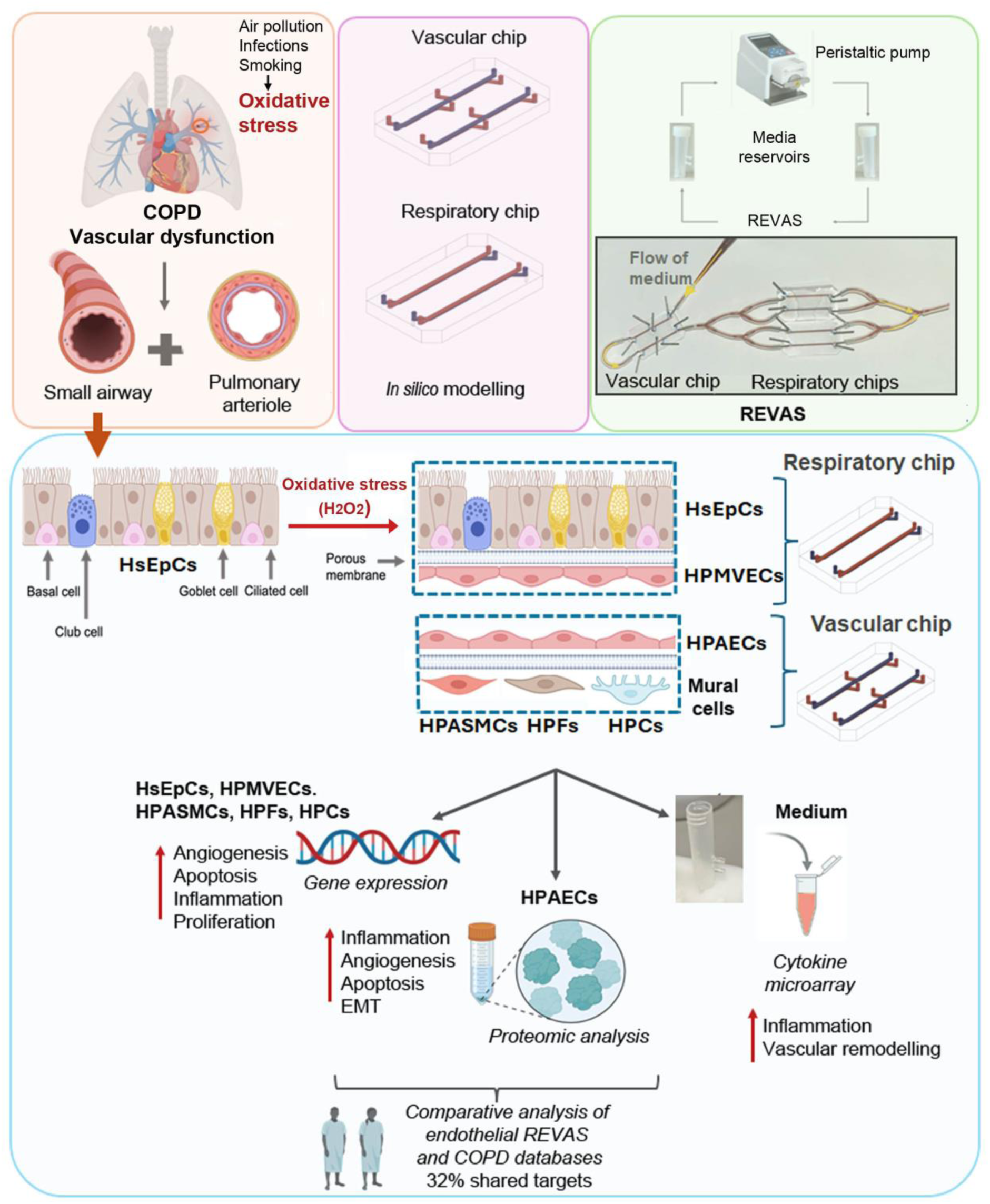

REVAS: a microfluidic platform developed to model multicellular interactions between airway epithelium and pulmonary vasculature under basal and oxidative stress. COPD: Chronic Obstructive Pulmonary Disease; EMT: endothelial-to-mesenchymal transition; HsEpCs: human small airway epithelial cells; HPMVECs: human pulmonary microvascular endothelial cells; HPAECs: human pulmonary artery endothelial cells; HPASMCs: human pulmonary artery smooth mucle cells; HPFs: human pulmonary fibcroblasts; HPCs: human pericytes.

## Introduction

COPD is the most prevalent respiratory disease and one of the top ten causes of death worldwide^1^. Oxidative stress induced by cigarette smoke, air pollution or infections plays a central role in driving COPD^2^ by promoting chronic inflammation, triggering cellular senescence and defective autophagy, reducing DNA repair capacity, enhancing autoimmune responses, increasing mucus production, and weakening the anti-inflammatory effects of corticosteroids^2–4^. COPD is defined by parenchymal destruction and small airway obliteration and is accompanied by profound vascular abnormalities, which are major contributors to disease progression and mortality^5^. These include loss of pulmonary microvasculature and remodelling of pulmonary arteries with medial and intimal hyperplasia, commonly resulting in mild to moderate pulmonary hypertension and, in some cases, a severe vascular phenotype. Microvascular rarefaction is thought to arise from dysregulated angiogenesis and endothelial apoptosis.^6^

Accumulating evidence identifies endothelial cells (ECs) as central regulators of vascular dysfunction in lung diseases, including COPD^7^. In response to airway injury, ECs release vasoconstrictive, pro-remodelling and proinflammatory mediators. These can drive vascular remodelling characterised by smooth muscle cell expansion, extracellular matrix deposition and neointimal formation, ultimately leading to pulmonary hypertension and worsened clinical outcomes^7^. Compromised endothelial barrier function and endothelial-to-mesenchymal transition (EMT) is also thought to play a contributory role^8^. Despite the importance of the respiratory-vascular crosstalk, mechanistic events coupling epithelial and vascular pathology in this disease are poorly understood.

Animal models and conventional *in vitro* systems provide important insights but do not fully recapitulate the pathology of human disease. Organ-on-chip (OOC) cell culture systems offer a promising alternative, although lung-on-a-chip models, focused primarily on alveolar or airway compartments^9,10^, have not captured integrated airway-vascular crosstalk.

To overcome this limitation, we developed REVAS, a modular organ-on-a-chip platform integrating microfluidic models of small airways and pulmonary arterioles. Using this platform, we investigated human respiratory-vascular cell-cell communication under basal conditions and following H_2_O_2_-induced oxidative epithelial injury. Functional, transcriptomic, and proteomic analyses revealed a profound impact of localised airway damage on pulmonary endothelial and vascular mural cell phenotypes, consistent with alterations observed in COPD. REVAS provides a mechanistically tractable, human-relevant platform to study early epithelial-vascular crosstalk in COPD and related lung diseases.

## RESULTS

### REVAS microfluidic platform design

The <u>RE</u>spiratory - <u>VAS</u>cular microfluidic platform (REVAS) was designed to enable the study of human respiratory-vascular cell-cell interactions. It comprises modular respiratory and vascular chips connected to a multichannel peristaltic pump, which drives airflow through the epithelial channels and medium flow through the endothelial channels, respectively. The dimensions of microchips were set to 20.0 mm in width and 32.0 mm in length (**Figure 1**), with microfluidic channels 200 μm in height and 1 mm in width, which falls within the range of sizes of small airways and arterioles in the lung^11^.

**Figure 1:**
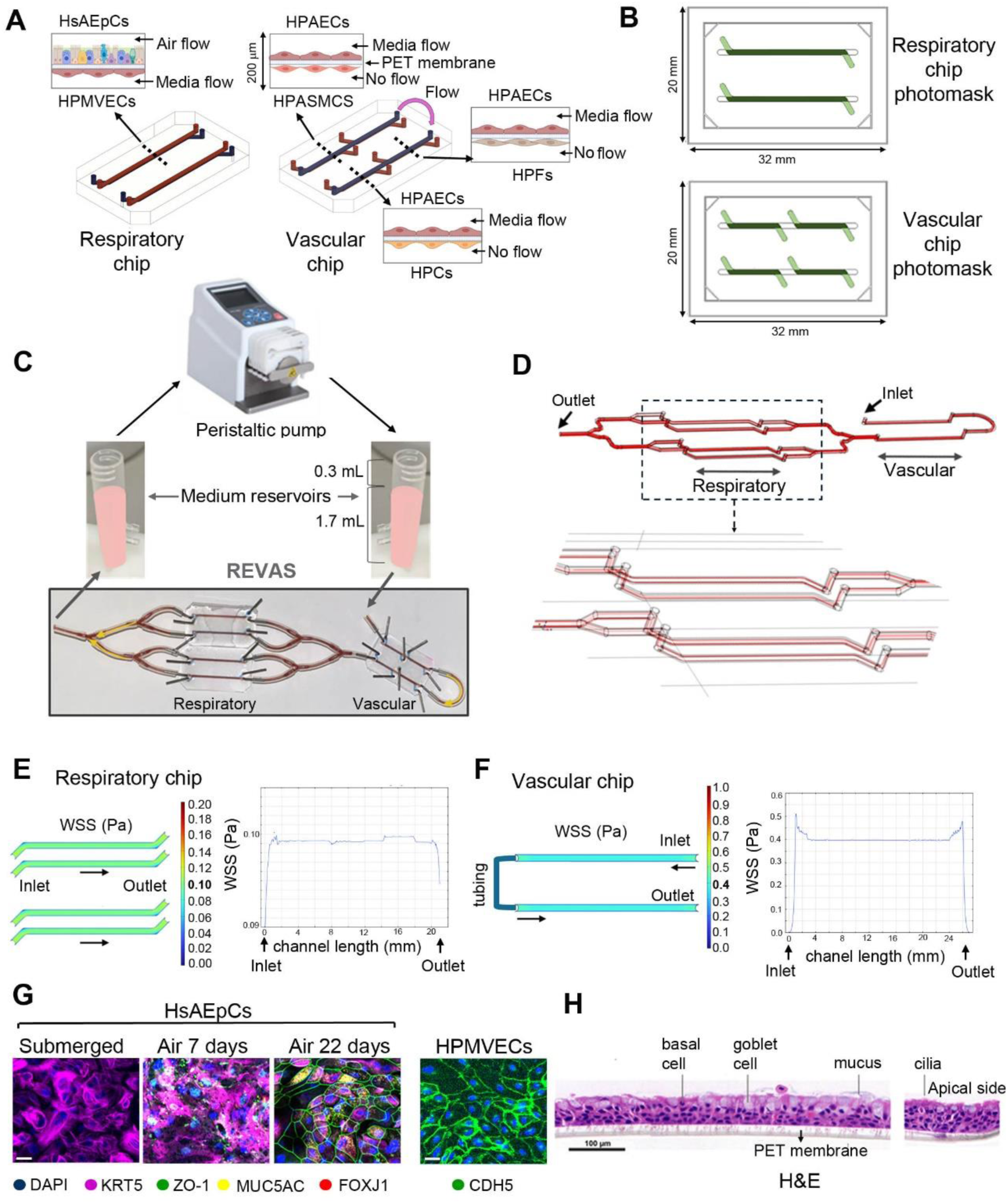
Overview of REVAS design. **(A)** Schematic diagram of the respiratory and vascular chips. Channel cross-sections (broken line) illustrate different cell types separated by a porous PET membrane in vascular and respiratory chips. (**B**) Photomask design for vascular and respiratory chips. The contact area between the top and bottom channels is shown in dark green. **(C)** Flow system schematic in REVAS; (**D)** Flow streamlines in REVAS microvascular channel with enlarged image of the boxed area below; **(E, F)** Wall shear stress (WSS) distribution in cross-sections of endothelial channels in the **(E)** respiratory and **(F)** vascular chips. **(G)** Human small airway epithelial cells (HsAEpCs) cultured under submerged and air-liquid interface conditions for 7 or 22 days, stained for nuclei (DAPI), *KRT5* (basal cells), *ZO-1* (tight junctions), *MUC5AC* (goblet cells), and *FOXJ1* (ciliated cells), as indicated. Right image: HPMVECs stained for adherens junctions marker, *CDH5*. Scale bar = 10 µm. **(H**) H&E staining (20×) of differentiated HsAEpCs cultured on porous PET membrane showing ciliated, basal, and mucus-producing goblet cells. Scale bar = 100 µm.

The respiratory chip comprises two channels, each divided into two parts by a semi-permeable porous (0.4µm pores) polyethylene terephthalate (PET) membrane (**Figure 1A**). The top channels of this chip are designed to host human small airway epithelial cells (HsAEpCs) whilst the bottom channels host human pulmonary microvascular endothelial (HPMVECs) cells cultured on the opposite side of the membrane, allowing direct cell-cell communication **(Figure 1A, B).**

The vascular chip consists of two upper channels lined with human pulmonary arterial endothelial cells (HPAECs) connected by tubing, and four transversely arranged lower channels separated from the top channels by a porous PET membrane. Three of the lower channels contain human pulmonary arterial smooth muscle cells (HPASMCs), human pulmonary fibroblasts (HPFs), and human pericytes (HPCs), which are collectively referred to as “mural cells” for simplicity **(Figure 1A).** Accordingly, cells grown in the respiratory chip are referred to as “respiratory cells,” whereas cells grown in the vascular chip are referred to as “vascular cells”.

All channels have separate access ports. The fourth channel was designed for potential incorporation of an additional cell type in future work. Microfluidic chips were produced using photolithography and soft lithography^12^. Photomask designs for respiratory and vascular chips are shown in **Figure 1B** and **Supplementary Figure 1.**

To mimic arteriolar branching, flow from the vascular top channels containing macrovascular endothelial cells (HPAECs) was divided into four equal streams using 0.76 mm inner diameter tubing equipped with 0.8 mm inner diameter flow splitters. Each stream was directed into one of four downstream microvascular endothelial channels (HPMVECs), distributed across two respiratory chips **(Figure 1C**). COMSOL Multiphysics simulations were used to determine flow rates required to achieve physiologically relevant wall shear stress (WSS). An inlet flow rate of 9.75 mL/hr in the vascular channel produced ∼4 dynes/cm² WSS in the HPAEC channel^13^. Due to flow splitting, downstream HPMVEC channels experienced a reduced flow rate of 2.44 mL/hr, corresponding to ∼1 dyne/cm² WSS, consistent with conditions in the lung microvasculature and in lung-on-chip models^14, 15,16^.

Velocity streamline analysis confirmed equal flow distribution across the downstream branches following flow splitting (**Figure 1D**). Simulations further demonstrated that shear stress was uniformly distributed across the channel length, with only minor local increases noted near channel edges and inlet/outlet regions (**Figure 1E, F**). Reynolds numbers remained below 2 across all regions, confirming laminar flow (mesh size: extra fine, Reynolds number: 1.15).

To reduce pulsatility generated by the peristaltic pump, the air-to-culture medium volume ratio was adjusted in the media reservoirs, so they functioned as pulse dampeners. Flow rate measurements with a thermal flow sensor showed that maximal reduction in flow oscillations was achieved when each reservoir contained 1.7 mL of medium and 0.3 mL of air **(Supplementary Figure 2)**.

### Cells cultured in REVAS retain differentiated phenotype

The number of cells in each channel was estimated by dividing the known channel surface area by the average spread area of each cell type, as determined from confocal image analysis. The top channel in the respiratory chip can host ∼ 17,000 HsAEpCs in contact with ∼11,000 HPMVECs cultured on the other side of the PET membrane in the bottom channel. The top channel of the vascular chip can host ∼10,000 HPAECs, which remain in contact with ∼2,000 HPASMCs, ∼3,000 HPFs, and ∼5,000 HPCs grown separately in transverse bottom channels. The size of total channel areas and regions of overlap between top and bottom channels, together with numbers of cells growing in each part are detailed in the **Supplemental Tables 1-6**.

Initially, we characterised respiratory and vascular chips separately, to confirm that cells cultured in them retained their differentiated phenotype. In the respiratory chip, HsAEpCs formed a confluent layer of basal cells under submerged culture conditions (**Figure 1F**). Air exposure for 7 days induced epithelial cell differentiation, characterised by increased gene-expression of airway epithelial markers including basal cells (*KRT5*, *TP63*), goblet cells (*MUC5AC*, *MUC5B*), ciliated cells (*FOXJ1*), club cells (*SCGB1A1*), and tight junction proteins (*TJP1-3*). Interestingly, expression of *MUC5AC* (P < 0.001), *MUC5B* (P < 0.0001) and *SCBG1A1* (P < 0.01) was significantly elevated in HsAEpCs co-cultured with HPMVECs compared with HsAEpCs monocultures **(Supplementary Figure 3)**. Prolonged, 22-day air exposure induced further differentiation of the airway epithelium, confirmed by the presence of cilia, basal and goblet cells (**Figure 1G, H**). Given that 7-day air exposure was sufficient to achieve a significant increase in epithelial differentiation and was experimentally more feasible, this timepoint was chosen in further experimentation.

HPMVECs co-cultured with epithelial cells showed a robust expression of differentiation markers *PECAM1* and *CDH5* (VE-cadherin) (**Supplementary Figure 3A**) and a cobblestone morphology with junctional localisation of VE-cadherin (**Figure 1G**).

HPAECs cultured in the vascular chip, showed flow-induced alignment (**Supplementary Figure 4**). Expression of differentiation marker genes in HPAECs, HPCs and HPASMCs cultured in microfluidic chip was significantly increased, compared with static monocultures, while HPFs remained relatively unaffected (**Supplementary Figure 4).**

### Vascular mural cells support endothelial quiescence and enhance endothelial barrier function

A fundamental question is whether endothelial cells would behave differently when co-cultured with other cellular components of the vascular wall. Organ-on-a-chip platforms enable the recreation of multicellular environments to investigate this directly. Here, we first evaluated the impact of vascular mural cells on barrier function and protein expression profile of HPAECs cultured in the vascular chip under basal, unstimulated conditions.

To evaluate effect of vascular mural cells on HPAEC barrier function, HPAECs were stimulated with thrombin, a known inducer of endothelial permeability, involved in thrombotic exacerbations of COPD^17^. The presence of vascular mural cells markedly attenuated thrombin- induced endothelial permeability (**Figure 2A**) and significantly increased expression of endothelial junctional proteins, *CDH5* and *CD31* (**Supplementary Figure 4**).

**Figure 2:**
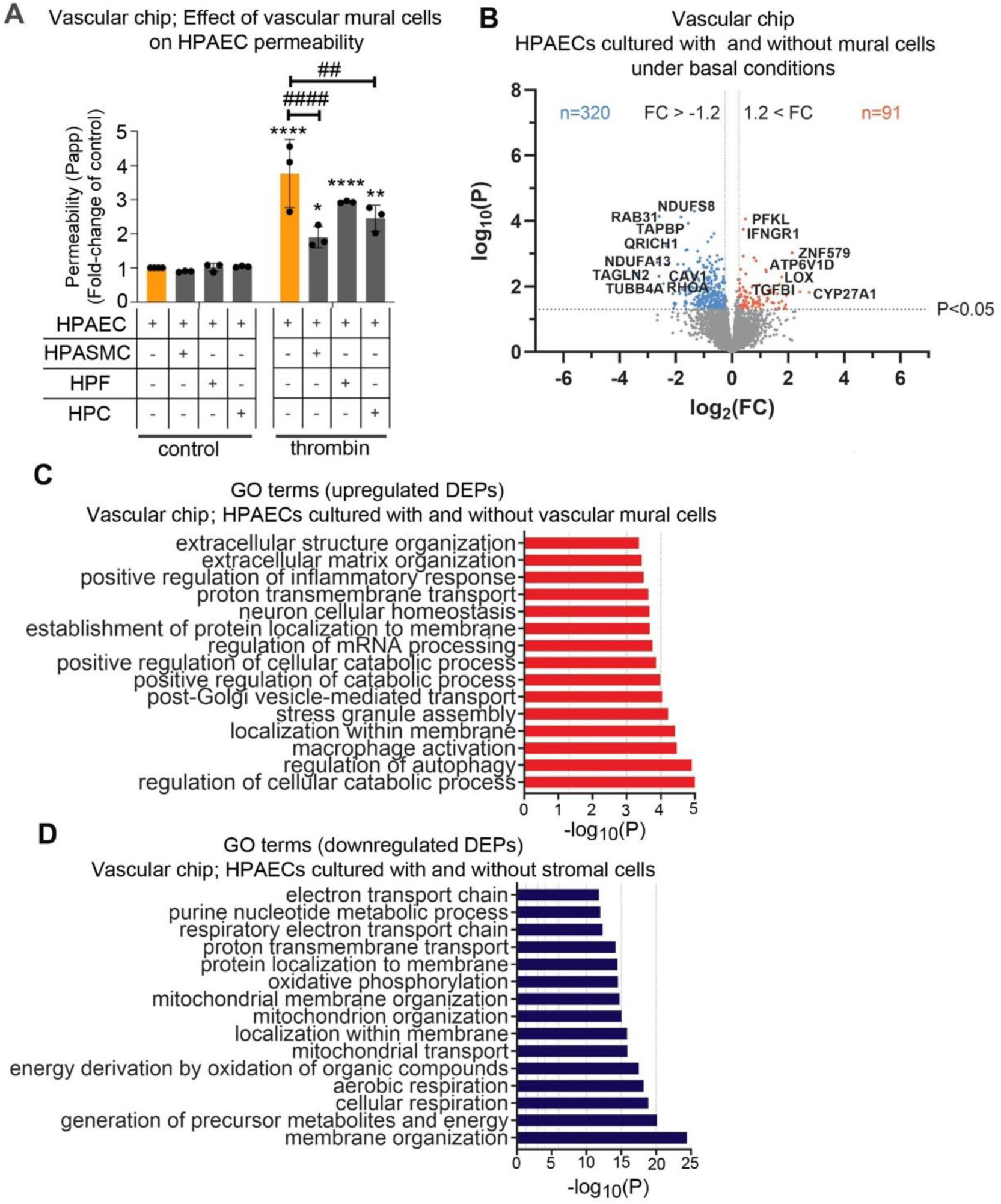
Effect of vascular mural cells on endothelial barrier function and protein expression profile in HPAECs cultured in the vascular chip. **(A)** Permeability changes in HPAECs cultured alone or co-cultured with HPASMCs, HFs, HPCs, with or without thrombin (1 U/mL, 1hr), as indicated. Passage of FITC-Dextran (1 mg/mL) was used to measure endothelial permeability. The apparent permeability was calculated using an equation based on Takeshita et al. ^14^. Error bars indicate mean ± SEM; two-way ANOVA with Tukey’s post-hoc correction test; *P < 0.05, **P < 0.01, ****P < 0.0001, comparison with static (Transwell) control; ^##^P < 0.01, ^####^P < 0.0001, comparison with thrombin endothelium-only control, as indicated. n=3. **(B**) Volcano plot shows differentially expressed proteins (DEPs) in HPAECs cultured alone vs HPAECs co-cultured with other vascular cell types in the vascular chip. Downregulated DEPs (blue), upregulated DEPs (red); P < 0.05; log_2_ FC <-1.2 or >1.2. n=5. **(C, D**) Metascape pathway and functional enrichment analysis of **(C)** upregulated and **(D)** downregulated GO protein-associated pathways. Bar graph shows Metascape pathway and process enrichment analysis of top 15 most upregulated protein-associated pathways of HPAECs cultured alone vs in co-culture with other vascular cell types. DEP selection criteria were FDR < 0.05, log_2_ FC <-1.2 or >1.2. n=5.

To better understand the impact of a multicellular vascular environment on the HPAEC phenotype at baseline, we performed an unbiased proteomic analysis comparing HPAECs cultured alone with those maintained in co-culture with HPASMCs, HPFs, and HPCs. This approach is novel in the context of lung-on-chip systems, which typically rely on transcriptomic analyses^18,19^.

This analysis identified 91 upregulated and 320 downregulated differentially expressed proteins (DEPs) (P < 0.05; 1.2 > log_2_ FC < -1.2) (**Figure 2B).** GO enrichment analysis (Metascape) revealed that the upregulated DEPs were significantly enriched in processes related to extracellular matrix (ECM) organisation, endothelial plasticity, immune signalling, and vesicular transport (**Figure 2C).** Downregulated DEPs were predominantly associated with mitochondrial respiration, signal transduction, and cytoskeletal organisation, likely reflecting the adoption of a low metabolic demand, quiescent endothelial phenotype. GO pathway enrichment analysis with lists of upregulated and downregulated DEPs in HPAECs cultured with vascular mural cells versus HPAECs cultured alone in a vascular chip are listed in the **Supplementary Tables 7 and 8.**

### Respiratory cells stimulate endothelial aerobic energy metabolism and modulate expression of vascular stability regulators in mural cells

The next step was to investigate the impact of cells from the respiratory chip (HsAEpCs + HPMVECs), on cells from the vascular chip (HPAECs + vascular mural cells) co-cultured under basal, unstimulated conditions. To this end, two respiratory chips were connected to the vascular chip to form a REVAS circuit, and the endothelial protein expression profile of HPAECs was analysed after 48 hrs of co-culture under flow.

Proteomic analysis revealed substantial alterations in the protein expression profile in HPAECs cultured in a complete REVAS circuit, compared with HPAECs cultured in the vascular chip alone. In total, 354 DEPs were upregulated and 188 were downregulated (P < 0.05; log₂ fold change > 1.2 or < -1.2) **(Figure 3A)**. Upregulated DEPs were significantly enriched in oxidative phosphorylation, aerobic respiration, and the electron transport chain pathways **(Figure 3B**). Notably, there was a marked upregulation in expression of mitochondrial and respiratory chain components, including subunits of complex I (e.g., NDUFA9, NDUFS1/2), complex II (SDHA/B), ATP synthase (ATP5F1A/B/C), and cytochrome c oxidase (COX5A/B, CYCS).

**Figure 3.**
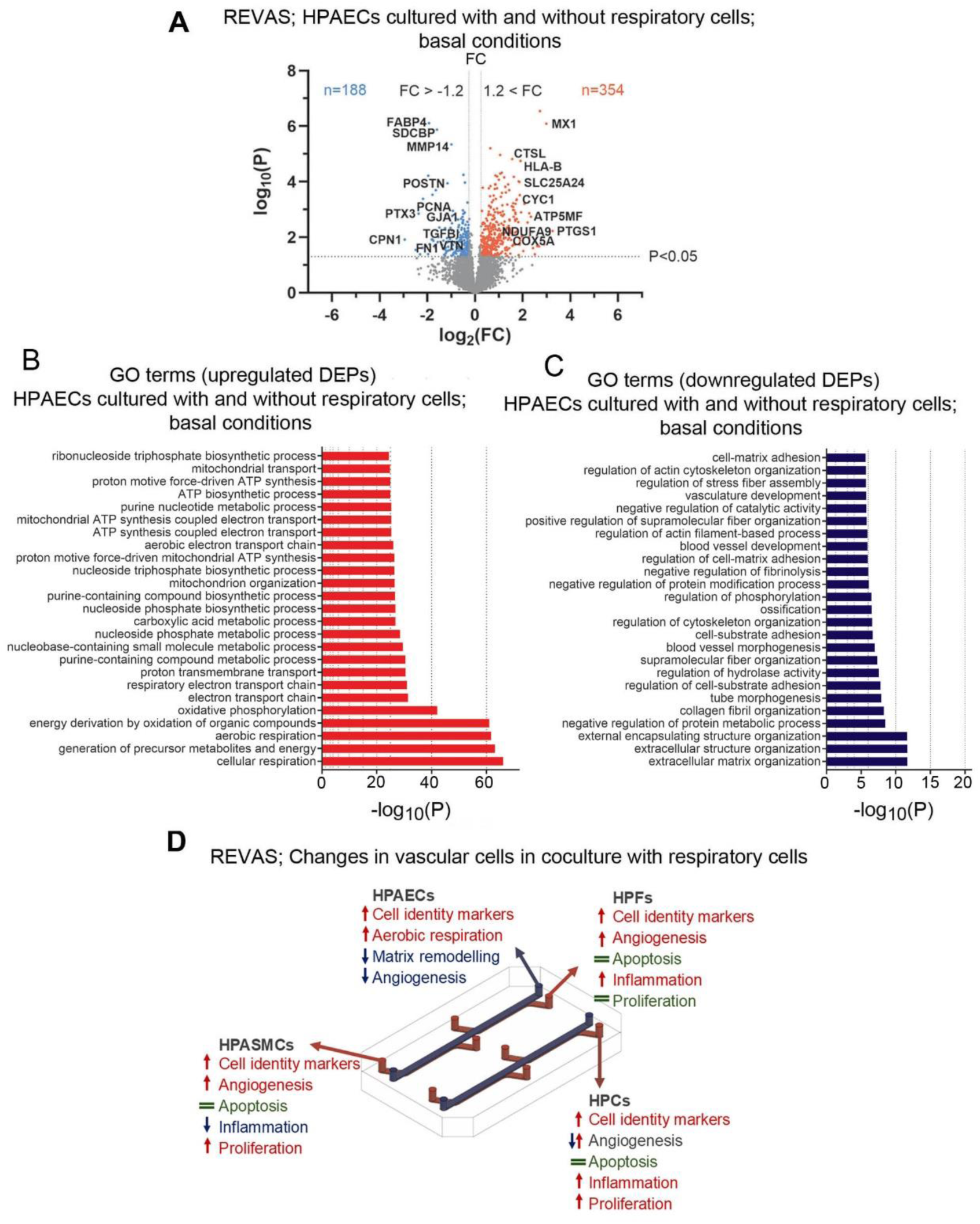
Effect of respiratory cells on vascular endothelial and mural cells in REVAS. **(A)** Volcano plot of differentially expressed proteins (DEPs) in HPAECs cultured in REVAS (respiratory + vascular chip) versus the vascular chip alone. Downregulated DEPs are shown in blue, upregulated DEPs in red. **(B)** Metascape pathway and functional enrichment analysis of upregulated protein-associated pathways. Bar graph shows the top 15 most significantly enriched GO pathways in HPAECs within REVAS. **(C)** Metascape pathway and functional enrichment analysis of downregulated protein-associated pathways. Bar graph shows the top 15 most significantly enriched GO pathways among downregulated DEPs. Analysis was performed using Metascape, based on GO biological processes. DEP selection criteria: FDR < 0.05, log_2_ FC < -1.2. n = 5. **(D)** Summary of responses observed in endothelial and vascular mural cells induced by contact with respiratory chip under basal conditions (corresponding to data in A-C and Supplementary Figures 5-7). The figure was made using PowerPoint (Microsoft) and Fusion360 (Autodesk).

In contrast, downregulated DEPs were enriched in pathways associated with ECM remodelling, cytoskeletal organisation, vascular morphogenesis, and cell-matrix adhesion (**Figure 3C**). Key ECM and structural proteins, including COL5A2, LOX, MMP14, TGFBI, FN1, VTN, POSTN and GJA1, which are typically linked to enhanced endothelial proliferation, migration, and wound healing^20–23^, were significantly downregulated. The top 25 enriched pathways, together with their associated up- and downregulated DEPs, are shown in **Supplementary Tables 9 and 10.**

In summary, co-culture of HPAECs with respiratory cells under basal, unstimulated conditions promoted endothelial metabolic shift towards aerobic energy production and reduced responses associated with proliferation and vascular remodelling.

Although our study primary focused on endothelial phenotype, we also performed a preliminary assessment of changes induced by respiratory cells in vascular mural cells. Specifically, we analysed the expression of selected marker genes related to cell identity, angiogenesis, apoptosis, inflammation, and proliferation (**Supplementary Figures 5-7**).

Overall, vascular mural cells cultured within REVAS exhibited increased expression of differentiation markers, *CNN1* and *TAGLN* in HPASMCs; *COL1A1* and *PDGFRα* in HPFs; and *NG2* and *PDGFRβ* in HPCs. Expression of angiogenesis regulators, including *ANG1*, *VEGFA*, and *POSTN*, was also elevated. Apoptosis-related markers (*BAK1*, *CASP9*) were unchanged. In contrast, inflammatory and proliferation markers showed cell type-dependent modulation. In HPASMCs, inflammatory mediators (*VCAM1*, *ICAM1*, *CCL5*, *IL6*) were reduced, whereas their expression increased in HPFs and HPCs. Proliferation marker genes (e.g., *BRD4*, *CDC45*) displayed variable changes in expression across different cell types. The overall effects of respiratory cells on HPASMCs, HFs, and HPCs are summarised in **Figure 3D**.

### Airway epithelial oxidative stress induces apoptotic, inflammatory, and reparative responses in vascular cells

Oxidative stress caused by cigarette smoke, air pollution, and respiratory infections is a major driver of lung epithelial injury and chronic inflammation in COPD^3^. To induce oxidative stress in our model, we used hydrogen peroxide (H_2_O_2_), a COPD biomarker^2^, widely used in *in vitro* modelling of oxidative stress-induced airway epithelial damage^24^.

In exposure optimisation studies, treatment of HsAEpCs with H₂O₂ (1-100 µM, 1 hr) induced a dose-dependent apoptotic response (**Supplementary Figure 8**). 20 µM H_2_O_2_ was the lowest concentration inducing measurable changes in the expression of apoptosis marker *BAK1* and inflammatory marker *ICAM1* without excessive loss of the epithelial monolayer (**Supplementary Figures 8 and 9**) and therefore this concentration was selected for further experiments. Considering that H_2_O_2_ rapidly degrades in culture (with half-life ∼30 minutes) ^25,26^, and that air-liquid interface (ALI) cultures pose challenges for uniform H_2_O_2_ exposure^27^, we used a 1-hour H_2_O_2_ treatment under submerged conditions in HsAEpCs, an approach commonly used in other respiratory epithelial injury models^28,29^.

H_2_O_2_ triggered robust inflammatory and apoptotic transcriptional responses in HsAEpCs (**Supplementary Figure 10**) and HPMVECs (**Supplementary Figure 11**). This was evidenced by increased expression of pro-apoptotic genes (*BAK1*, *CASP9*) and inflammatory markers (*VCAM1*, *CCL5*, *IL6*). In HPMVECs, *VEGFA* expression was also significantly upregulated, potentially reflecting an early reparative or angiogenic response to oxidative injury.

Our primary objective was to evaluate vascular cell response to oxidative stress-induced epithelial damage in REVAS. Specifically, we investigated whether epithelial injury can alter the behaviour of endothelial and vascular mural cells within the vascular chip. A secondary objective was to determine how vascular mural cells modulate endothelial responses in this environment.

Following exposure of HsAEpCs to H_2_O_2_ for 1 hour, the respiratory chip was connected to the vascular chip. To further exclude the possibility that residual H_2_O_2_ directly affected the cells in the vascular chip, HsAEpCs were either left without a media change or were thoroughly washed with fresh medium before setting up a co-culture. No significant difference in apoptotic response was observed in HPAECs cultured under either condition, confirming that direct carryover of H_2_O_2_ did not contribute to HPAEC activation (**Supplemental Figure 12).** After 48 hrs of co-culture in REVAS, all vascular cell populations were harvested for subsequent proteomic (HPAECs) and transcriptomic (vascular mural cells: HPASMCs, HPFs, HPCs) analyses, whilst REVAS culture medium was collected for quantification of circulating cytokines. The overall experimental design of oxidative injury model with output measurements is summarised in **Figure 4A**.

**Figure 4.**
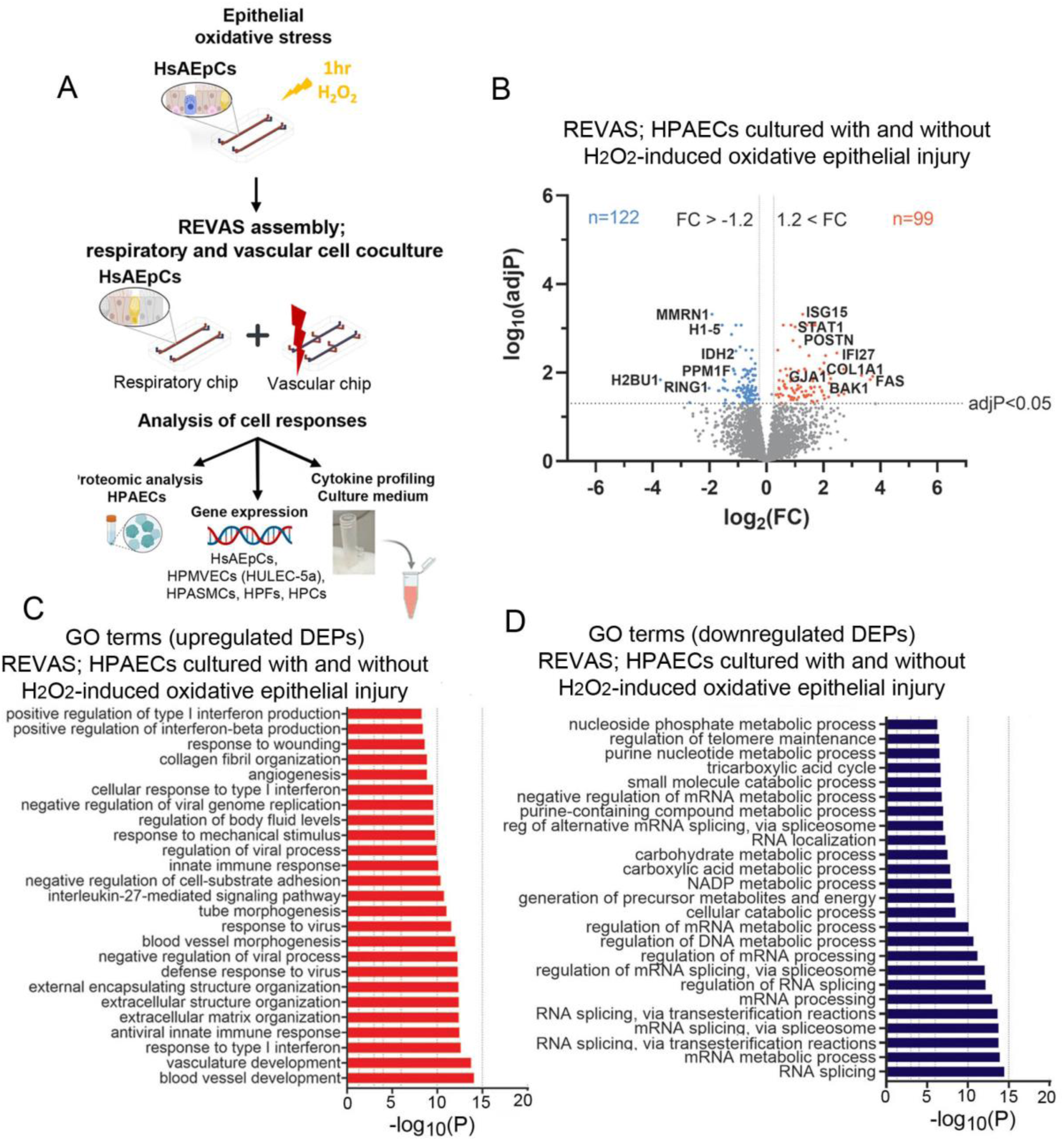
Effect of HsAEpCs oxidative injury on HPAECs. **(A)** Airway epithelial injury model in REVAS: HsAEpCs cultured in the respiratory (RE) chip were exposed to 20 µM H₂O₂ for 1 hour to induce epithelial injury. After, the RE chip was connected to the vascular (VAS) chip, which contained vascular cell types (HPAECs, HPASMCs, HPFs, HPCs) in co-culture. The connected REVAS circuit was perfused for 48 hours, after which HPAECs were collected for downstream proteomic analysis **(B)** Volcano plot shows differentially expressed proteins (DEPs) in HPAECs co-cultured with H₂O₂-treated airway epithelium in REVAS, compared with corresponding untreated controls. Red dots: upregulated DEPs, blue dots: downregulated DEPs (adj P < 0.05; log₂ FC > 1.2 or < -1.2); n = 5. **(C, D)** Bar charts show 25 most significantly enriched GO Biological Processes for upregulated and downregulated DEPs, respectively (Metascape).

Proteomic analysis of HPAECs cultured in REVAS under respiratory epithelial stress conditions identified 221 DEPs, including 122 downregulated and 99 upregulated DEPs (Padj < 0.05; log₂ fold change >1.2 or <-1.2) (**Figure 4B**).

The upregulated DEPs were enriched in innate immunity, extracellular remodelling, and vascular development pathways **(Figure 4C).** Notable protein targets included STAT1, IFI27, ISG15, and TGFBR2 (inflammation and immune activation), BAK1 and IFI27 (apoptosis); COL1A1, CCN2, MMP14, POSTN (structural reorganization and matrix remodelling), ENG, SERPINE1, THBS1, and ECGF1 (wound healing and angiogenesis) and GJA1 (cell-cell communication). The downregulated DEPs were associated with RNA splicing, processing, mRNA metabolism as well as tricarboxylic acid cycle, NADP metabolic processes, and carboxylic acid metabolism (**Figure 4D**). Top 25 pathways with associated upregulated and downregulated DEPs are listed in **Supplementary Tables 11 and 12.**

Response of HPASMCs, HPFs and HPCs to respiratory epithelial oxidative damage was studied using selected marker genes of cell identity, angiogenesis, apoptosis, inflammation, and proliferation (**Supplementary Figures 13-15**). Most prominent changes were noted in HPFs, which exhibited a significant downregulation of fibroblast identity markers, including *COL1A1*, *PDGFRα*, and *VIM*, increased expression of immune (*VCAM1*, *CCL5*), apoptotic (*BAK1*, *CASP9*), angiogenic (*POSTN*) and proliferative (*BRD4*) marker genes. HPASMCs showed upregulation of *POSTN* and *ACTA2*, whilst HPCs were relatively less affected, showing only small changes in the expression of apoptosis markers *BAK1* and *CASP9*, leukocyte adhesion receptors *ICAM1* and *VCAM1* and proliferation marker *BRD4*.

### Vascular mural cells enhance endothelial reparative response to epithelial oxidative injury

Our earlier results demonstrated that the presence of vascular mural cells significantly influenced the HPAEC phenotype under basal, unstimulated conditions. Next, we investigated how the vascular mural cells affect HPAEC response to epithelial oxidative stress. To address this, we compared the protein expression profiles of HPAECs cultured with or without other vascular cell types under oxidative epithelial stress conditions in REVAS. Proteomic analysis identified 438 differentially expressed proteins (196 upregulated and 242 downregulated; Padj < 0.05; log_2_ FC >1.2 or <-1.2) in HPAECs co-cultured with mural cells compared with HPAECs cultured without mural cells in this model (**Figure 5A).**

**Figure 5.**
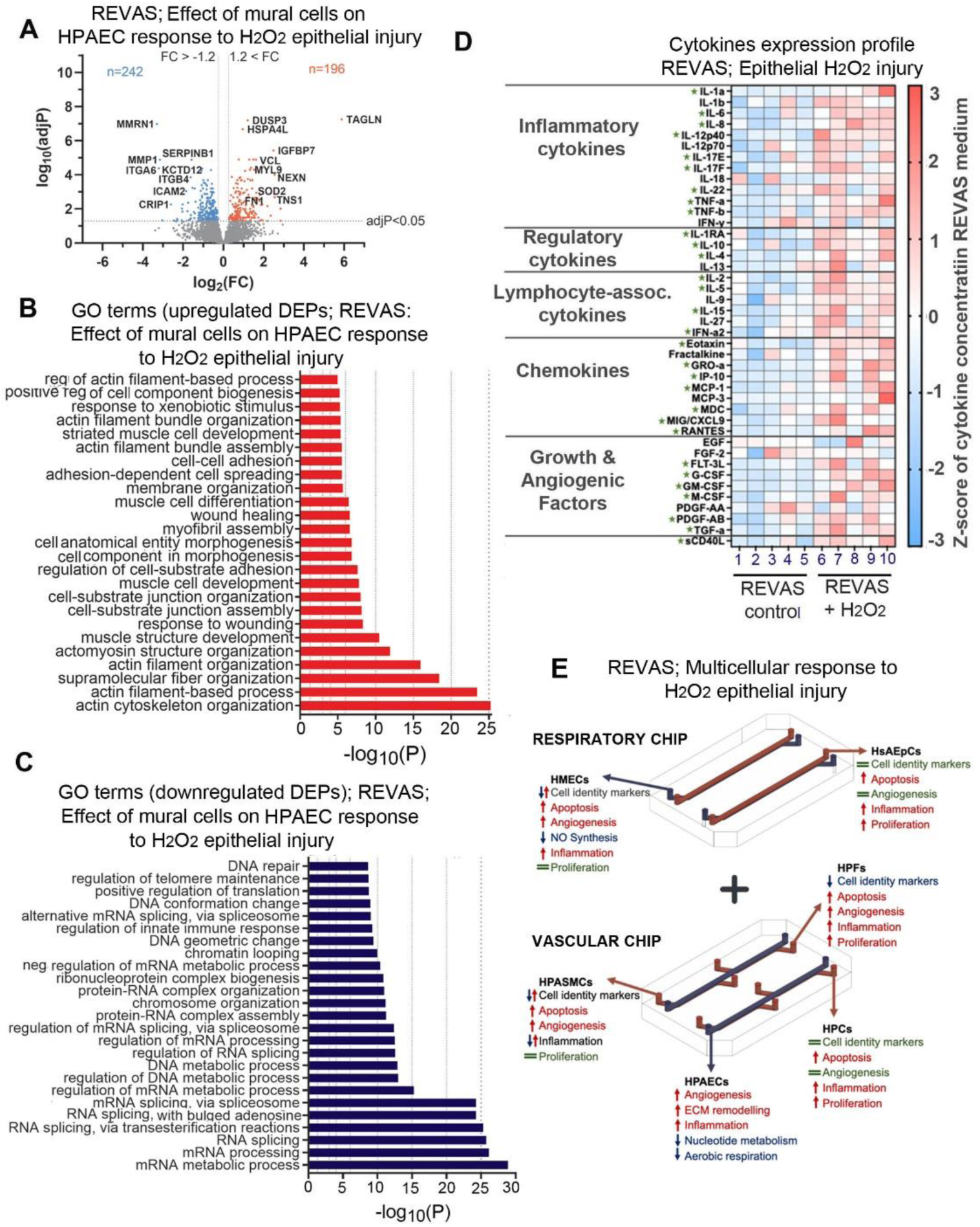
Effects of vascular mural cells on HPAEC response to epithelial oxidative injury and cytokine profiling in REVAS. **(A)**Volcano plot displaying DEPs in HPAECs cultured with, or without vascular mural following epithelial H₂O₂ exposure. Blue dots are downregulated DEPs; red dots are upregulated DEPs (Padj < 0.05; log_2_ FC > 1.2 or < −1.2) (n=5 **(B)** Bar chart shows top 20 most significantly enriched GO Biological Processes for upregulated DEPs; **(C)** 20 most significantly enriched GO Biological Processes for downregulated DEPs**. (D)** Heatmap showing changes in cytokine levels in REVAS culture medium 48 hours following H_2_O_2_ exposure (48-plex human cytokine assay). Z-scores were calculated for each cytokine across samples to allow for standardised comparison. n=5/treatment group. Red indicates increased expression relative to the mean (positive z-score), blue indicates decreased expression (negative z-score). **\***marks statistically significant change in cytokines levels; unpaired t-test; n=5. **(E)** Summary of H_2_O_2_-induced changes in cells cultured in REVAS, based on proteomic (HPAECs) and transcriptomic analyses (HsAEpCs, HPMVECs, HPASMCs, HPFs and HPCs).

Pathway enrichment analysis of upregulated DEPs showed that the presence of vascular mural cells increased expression of proteins associated with cytoskeletal remodelling, adhesion, and contractility in HPAECs following H_2_O_2_-induced airway epithelial injury. Enriched terms included actin cytoskeleton organisation, cell-substrate junction assembly, membrane organisation, and wound healing, as well as muscle cell development, myofibril assembly, and striated muscle cell differentiation (**Figure 5B**). Conversely, downregulated DEPs were predominantly involved in RNA splicing, chromatin remodelling, and transcriptional regulation (**Figure 5C**). Top 25 GO pathways together with associated DEPs are shown in **Supplementary Tables 13 and 14.**

These results emphasise the importance of vascular mural cells as modulators of pulmonary endothelial response to airway injury, likely to act as enhancers of vascular repair.

### Cytokine and chemokine release induced by epithelial oxidative injury in REVAS

To complement proteomic and transcriptomic analyses, we also performed cytokine profiling (48-plex human cytokine assay) of media collected from REVAS flow circuit 48 hrs after H₂O₂-induced epithelial injury. Changes in the expression of cytokines grouped into different functional categories are illustrated in heatmap in **Figure 5D**, with cytokines that were significantly altered marked with an asterisk. Notable trends, consistent with the results of proteomic profiling, included a significant upregulation of inflammatory cytokines (IL-6, IL-8/CXCL8, TNF-α/β, IFN-γ) and chemokines (CXCL9/MIG, CCL2/MCP-1, CCL5/RANTES), alongside increased secretion of factors promoting vascular remodelling (TGF-β, PDGF).

Collectively, these findings demonstrate that the REVAS system elicits a robust multicellular paracrine response to epithelial oxidative injury. Injury within REVAS triggers a coordinated vascular endothelial program marked by inflammatory activation, extracellular matrix remodelling, and pro-angiogenic signalling. Concurrently, vascular mural cells exhibit cell type-specific alterations in inflammatory, apoptotic and proliferative activity, highlighting the integrated and dynamic nature of the multicellular response.

A summary of cell responses to epithelial oxidative injury in REVAS is shown in **Figure 5E**.

### Relevance of REVAS in disease modelling

DisGeNET database analysis^30^ revealed significant associations of HPAEC DEPs from epithelial oxidative injury model in REVAS with Vascular Diseases, Pulmonary Arterial Hypertension (PAH), Fibrosis, Scleroderma and Idiopathic Pulmonary Arterial Hypertension (**Figure 6A**). Examples of proteins driving these associations include mediators of inflammation (STAT1, IFI27)^31,32^, fibrosis (POSTN, TGFBR2) ^33,34^, apoptosis (BAK1, IFI27) ^35,36^, and vascular remodelling (ENG, TGFBR2, POSTN) ^37–39^.

**Figure 6.**
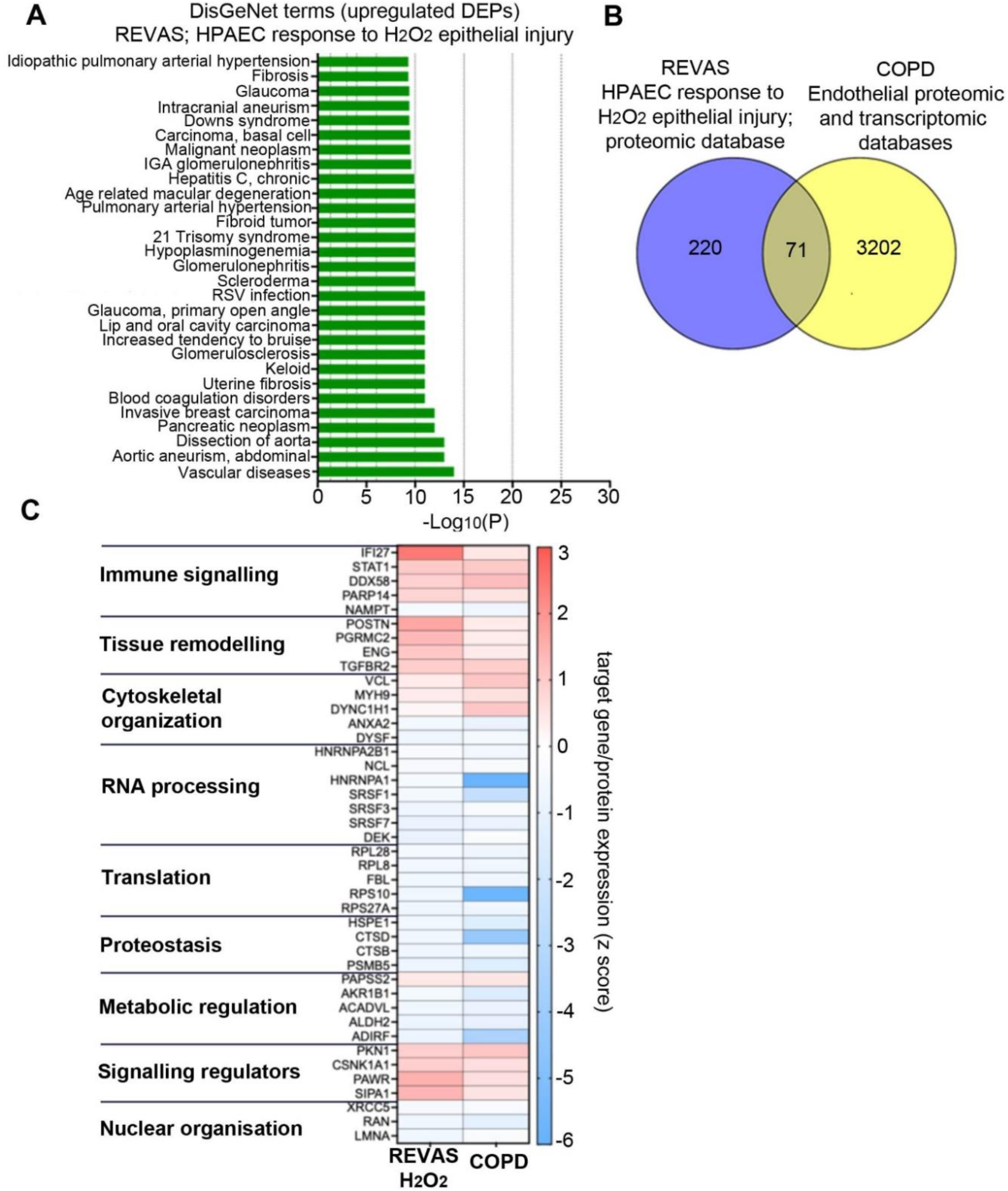
Comparative validation of the REVAS oxidative injury dataset against COPD and disease-associated datasets. **(A)** DisGeNET disease enrichment analysis of DEPs from HPAECs following epithelial oxidative injury. Bar plot shows 25 top disease associations. DEP selection criteria: Padj < 0.05; log_2_ FC > 1.2 or < −1.2; n=5. **(B)** Overlap between REVAS and COPD lung endothelial databases. Venn diagram illustrates the number of DEPs in HPAECs from the REVAS H₂O₂ injury model compared to proteomic endothelial COPD dataset and differentially expressed endothelial genes identified by single-cell RNA sequencing in COPD lungs. **(C)** Heatmap of proteins exhibiting concordant differential expression between REVAS and COPD endothelial datasets (n=71). Each row represents a shared protein/gene; columns represent experimental conditions. Red indicates upregulation and blue indicates downregulation, relative to respective controls. Differential expression was determined using Bonferroni-adjusted P-value < 0.05. The COPD log₂ fold change values were calculated from averaged expression across all endothelial subtypes.

To further evaluate the potential translational relevance of REVAS in modelling disease-relevant endothelial stress responses, the HPAEC DEP dataset from REVAS oxidative injury model (n = 220) was cross-referenced with proteomic endothelial COPD dataset^40^ and two endothelial single-cell RNA seq datasets from COPD lung tissues^41,42^. Datasets were filtered using an adjusted p-value threshold (Padj < 0.05) to ensure statistical stringency.

Overall, 71 out of 220 DEPs from the REVAS oxidative injury model (32 % of the REVAS dataset) were shared with the COPD endothelial proteomic and transcriptomic datasets **(Figure 6B)**. **Figure 6C** shows DEGs and DEPs that are upregulated or downregulated in the same direction across all studied datasets. **Supplementary Results Table 15** lists the shared differentially expressed gene and protein targets.

Numerous targets involved in inflammatory response, tissue remodelling/endothelial-to-mesenchymal transition (EndMT), cytoskeletal remodelling, and transcriptional/translational regulation showed similar direction of changes **(Figure 6C**). Notably, inflammatory mediators STAT1 and IF127 were upregulated, consistent with activation of cytokine- and interferon-driven signalling pathways^43,44^. Increased expression of proteins involved in extracellular matrix regulation and vascular remodelling, including TGFBR2^45^, ENG^46^ and POSTN^47^, further supports convergence with COPD-associated vascular changes. Markers of endothelial adhesion and structural integrity, particularly VCL and MYH9^48^, were similarly upregulated in both REVAS and COPD datasets, suggesting conserved alterations in cell-cell and cell-matrix interactions. Additionally, downregulation of RNA-binding and splicing-associated proteins (HNRNPA1, RPL28, RPS10, and SRSF3) in both contexts’ points to shared perturbations in RNA metabolism.

Despite these similarities, some distinctions emerged. For example, protein expression of inflammatory marker ICAM1^49^ was increased in HPAECs in oxidative injury model in REVAS but ICAM-1 mRNA levels were downregulated in COPD endothelium. Conversely, proteins associated with cytoskeletal dynamics and cell migration, including AHNAK ^50^ and MACF1^51^ were reduced in REVAS endothelium but their mRNA levels were elevated in COPD **(Supplementary Table 15).**

Taken together, these findings show that endothelial and vascular mural cell response to epithelial oxidative stress modelled in REVAS reproduces important features of vascular dysfunction in COPD. This underscores the utility of REVAS as a model for investigating COPD-relevant oxidative stress-induced vascular injury and identifying potential therapeutic intervention targets.

## DISCUSSION

We developed REVAS, a modular microfluidic platform that integrates respiratory and vascular units within a single perfused circuit, incorporating six human cell types: small airway epithelial, microvascular endothelial, smooth muscle, fibroblast, pericyte, and arterial endothelial cells. Compared with conventional binary co-culture systems, such as endothelial-smooth muscle or endothelial-fibroblast models^12,52^, this configuration enables simultaneous observation of multiple respiratory and vascular cell types under controlled physiological and pathological conditions while maintaining modularity for targeted analyses.

Cellular crosstalk is important for the lung development and regulation of tissue homeostasis within the respiratory system^53,54^. REVAS data show that multicellular microenvironments play a critical role in respiratory and vascular cell differentiation, consistent with observations *in vivo* and in physiologically relevant *in vitro* models^53,55,56^. In the respiratory chip, co-culture of HsAEpCs with HPMVECs significantly increased the expression of mucus-secreting and club cell markers in HsAEpCs, supporting the role of endothelial cells in driving epithelial differentiation^57,58^. HPAECs co-cultured with vascular mural cells exhibited enhanced expression of endothelial identity markers and improved endothelial barrier integrity compared with endothelial monocultures, reaffirming the protective role of mural cells on the endothelial stability and function^59–62^.

Given the critical role of the pulmonary vascular endothelium in the maintenance of lung homeostasis and response to injury^63^, we focused on characterising changes in HPAEC protein expression profile at baseline and under H_2_O_2_-induced epithelial (HsAEpCs) oxidative stress conditions. Endothelial responses were studied using proteomic analysis, which is a novel approach in lung-on-chip systems, which typically rely on transcriptomic analyses^18,19^. The advantage of proteomic profiling is that it provides a functional readout of cellular state by capturing post-transcriptional regulation and pathway activity that may not be apparent at the transcript level.

HPAECs showed upregulation of differentiation markers and pathways associated with aerobic respiration, extracellular matrix (ECM) organisation, cytoskeletal remodelling, immune signalling, metabolism, and vesicular transport. This response is likely to reflect a cross-talk between airway basal cells and the endothelium *in vivo*, known to support and sustain basal cell growth^64^.

In vascular mural cells, integration with the respiratory chip enhanced the expression of cell identity and angiogenesis markers in all cell types. Interestingly, a degree of inflammatory activation was observed in HPFs and HPCs, alongside selective upregulation of proliferation markers, most notably in HPCs. This partial “alert” state may result from paracrine signalling between airway basal cells and vascular cells and may play a role in supporting airway homeostasis, tissue regeneration and priming the system for potential immune challenges^64,65^.

Following analysis of respiratory-vascular cell interactions at baseline, we investigated response of respiratory and vascular cells to H_2_O_2_- induced HsAEpC oxidative injury. In the respiratory chip, H_2_O_2_ triggered a robust inflammatory and pro-apoptotic response in both HsAEpCs and HPMVCs, characterised by markedly upregulated expression of *BAK1*, *IL6*, *VCAM1*, and *CCL5*. The cells also showed upregulated levels of *VEGFA*, suggesting activation of a reparative response, also seen in other models of oxidative lung injury^66^.

In the vascular chip, the response to respiratory oxidative stress was most pronounced in the vascular endothelium. HPAECs showed activation of interferon- and cytokine-mediated signalling pathways (STAT1, IFI27, ISG15), extracellular matrix remodelling (TGFBR2, MMP14, POSTN, ENG, SERPINE1), gap junction regulation (GJA1), and apoptotic signalling (BAK1, IFI27). Notably, several proteins associated with endothelial-to-mesenchymal transition (TGFBR2, MMP14, THBS1, COL1A1, and POSTN), were also upregulated, consistent with the direction of changes observed in pulmonary vasculature in PH and COPD^67–71^. HPAEC proteome was impacted by co-culture with mural cells, which increased endothelial expression of proteins involved in cytoskeletal remodelling, adhesion, contractility and wound healing. This is consistent with data from genetic mouse models showing that pericytes and vascular smooth muscle cells can enhance endothelial reparative potential under disease conditions^59,72^. Vascular fibroblasts, particularly perivascular fibroblast populations, can also contribute to endothelial resilience and reparative capacity by supporting vascular stability, depositing extracellular matrix, and promoting neovascularization; emerging evidence further indicates that these cells can act as stromal progenitors and directly participate in vascular regeneration following injury^73,74^. Epithelial oxidative stress also had a significant impact on mural cells, in particular HPFs, which showed reduced expression of differentiation markers (*COL1A1*, *VIM*, *PDGFRα*), increased expression of immune (*VCAM1*, *CCL5*), apoptotic (*BAK1*, *CASP9*)^75^, angiogenic (*POSTN*) and proliferative (*BRD4*) markers, changes also noted in vascular remodelling ^76–78^ and lung fibrosis^79^.

Consistent with this activation state, the circulating medium in REVAS showed increased levels of pro-inflammatory and immunomodulatory cytokines and chemokines, involved in immune cell recruitment, activation, and tissue remodelling in COPD, including IL-6^80,81^, TNF-α/β^81,82^, IL-8^81,83^, CCL5^81^, CXCL9^81^, PDGF^84^, and TGF-β^81^. Identifying individual contributions of different cellular components of REVAS to this response will require further studies.

Comparative analysis of HPAEC datasets from the REVAS model with proteomic^40^ and transcriptomic^41,42^ endothelial COPD datasets revealed that approximately 32 % of REVAS DEPs showed an overlap with DEPs and DEGs dysregulated in COPD endothelium. This overlap highlights shared molecular signatures associated inflammation, cytoskeletal reorganisation, and vascular remodelling, encompassing key regulators such as STAT1^85^, TGFBR2^67,86^, POSTN^70^, ENG^87^, TPM1^88^, MYH9^89^, and VCL^90^.

At the same time, distinct differences emerged. For example, ICAM1 and SERPINE1 were upregulated in the REVAS database but downregulated in COPD datasets, whereas regulators of cell motility such as AHNAK and MACF1 showed the opposite pattern. This divergence likely reflects the contrast between acute endothelial activation captured in REVAS and the chronic adaptive changes characteristic of established disease, as well as the lack of a simple correlative relationship between proteomic and transcriptomic datasets^91^.

Taken together, these findings highlight the importance of multicellular *in vitro* models for studying vascular responses in lung injury and position REVAS as a promising translational platform for modelling respiratory-vascular interactions relevant to COPD. The system captures early pathogenic signalling events that may precede vascular dysfunction and provides a foundation for future studies of chronic injury, including those using patient-derived cells. Its modular plug-in/plug-out design also allows the respiratory and vascular compartments to be studied independently or in combination, enabling analysis of both distinct and coordinated responses to pollutants, infections, and other drivers of lung disease.

## MATERIALS AND METHODS

Detailed description of materials and methods can be found in Supplemental Methods.

### REVAS Design and Fabrication

Photomasks were designed using the Computer Aided Design (CAD) software, AutoCAD 2020 (AutoDesk Inc, Ca, USA). Photomasks for top and bottom parts of microfluidic chips were printed on high resolution photomask films (MicroLitho, UK). The number of cells grown in each channel was evaluated by dividing channel area by mean spreading area of different cell types obtained from confocal microscopy images analysed using ImageJ 2020 (NIH, University of Wisconsin, USA).

Chips were fabricated by photolithography and soft lithography techniques using polydimethylsiloxane (PDMS; Dow Corning, USA, Sylgard 184 Elastomer Kit; Cat. No. 634165S) poured onto pre-prepared patterns on silicon dioxide wafers (IDB Technologies, UK; W-Si Wafer 04/007), as described in ^12^.

### Flow simulation

Flow simulations were performed in COMSOL Multiphysics® version 6.1 (COMSOL AB, Sweden) using a laminar flow model to define operating conditions that yield physiologically <u>relevant</u> wall shear stress (WSS) within the REVAS platform. A detailed description of the model and PDMS wall model parameters are provided in the Supplementary Methods.

To confirm that the flow rates defined by in silico modelling were accurately delivered within the microfluidic system, flow rate measurements were performed using a Sensirion flow sensor (Liquid Flow Sensor, Cat. No. 403-SLF3S-1300F). The flow circuit was primed with distilled water, following the manufacturer’s instructions. After priming, the device was linked to the reservoirs through tubing and perfused with distilled water for 1 minute at a flow rate of 160 μl/min, corresponding to a WSS of 4 dynes/cm². Data was collected and analysed using the Sensirion Data Viewer software (Sensirion AG, Switzerland).

### Cell culture

All cells were maintained in T75 cell culture flasks (Sarstedt, Germany, Cat. No. 833911002) coated with 0.2 % porcine gelatine (Sigma, Cat. No. G1890) in a humidified incubator (37 °C, 5 % CO_2_). Cells were used between passages 4 – 8.

#### Human pulmonary artery endothelial cells

(HPAEC; PromoCell, Germany, Cat. No. C-12241) from 3 different biological donors were cultured in Endothelial Growth Medium 2 (EGM-2, PromoCell, Germany C-22211), <u>Human pulmonary artery smooth muscle cells</u> (HPASMC; Lonza, UK, Cat. No. CC-2581) from 3 different biological donors were cultured in Smooth Muscle Cell Growth Medium 2 (SmGM-2, PromoCell, Cat. No. C-22062); <u>Human pulmonary</u> <u>fibroblasts</u> (HPF; PromoCell, Cat. No. 12360) from 3 different biological donors were cultured in Fibroblast Growth Medium 2 (PromoCell, Cat. No. C-23020); <u>Human pericytes</u> (HPC; PromoCell, Cat. No. C-12980) from 3 different biological donors were cultured in Pericyte Growth Medium 2 (PromoCell, Cat. No. C-28041); <u>Human pulmonary microvascular</u> <u>endothelial cells</u> (HPMVEC; PromoCell, Cat. No. C-12281) from 3 different biological donors were cultured in in Endothelial Cell Growth Medium MV2 (PromoCell, Cat. No. C-22221); <u>Immortalised human pulmonary microvascular endothelial cells (HULEC-5a;</u> ATCC, Cat. No. CRL-3244) were cultured in MCDB131 medium (Merck, Cat. No. M8537) substituted with 2% FCS, L-Glutamine (0.01 mmol/mL; Merck, Cat. No. G5792), Hydrocortisone (Stemcell, Cat. No. 07926; 1 μg/mL), EGF (10 ng/mL). All media were supplemented with 1% Streptomycin/Penicillin (100 μg/mL) and 1 % MycoZap™.

#### Human small airway epithelial cells

(HsAEpC; Lonza, Cat. No. CC-2547) from 3 different biological donors were cultured in Small Airway Epithelial Cell Growth Medium (Lonza, Cat. No. CC-3118). The donor list is provided in **Supplementary Methods Table 2**.

After the assembly of respiratory and vascular chips, 20G stainless steel tubes (Coopers Needle Works, UK) were inserted into access ports of chips and secured with PDMS glue, which cured overnight at room temperature. Microfluidic channels were then coated with sterile filtered 0.2 % gelatine for 1 hour at 37 °C. For the vascular chip, macrovascular endothelial cells were seeded in top channels at the density of 10,000 cells/μl in 17 µl and incubated at 37 °C for 2 hours to promote cell attachment. Other cell types (HPASMCs, HPFs, HPCs) were seeded in bottom channels at the same density in 9 µl and incubated similarly. In the respiratory chips, HsAEpCs and HPMECs were seeded in top and bottom channels, respectively, at the same density of 10,000 cells/μl in 17 µl and left to adhere for 2 hours. Detailed protocols of cell seeding in REVAS are provided in Supplemental Methods.

For the cell co-culture in REVAS, an optimised medium based on basal EGM2 medium, previously used in pulmonary-artery-on-a-chip ^92^, was used. To prevent excessive proliferation of HPFs within the REVAS system, the FBS was reduced from 10 % to 5 % as suggested by Lopez-Martinez et al. ^93^. VEGF, heparin, ascorbic acid, and hydrocortisone were excluded from EGM2 growth supplement to avoid dysregulated proliferation of HPASMCs, HPFs and HPCs^94–97^.

After 2 hours incubation, all channels were gently flushed with co-culture medium, and access needles were connected to 0.8 mm ID flexible tubing (IBIDI, Germany, Cat. No. 10840). Co-culture medium was perfused at the wall shear stress rate of 4 dynes/cm² (vascular channel) and 1 dyne/cm² (respiratory channel) using a 4-channel peristaltic pump (Ismatec, Cole-Parmer, UK). Volumetric flow rate was calculated via COMSOL simulations.

### Air-liquid interface (ALI)

Cell culture at an air-liquid interface requires two sets of media: one for cell seeding and the initial submerged phase of cell culture and the other one for culture under air exposure.

For 500 mL of media for submerged culture, 250 mL of Airway Epithelial Growth Medium was mixed with 250 mL of DMEM and supplemented with HEPES (5.96 mg/mL), BSA (1.5 µg/mL), BPE (4 µl/mL), rhEGF (10 ng/mL), Hydrocortisone (0.5 µg/mL), Epinephrine (0.5 µg/mL), Transferrin (10 µg/mL), Retinoic Acid (0.1 ng/mL), 1 % Streptomycin/Penicillin (100 μg/mL) and 1 % MycoZap™.

For 500 mL of media for air-exposed culture, 450 mL of PneumaCult™-ALI Basal Medium was supplemented with 50 mL of PneumaCult™-ALI 10X Supplement, 5 vials oPneumaCult™-ALI Maintenance Supplement, Heparin (4 µg/mL), Hydrocortisone (0.48 µg/mL), 1 % Streptomycin/Penicillin (100 μg/mL) and 1 % MycoZap™.

To set up an air-liquid interface (ALI), 6.5 mm Transwell inserts with 0.4 µm pore size polyester membranes (Corning, USA, Cat. No. 3470) were coated with sterile 0.2 % gelatin for 1 hour. HsAEpCs (30,000 cells) were suspended in 200 µl of the pre-mixed airway epithelial growth medium and seeded onto the PET membrane in each insert. The basal compartment was filled with 500 µl of medium for submerged culture, and the medium was changed every two days until the cells reached confluence. Once confluence was achieved, the medium was removed from the apical compartment to expose the cells to air. Pre-mixed PneumaCult-ALI (air exposure) medium was added to the basal compartment only, and the medium was changed every two days to maintain the cells at the air-liquid interface until full differentiated.

### Immunocytochemistry

Cells were fixed in 4 % paraformaldehyde (Sigma Aldrich, UK, Cat. No. 1004968350), permeabilised in 0.1 % Triton-X-100 (Sigma Aldrich, UK, Cat. No. X100-5ML), washed thrice in PBS and incubated in 2 % BSA (Sigma Aldrich, UK, Cat. No. A9418), as described in ^9^. List of primary and secondary antibodies is shown in Supplement. Samples were mounted in VECTASHIELD® Antifade Mounting Medium with DAPI (Vector Laboratories, Cat No. H-1200). Images were taken under a STELLARIS 8 Confocal Microscope (Leica Microsystems).

### H&E staining

To perform H&E (Hematoxylin & Eosin) staining on HsAEpCs and HPMECs, cells growing on PET membranes on-chips or transwell plates were fixed in 4 % paraformaldehyde (PFA), washed, placed apical side up into a 24-well plate containing 4 % low melting point agarose (ThermoFisher, Cat. No. R0801) at 37 °C. The plate was then placed on ice until the agarose solidified. The agarose-embedded samples were then sectioned into 4 µm slices and stained with H&E by the Research Histology Facility at the South Kensington Campus. Images were captured using the Aperio Versa 8 (Leica Biosystems, Germany).

### Image analysis

All image analyses were carried out in ImageJ (FIJI, University of Wisconsin). Cell alignment relative to the direction of flow^12^.

### Permeability assay

Endothelial and epithelial barrier function was evaluated by measuring passage of fluorescent Dextran through cell monolayer growing on top of the porous PET membrane in Transwell filters or in microfluidic chips. In microfluidic chips, a 1 mg/mL solution of 40 kDa FITC-Dextran (Sigma Aldrich, Dorset, UK, Cat. No. FD40S) was introduced into the co-culture medium and perfused through the upper (endothelial) vascular channel at shear stress of 4 dynes/cm². After 1 hour, the medium from the bottom channel was collected by flushing the channel with 250 μl of medium, which was collected at the channel outlet. For permeability experiments involving thrombin, 1 U/mL thrombin (Sigma Aldrich, Dorset, UK, Cat. No. T7513) was added to the co-culture medium containing 1 mg/mL FITC-Dextran, and the experiment was conducted following the same protocol. Measurements of the amount of FITC-Dextran that passed through the membrane were performed using a GLOMAX spectrophotometer (Promega, USA) with excitation/emission wavelengths of 490/525 nm. The apparent permeability (Papp [cm/s]) was determined using the formula described by Maoz et al.^81^ as shown below.

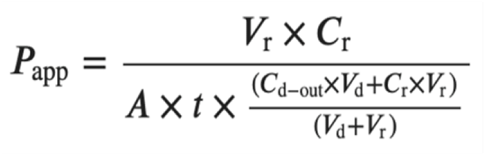

V_r_: Volume of receiving channel after time t

V_d_: Volume of the dosing channel after time t

A: area of membrane

C_r_: measured change in concentration of the tracer dye in the receiving channel

C_d_: measured concentration of the tracer dye in the dosing channel

t: 3600 seconds

### TaqMan qPCR

Gene expression analysis of cell differentiation markers in monocultures and co-cultures was conducted by qPCR. Cells were collected from PET membranes by trypsinisation, and RNA was extracted using the Monarch Total RNA Miniprep Kit (New England BioLabs, UK, Cat. No. T2010S) as per manufacturers’ instructions. cDNA was synthesised using the LunaScript RT SuperMix Kit (NEB, Cat. No. E3010S) on the SimpliAmp™ Thermal Cycler (ThermoFisher, Cat. No. A24811). The cDNA synthesis protocol included a primer annealing step at 25 °C for 2 minutes, followed by reverse transcription at 55 °C for 10 minutes, and heat inactivation at 95 °C for 1 minute.

qPCR was performed using a ViiA 7 Real-Time PCR System (ThermoFisher). Primer mixes were prepared by combining 12 µl of TaqMan primers (ThermoFisher) listed in Supplement, 12 µl of ddH₂O, and 120 µl of TaqMan Mastermix (ThermoFisher, Cat. No. 4305719). For each reaction, 3.6 µl of this primer mix was combined with 2.4 µl of cDNA, normalised to 1 ng/µl. The qPCR was run for a total of 45 cycles. The cycling conditions included a hold stage at 50 °C for 2 minutes, followed by 20 seconds at 95 °C. The PCR stage consisted of 1 second at 95 °C and 20 seconds at 60 °C for extension.

The relative gene expression was determined by the comparative ⊗CT method/ 2^-ÄÄCT^ method. Genes were normalised to Tyrosine 3-Monooxygenase/Tryptophan 5-Monooxygenase Activation Protein Zeta (*YHWAZ*) expression.

### Apoptosis Assay

To induce oxidative stress, HsAEpCs were treated with hydrogen peroxide (H₂O₂: Sigma-Aldrich, Cat. No.: H1009-500 mL) at concentrations ranging from 1 µM to 100 µM for 1, 2, and 4 hours to optimise conditions for studying oxidative stress-induced epithelial injury. Cells were seeded in 96-well plates (seeding density: 10,000 cells/well) and cultured for 48 hours. Caspase-Glo® 3/7 Reagent (Promega, Cat. No. G8090) was added directly to the wells, initiating cell lysis and generation of a luminescent signal through caspase-mediated substrate cleavage and luciferase activity. After a 1-hour incubation at room temperature, luminescence was measured using a GloMax® Luminometer (Promega). Data was normalised to untreated control.

### Optimisation of H_2_O_2_ treatment

To identify optimal effective concentration of H_2_O_2_ capable of inducing inflammatory response, HsAEpCs grown in 6-well plates (Thermo Fisher Scientific, Cat. No. 10578911) were exposed to H₂O₂ at concentrations of 10, 20, and 50 µM for 1 hour. 24 hours later, the medium collected from control or H_2_O_2_-stimulated HsAEpCs was transferred to HPAECs growing in a 6-well plate. After 48 hours incubation, cell pellets were collected from both HPAECs and HsAEpCs for qPCR analysis of apoptotic and inflammatory markers. TNF-α (R&D Systems, Cat. No. 210-TA) at 100 ng/ mL was used as a positive control.

### Experimental timeline for oxidative stress injury in REVAS

#### Day 1

Seeding submerged HsAEpCs in the top channel of respiratory chip.

#### Day 2

Seeding HMECs in the respiratory chip and seeding vascular chip (HPAECs, HPASMCs, HPFs, HPCs).

#### Day 3

Exposing HsAEpCs to 25 µM H_2_O_2_ (1 hour), then connecting respiratory and vascular chips for 48-hour perfusion in complete REVAS circuit.

#### Day 5

Harvesting cells for proteomic or qPCR analyses, collecting media for cytokine microarray.

### Protein extraction and mass spectrometry analysis

HPAECs from the two parallel top channels were collected by trypsinization and pooled together. A total of five independent REVAS circuits were used per condition (n=5). Cells were centrifuged at 1500 rpm and washed with PBS. Pellets were lysed with 25 µl of 8M urea lysis buffer on ice for 10 minutes. Following this, samples were sonicated in a water bath for 10 minutes, centrifuged at 10,000 xg for 10 minutes at 4 °C, and supernatants transferred to Protein LoBind® Tubes (Eppendorf, Cat. No. 0030122283). Mass spectrometry analysis was performed as outlined in ^12^. Protein concentrations were quantified using a Bradford assay. 20 µg of orotein was digested using an Andrews+ robot (Waters) in a single-pot digestion with 40 mM TCEP, 10 mM chloracetamide, 100 mM ammonium bicarbonate and 0.2 µg trypsin (Porcine Sequencing Grade, Promega), desalted with HLB μElution plates (Waters), and quantified by peptide assay. Peptides were analysed on a zenoTOF 7600 mass spectrometer (SCIEX) in positive mode using 85 variable windows spanning 400 to 900 mz zenoSWATH data independent acquisition (DIA). MS1 was acquired between 400 and 1500mz with 0.1s accumulation time. MSMS was acquired between 140 and 1800mz with an accumulation time of 0.013s. Interfaced with the MS, the peptides were separated across a 10 cm 150 µm internal diameter C18 column (Phenomonex) over a gradient spanning 3-35 % acetonitrile + 0.1 % (V/V) formic acid and 97-65 % water + 0.1 % (V/V) formic acid. The samples were randomised, and mass spectrometry data were processed in DIA-NN v1.8 using a library-free method using default settings. A predicated library was generated from a human canonical database (Swissprot) downloaded on the 23rd August 2022. Differential expression analysis of proteins that were quantified at least twice in both comparison groups was performed using the DEqMS V1.26 package in R^82^ on Log_2_ transformed data.

The mass spectrometry proteomics data have been deposited to the ProteomeXchange Consortium via the PRIDE^98^ partner repository with the dataset identifier PXD077544.

### Functional enrichment and protein network analysis

Enrichment analysis (GO Biological Processes^99^ and DisGeNET^30^) was performed using Metascape^100^. Differentially expressed proteins (DEPs) were selected for enrichment analysis based on the p-value or Benjimini-Hochberg^101^ adjusted p-value < 0.05 as indicated, a false discovery rate (FDR) < 0.05, and Log_2_ fold change of <-1.2 or >1.2.

### Cytokine profiling of REVAS culture medium

Cytokine profiling was performed on 100 µl of culture medium collected from the REVAS system after 48 hours of co-culture under basal or oxidative injury conditions (n=5 independent REVAS circuits/treatment group) using 48-plex Human Cytokine Assay platform (Eve Technologies, Canada). The fold change was calculated relative to the REVAS control group and plotted in bar-charts. For the heatmap, z-score of the concentration was plotted. Cytokines with concentrations below the limit of detection (OOR; out of range) were assigned a concentration of 0 pg/mL, according to the provider’s guidelines. Statistical comparisons were conducted using two-tailed unpaired t-tests.

### Comparative analysis of REVAS and COPD datasets

Endothelial gene expression data from age- and sex-matched endothelial cells of control and COPD lungs were kindly provided in pre-processed Excel format by Dr. Daria Kostyunina and Prof. Paul McLoughlin (University College Dublin)^40^.

Pulmonary endothelial transcriptomic data were derived from four publicly available scRNA-seq datasets^102–105^. Data were filtered to include only age-matched donors (50-76 years old) of both sexes (n males con/COPD, n females con/COPD). We focused on three main endothelial cell populations that participate in vascular remodelling: general capillary cells, aerocytes, and arterial endothelial cells. The differentially expressed genes for the endothelial cell subtypes were combined into one COPD endothelial dataset. Analysis was performed using the Seurat pipeline, with differential gene expression calculated using the Wilcoxon two-sample test and Bonferroni correction. Differentially expressed genes (DEGs) between control and COPD samples were selected based on adjusted P-value < 0.05 and a log₂ fold change > 0.6 (corresponding to >1.5-fold change) ^106^.

### Statistical analysis

Statistical analyses were performed using GraphPad Prism 9 software (GraphPad Software Inc., CA, USA). All experiments were conducted in at least three replicates, each with three technical replicates, unless otherwise specified. Data normality was assessed using the Shapiro-Wilk test. For comparisons between two groups, an unpaired Student’s t-test was applied to normally distributed data. For comparisons involving three or more groups, one-way or two-way ANOVA was used, as appropriate. Statistical significance was set at P < 0.05. Data are presented as mean ± SEM, with error bars indicating the standard error of the mean.

## Supporting information

Supplemental Figures, Tables and Methods Haensel et al.

## Data availability

Proteomics data generated and analysed during this study are available via ProteomeXchange with identifier PXD077544. All other data are available from the corresponding author (or other sources, as applicable) on reasonable request.

## Acknowledgements

The authors wish to thank Prof. Darryl Overby (Department of Bioengineering, Imperial College London) for his helpful advice on chip design; Dr Daria Kostyunina (NHLI, Imperial College London, London, UK) and Prof. Paul McLoughlin (School of Medicine, University College Dublin, Ireland) for providing access to COPD RNAseq databases; Prof. Lan Zhao (Imperial College London, London, UK) for providing HULEC-5a (ATCC® CRL-3244™); Dr Regis Joulia, Dr Laura Yates, Dr Helen Stoelting, Sara Patti and Dr Rafaela Konstantinidi (NHLI, Imperial College London, London, UK) for their help and advice in ALI culture, immunohistochemistry and cell imaging; Mr Steve Rothery (Imperial FILM facility) for his assistance and advice with image acquisition and analysis. All widefield/confocal imaging analysis was performed at the Facility for Imaging by Light Microscopy (FILM) at Imperial College London. We also thank Research Histology Facility at the South Kensington Campus, Imperial College London, London, UK) for assistance in histological analysis. Infrastructure support was provided by the National Institute for Health Research (NIHR) Imperial Biomedical Research Centre (BRC).

## Author contributions

MH designed and performed experiments, analysed data, and wrote the manuscript; RM performed experiments, and analysed proteomics data; HW preformed proteomic analyses, AA contributed to REVAS design and critically evaluated manuscript; DK analysed COPD proteomics data and critically evaluated manuscript; LB performed flow simulations; JvBS analysed flow pulsatility and critically evaluated the manuscript; CL provided guidance in experimental design, secured funding and critically evaluated the manuscript; BWS conceived the study, secured funding, performed experiments and wrote the manuscript.

