## Supplemental Figures, Tables and Methods Haensel et al. for "Organ on chip model of respiratory vascular interactions under COPD relevant oxidative stress"

### SUPPLEMENTAL RESULTS AND MATERIALS AND METHODS

Supplemental Results: Figures page 2- 18

Supplemental Results: Tables page 19-34

Supplemental Materials and Methods with Supplemental Methods Tables: page 35-57

Supplemental References: page 57-58

SUPPLEMENTAL FIGURES

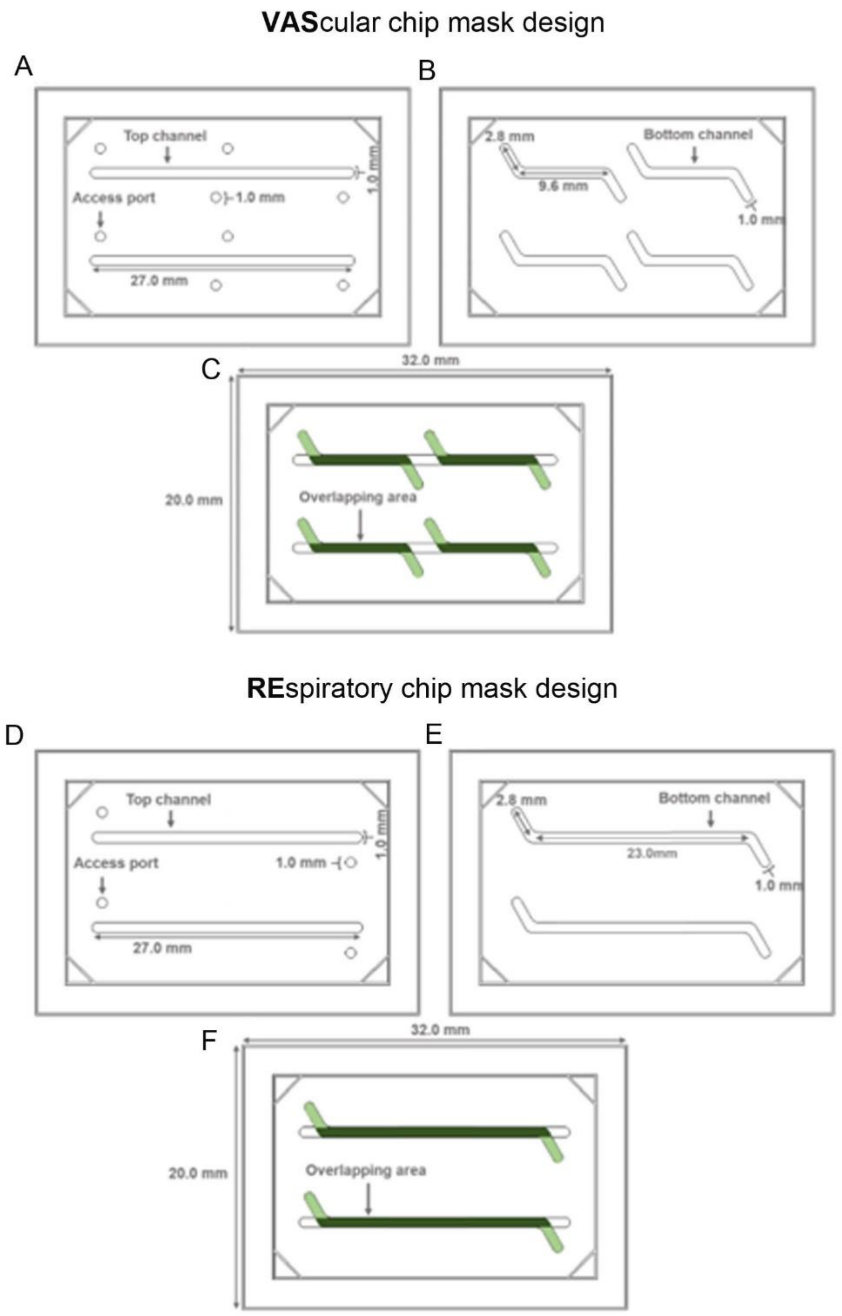

**Figure 1: Photomask design for vascular and respiratory chips in REVAS. (A)** Two parallel top channels were designed as linear, each measuring 27 mm in length and 1 mm in width, with guidelines for punching access ports to the bottom channels illustrated in **(B) (C)** Four bottom channels were designed in a transverse configuration, each measuring 15.2 x 1.0 mm. **(D)** In the respiratory chip, two straight parallel top channels each measured 27 mm in length and 1 mm in width, with guidelines for punching access ports to the bottom channel. **(E)** Two bottom channels, each measuring 28.6 x 1.0 mm. **(F)** The contact area between the top and bottom channels is shown in dark green, while the non-overlapping region is shown in light green. The designs were created using AutoCAD 2020 software.

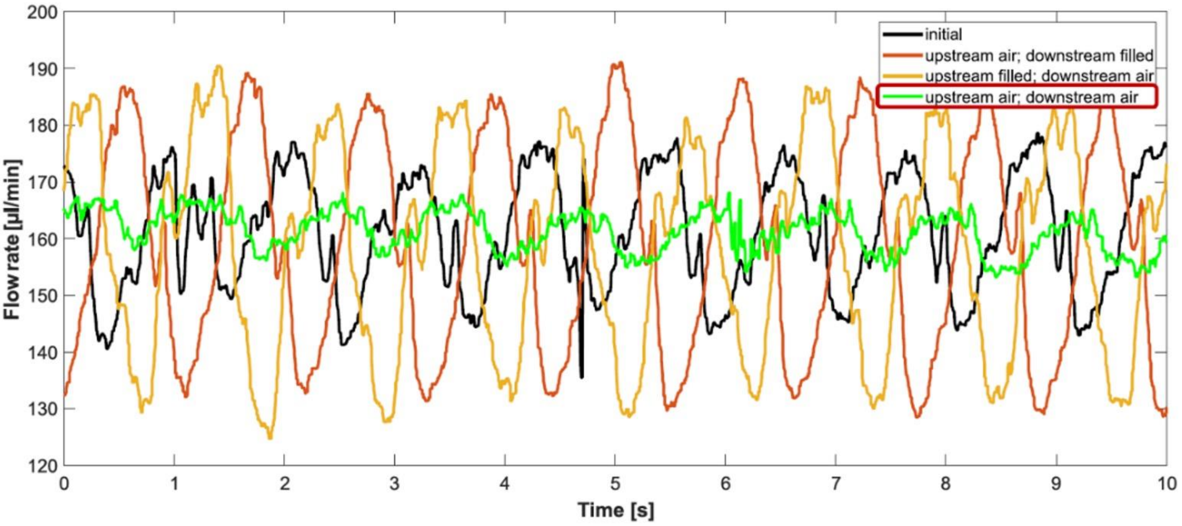

**Figure 2. Dampening of flow pulsatility by media reservoirs**. Flow rate (in µl/min) was measured over time (in seconds) under different upstream (inlet) and downstream (outlet) reservoir conditions. The black line represents both reservoirs fully filled with 2 mL medium (no air gap). The red and orange lines represent asymmetric conditions, where one reservoir contained 2 mL medium and the other 1.7 mL medium, leaving a 0.3 mL air gap. The green line represents both reservoirs containing 1.7 mL medium with a 0.3 mL air gap. Flow within the chip (4 dynes/cm$^2$ = 160 µL/min) was performed with a Sensirion Liquid Flow Sensor (# 403-SLF3S-0600F, Sensirion AG, Switzerland)

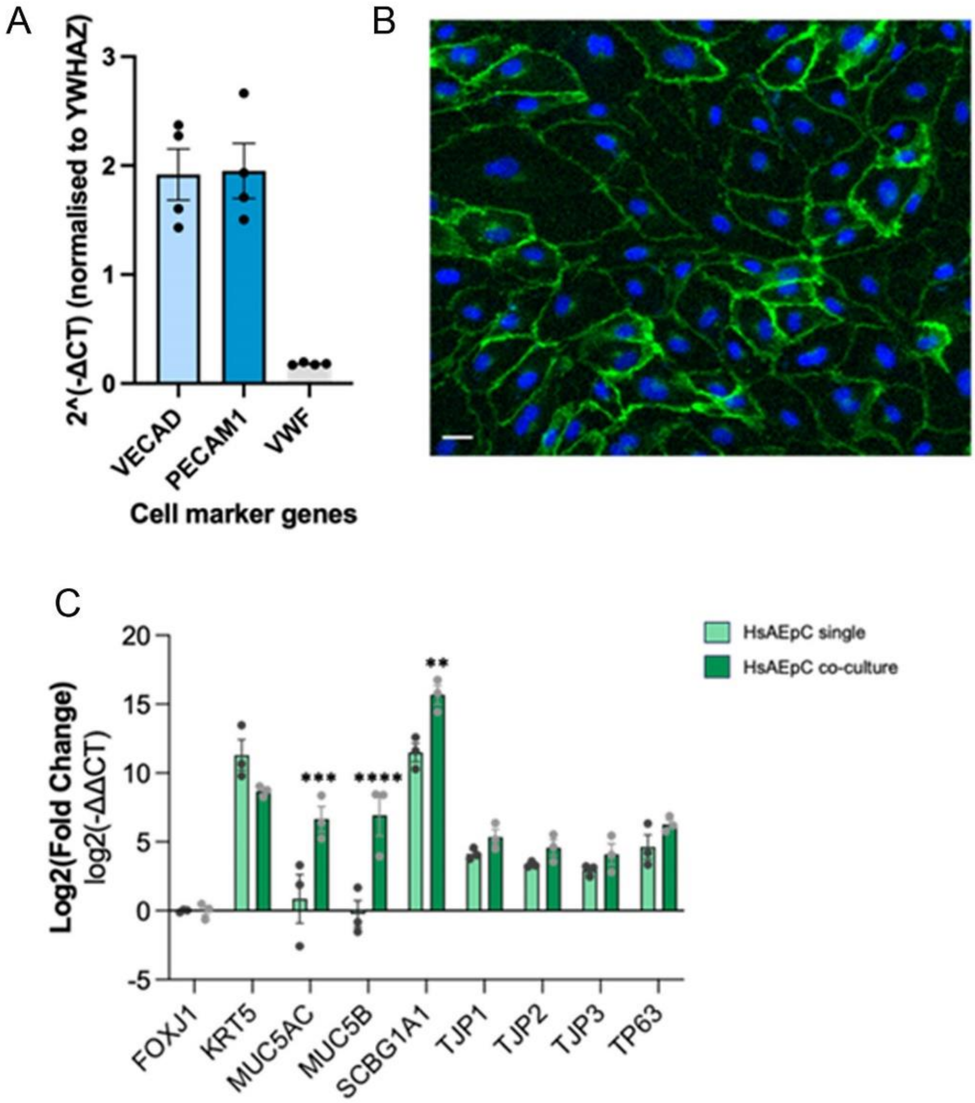

**Figure 3. Differentiation marker expression in cells cultured in the respiratory chip. (A)** Endothelial cell marker expression in human pulmonary microvascular endothelial cells (HPMVECs) cultured under flow (4 dynes/cm$^2$, 48 hrs). Gene expression was normalised to Tyrosine 3-Monooxygenase/Tryptophan 5-Monooxygenase Activation Protein Zeta (*YHWAZ*) expression. Error bars are means ± SEM. n=4. **(B)** Human microvascular endothelial cells cultured for 3 days; nuclei DAPI: blue, VE-Cadherin: green, immunofluorescence. scale bar = 20 µm. **(C)** Expression of airway epithelial cell differentiation markers in HsAEpCs cultured alone or in co-culture with HPMVECs. 30,000 HsAEpCs were seeded onto the PET membrane and cultured until confluent. The apical compartment was air-exposed, for 7 days. Forty-eight hours prior to completion, inserts were inverted and HsAEpCs were co-cultured with HPMVECs seeded on the basal side. Relative gene expression was determined by the comparative ΔCT method/ 2-ΔΔCT method. Genes were normalised to *YHWAZ* expression. Differences between groups were analysed with two-way ANOVA with Šidák's multiple comparisons test. Error bars indicate mean ± SEM. ** $P < 0.01$; *** $P < 0.001$; **** $P < 0.0001$, comparison between monoculture and respiratory chip co-culture; n=3.

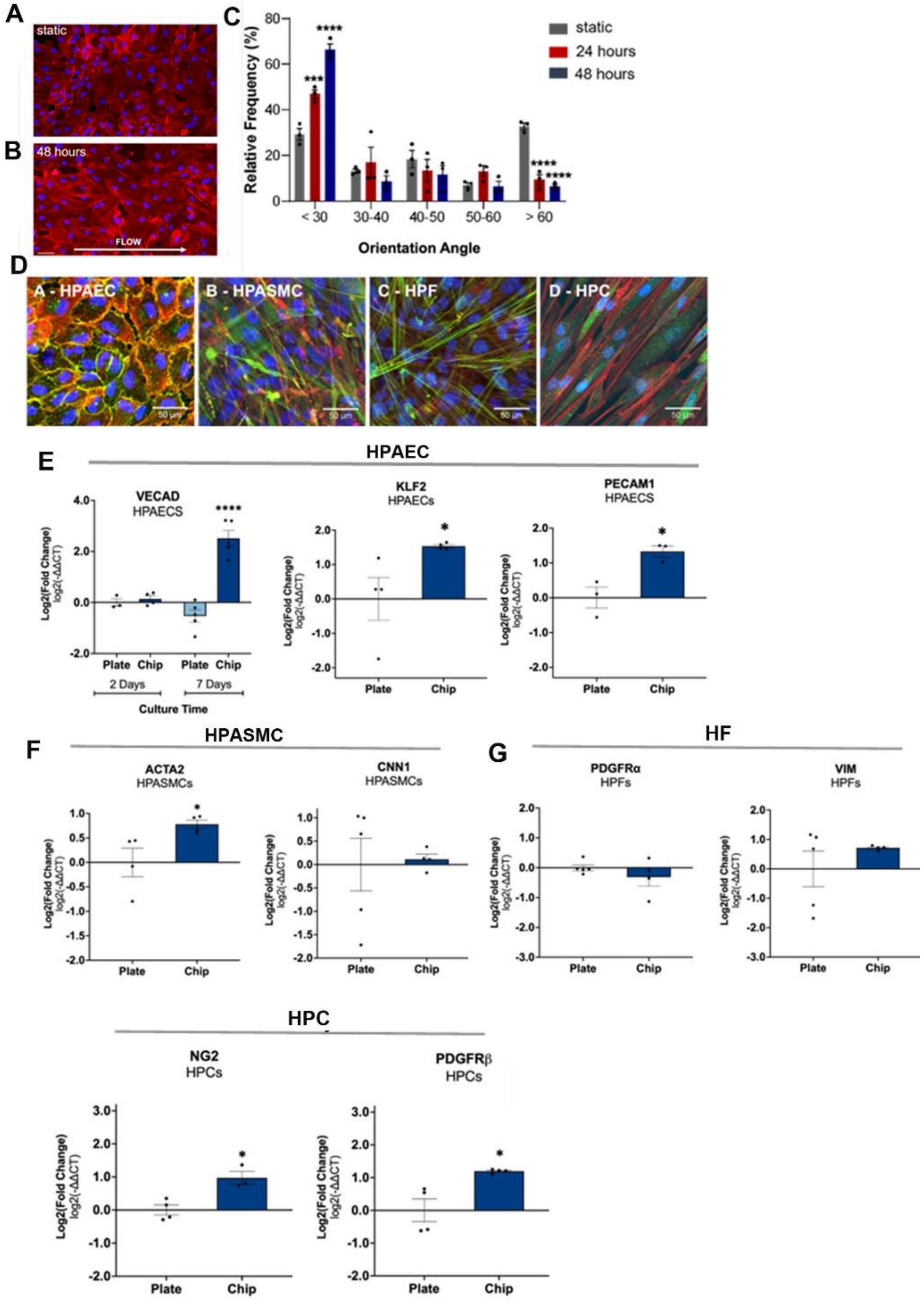

**Figure 4. Differentiation marker expression in cells cultured in the vascular chip. (A, B)** Representative images of HPAECs cultured under **(A)** static condition (top) or **(B)** under laminar flow conditions at 4 dynes/cm² for 48 hours (bottom) in the vascular chip. Scale bar: 20 µm; arrow: direction of flow; red: TRITC (F-Actin), blue: Dapi (nuclei). **(C)** HPAEC orientation changes under flow. The angle of orientation was measured between the cell's long axis (Feret's diameter) and the direction of

flow. Cells were considered aligned if the orientation angle was < 30° relative to the flow direction. Two-way ANOVA with Dunnett's multiple comparisons test. Error bars are means ± SEM. *P<0.05, *** P < 0.001; **** P < 0.0001, comparisons with respective static controls; n=3. **(D)** Images showing differentiation marker expression in HPAECs (VE-Cadherin; green), HPASMCs (calponin; green), HPFs (TAGLN; green) and pericytes (NG2; green), as indicated. Confocal microscopy, scale bar = 50 µm. **(E, F, G, H)** Differentiation marker gene expression in **(E )** HPAECs, **(F)** HPASMCs, **(G)** HPFs and **(H)** HPCs co-cultured in the vascular chip for 48 hours. Relative gene expression was determined by the comparative ΔCT/ $2^{-\Delta\Delta CT}$ method. Genes were normalised to Tyrosine 3-Monooxygenase/Tryptophan 5-Monooxygenase Activation Protein Zeta (*YHWAZ*) expression. * P < 0.05; unpaired t-test. Error bars indicate mean ± SEM. n≥3.

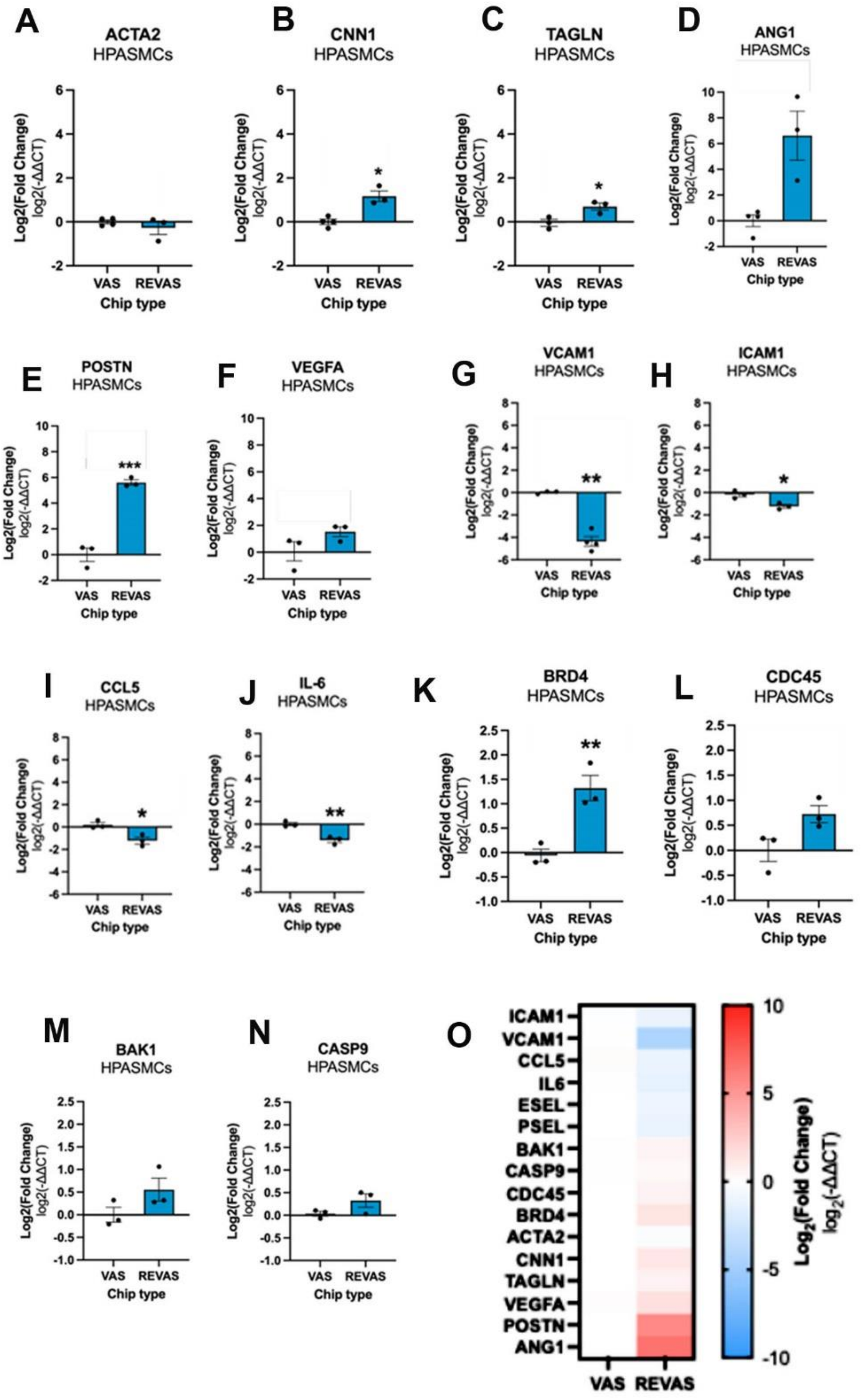

**Figure 5. Effect of REVAS co-culture on HPASMCs gene expression under basal conditions.** HPASMCs were cultured within the vascular chip (VAS) or within the complete REVAS circuit for 48h under basal conditions. **(A, B, C)** SMC differentiation marker gene expression (*ACTA2*, *CNN1*, *TAGLN*); (**D, E, F**) Angiogenic marker gene expression (*ANG1*, *POSTN*, *VEGFA*); (**G, H, I, J**) Inflammatory marker gene expression (*VCAM1*, *ICAM1*, *CCL5*, IL6); (**K, L**) Proliferation marker gene expression (*BRD4*, *CDC45*); (**M, N**) Apoptosis marker gene expression (*BAK1*, *CASP9*), as indicated. Relative gene expression of cell identity markers was assessed using the comparative $\Delta CT/2^{\wedge-\Delta\Delta CT}$ method and normalised to *YHWAZ* expression. Data are represented as mean ± SEM; n≥3. Statistical analysis was performed using an unpaired t-test.* $P < 0.05$, **$P < 0.01$, ***$P < 0.001$, comparison between cells cultured in VAS vs REVAS ; **(O)** Heatmap summarising gene expression changes shown in (A-N).

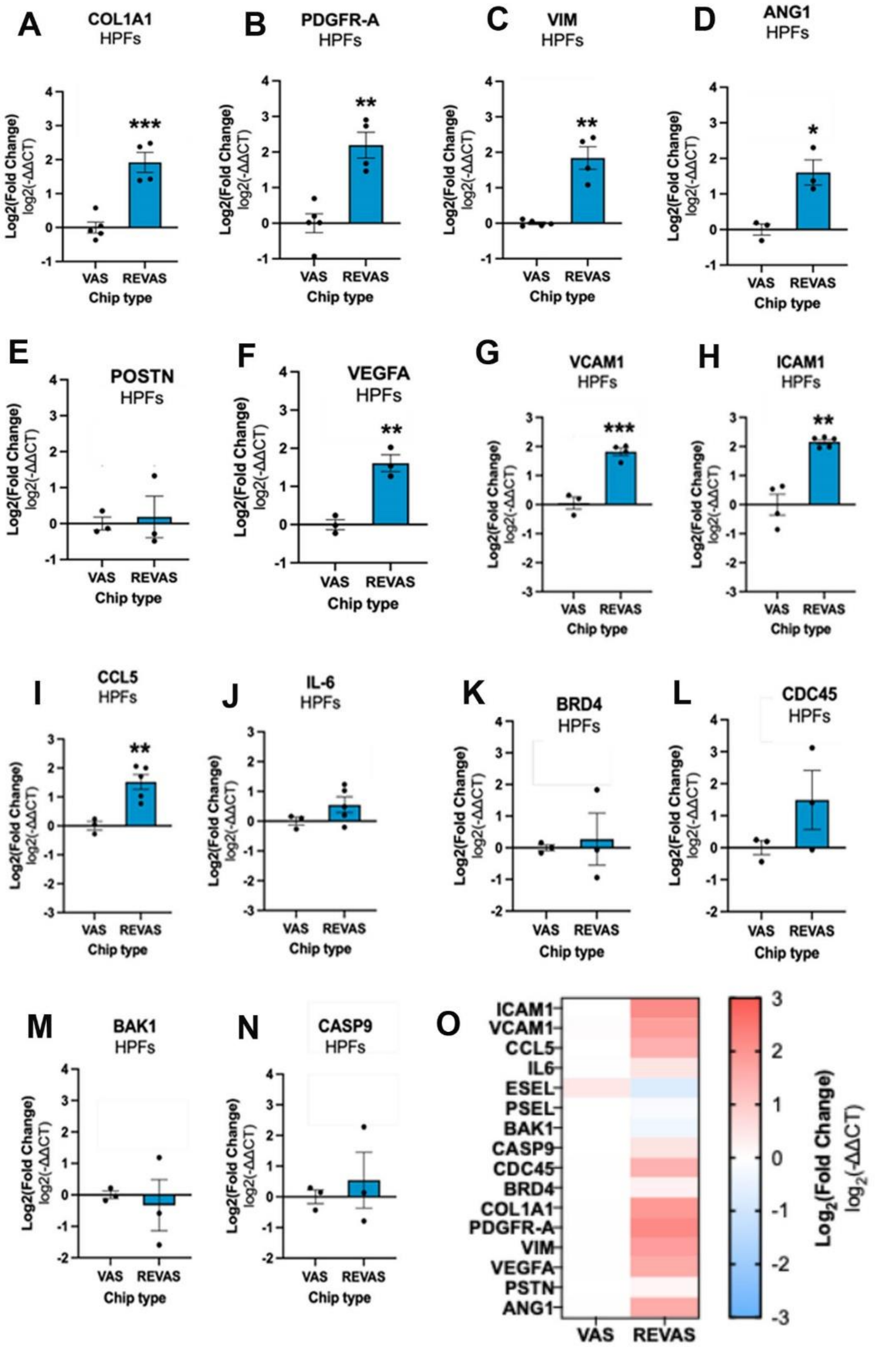

**Figure 6. Effect of REVAS co-culture on HPFs gene expression under basal conditions.** HPFs were cultured within the vascular chip (VAS) or within the complete REVAS circuit for 48h under basal conditions. **(A, B, C)** HFs differentiation marker gene expression (*COL1A1*, *PDGF-A*, *VIM*); (**D, E, F**) Angiogenic marker gene expression (*ANG1*, *POSTN*, *VEGFA*); (**G, H, I, J**) Inflammatory marker gene expression (*VCAM1*, *ICAM1*, *CCL5*, *IL6*); (**K, L**) Proliferation marker gene expression (*BRD4*, *CDC45*); (**M, N**) Apoptosis marker gene expression (*BAK1*, *CASP9*), as indicated. Relative gene expression of cell identity markers was assessed using the comparative $\Delta CT/2^{\wedge-\Delta\Delta CT}$ method and normalised to *YHWAZ* expression. Data are represented as mean ± SEM; n≥3. Statistical analysis was performed using an unpaired t-test; *P < 0.05, **P < 0.01, ***P < 0.001, comparison between cells cultured in VAS and REVAS. **(O)** Heatmap summarising gene expression changes shown in (A-N).

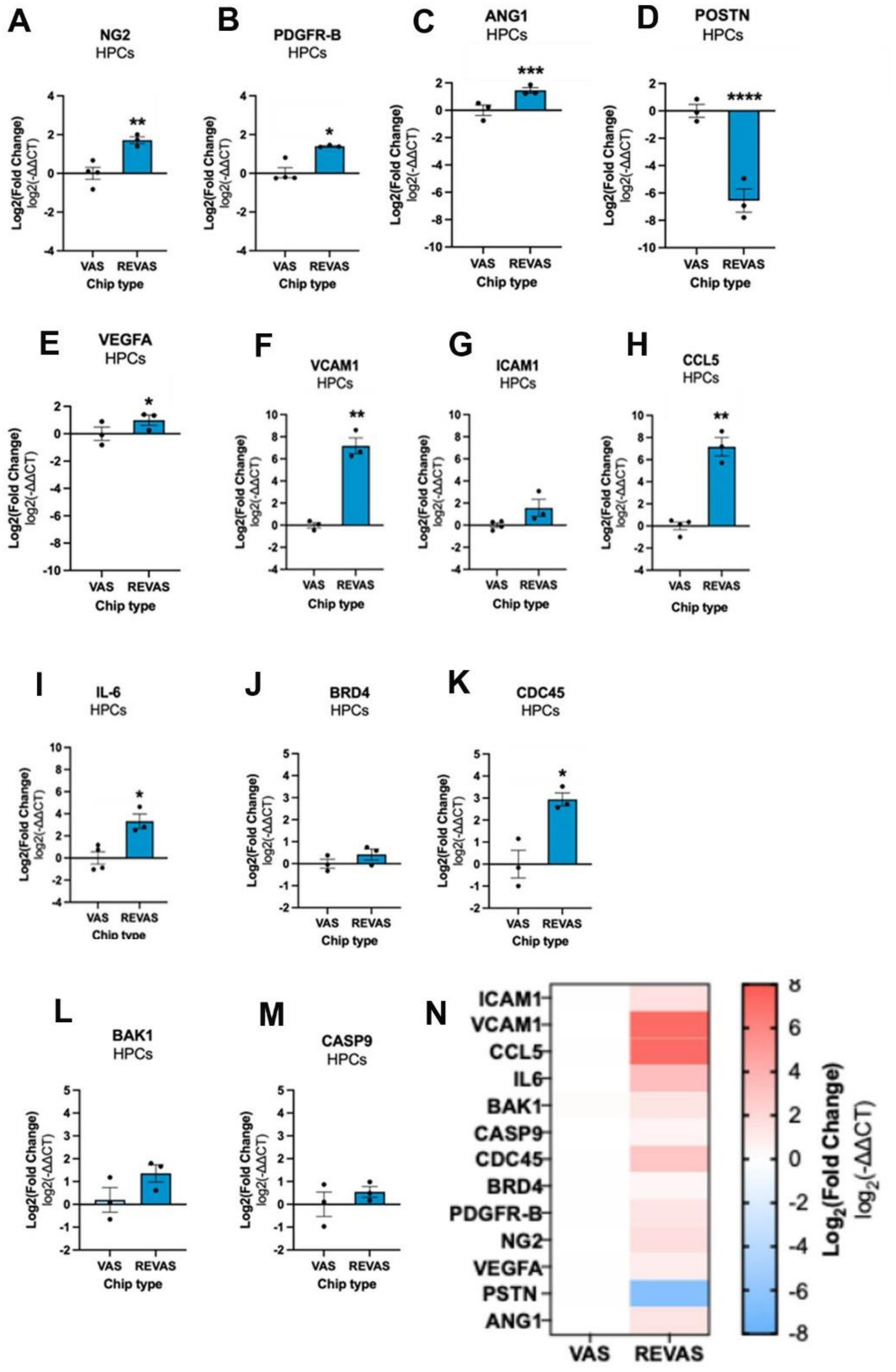

**Figure 7. Effect of REVAS co-culture on HPCs gene expression under basal conditions.** HPCs were cultured within the vascular chip (VAS) or within the complete REVAS circuit for 48h under basal conditions. **(A, B,)** HPCs differentiation marker gene expression (*NG2*, *PDGFR-B*); **(C, D, E)** Angiogenic marker gene expression (*ANG1*, *POSTN*, *VEGFA*); **(F, G, H, I)** Inflammatory marker gene expression (*VCAM1*, *ICAM1*, *CCL5*, *IL-6*); (**J, K**) Proliferation marker gene expression (*BRD4*, *CDC45*); (**L, M**) Apoptosis marker gene expression (*BAK1*, *CASP9*), as indicatedRelative gene expression of cell identity markers was assessed using the comparative $\Delta CT/2^{\wedge -\Delta\Delta CT}$ method and normalised to *YHWAZ* expression. Data are represented as mean ± SEM; n≥3. Statistical analysis was performed using an unpaired t-test; * $P < 0.05$, ** $P < 0.01$, *** $P < 0.001$, **** $P < 0.0001$, comparison between cells cultured in VAS and REVAS. **(N)** Heatmap summarising gene expression changes shown in (A-N).

A

B

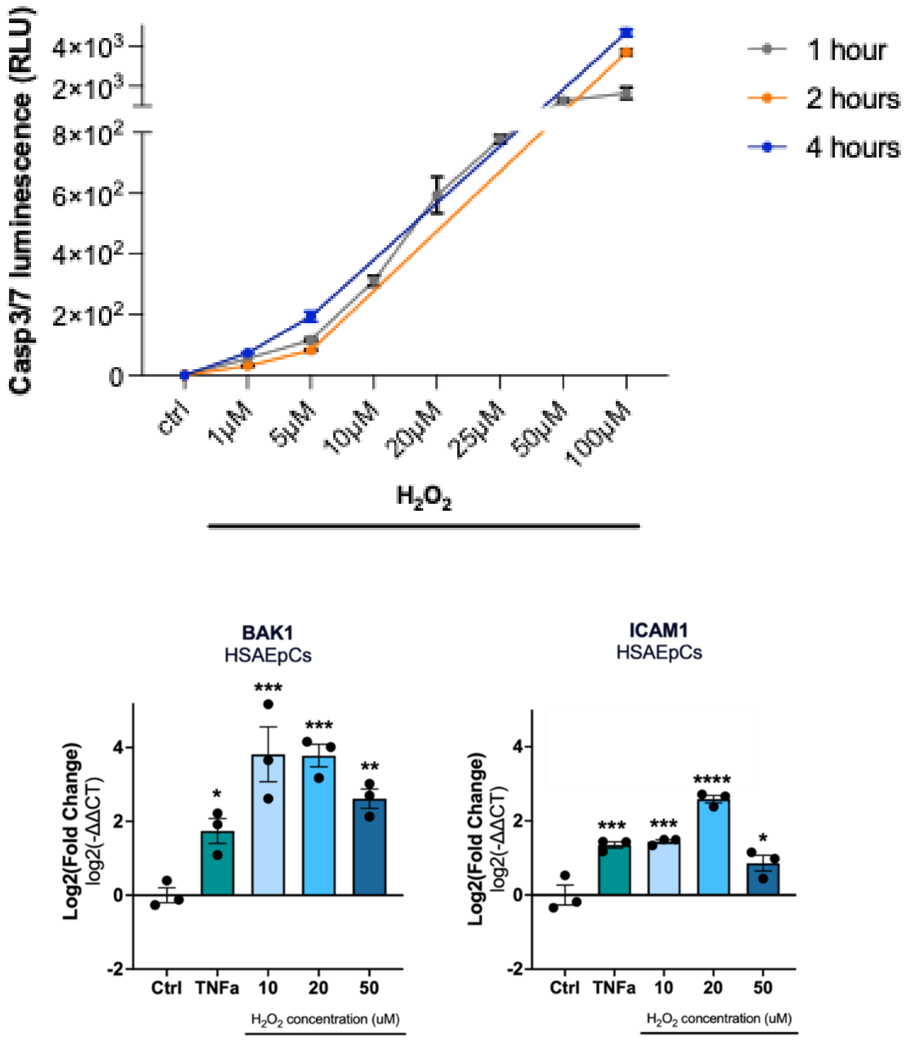

**Figure 8. Apoptotic and inflammatory activation of human small airway epithelial cells (HsAEpC) treated with $H_2O_2$. (A)** Apoptotic response measured in caspase 3/7 assay. HsAEpCs were treated with 1, 5, 10, 20, 25, 50 and 100 µM $H_2O_2$ for 1, 2 and 4 hours to induce oxidative stress. A corresponding number of cells was left untreated. The Caspase-Glo® 3/7 Reagent was added directly to the cells in 96-well plates. This led to cell lysis, followed by caspase cleavage of the substrate, then generation of a "glow-type" luminescent signal, produced by luciferase. After 1 hour incubation at room temperature, the luminescence was read with a Glomax Luminometer and data was normalised to control. Error bars indicate mean ± SEM, n=3. (**B**) Apoptotic and inflammatory marker expression in HsAEpCs treated with $H_2O_2$ in the static REVAS setup**.** HsAEpCs were cultured in 6-well plates and treated with $H_2O_2$ (10, 20, or 50 µM) for 24 hours. A concentration of 100 ng/mL TNFα was used as a positive control. After 24 hours, cells were harvested and analysed by qPCR for expression of apoptotic and inflammatory markers. Relative gene expression of cell identity markers was determined using the comparative $\Delta CT/2^{\wedge-\Delta\Delta CT}$ method and normalised to *YHWAZ* expression. Error bars indicate mean ± SEM. Statistical significance was assessed using a one-way ANOVA with a Šidák multiple comparisons test; *P < 0.05, **P < 0.01, ***P < 0.001, ****P < 0.0001; n=3.

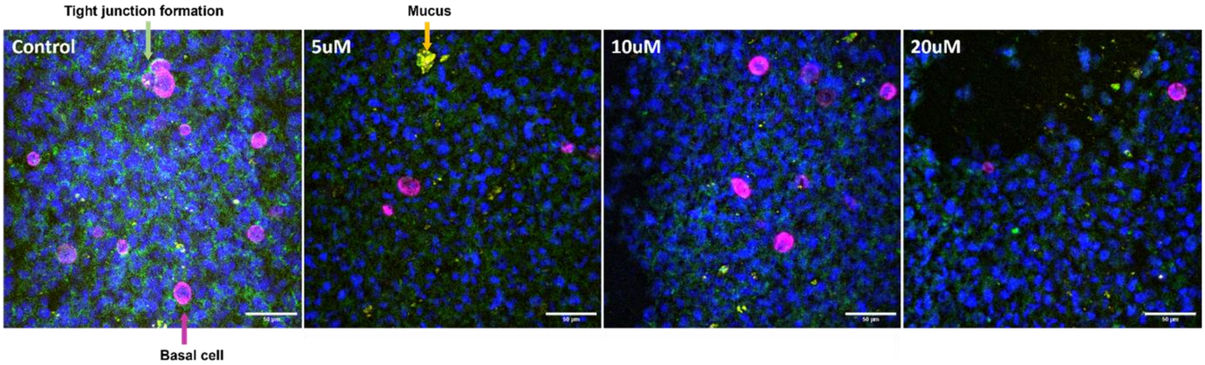

**Figure 9. $H_2O_2$ treatment of air-exposed HsAEpCs.** HsAEpCs were cultured in the top channel of the respiratory chip under a continuous medium flow of 100 µl/hr for three days. Following this, the cells were air-exposed and maintained under air flow for an additional eight days. $H_2O_2$ was then introduced into the top channel at concentrations of 5, 10, and 20 µM and incubated for 1 hour before fixation with 4 % PFA and staining. Scale bar = 50 µm.
●Dapi: nuclei ●KRT5: basal cells● ZO-1: tight junctions ●MUC5AC: goblet cells

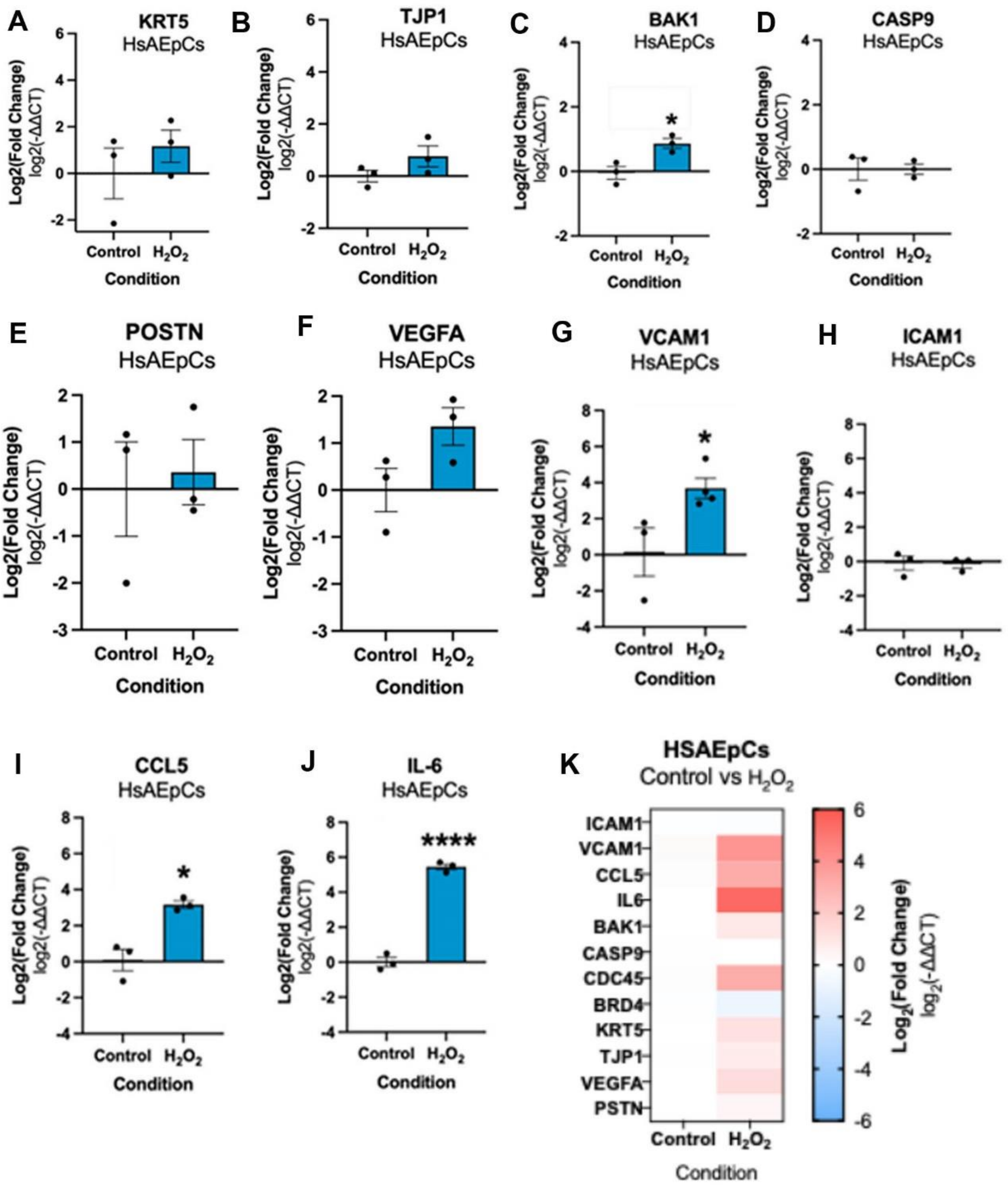

**Figure 10. $H_2O_2$-induced gene expression changes expression in HsAEpCs.** HsAEpCs were co-cultured in the respiratory chip with HMECs. 1h following $H_2O_2$-treatment of HsAEpCs, the respiratory and vascular chips were connected and 48 hours later marker gene expression in HsAEpCs was measured by qPCR. (**A, B**) Expression of epithelial cell identity markers *KRT5* and *TJP1*; **(C, D)** Expression of apoptosis markers *BAK1* and *CASP9*; **(E, F)** Expression of angiogenesis markers *POSTN* and *VEGFA*; **(G-J)** Expression of inflammatory marker genes *VCAM1*, *ICAM1*, *CCL5* and *IL-6* in HsAEpCs cultured in the respiratory chip under basal and $H_2O_2$-treated conditions. Relative gene expression was assessed using the comparative $\Delta CT/2^{-\Delta\Delta CT}$ method and normalised to *YHWAZ* expression. Data are presented as mean ± SEM; $n \geq 3$. *$P < 0.05$; ****$P < 0.0001$, unpaired t-test, comparison with untreated controls. **(K)** Heatmap summarising changes in gene expression shown in (A-J).

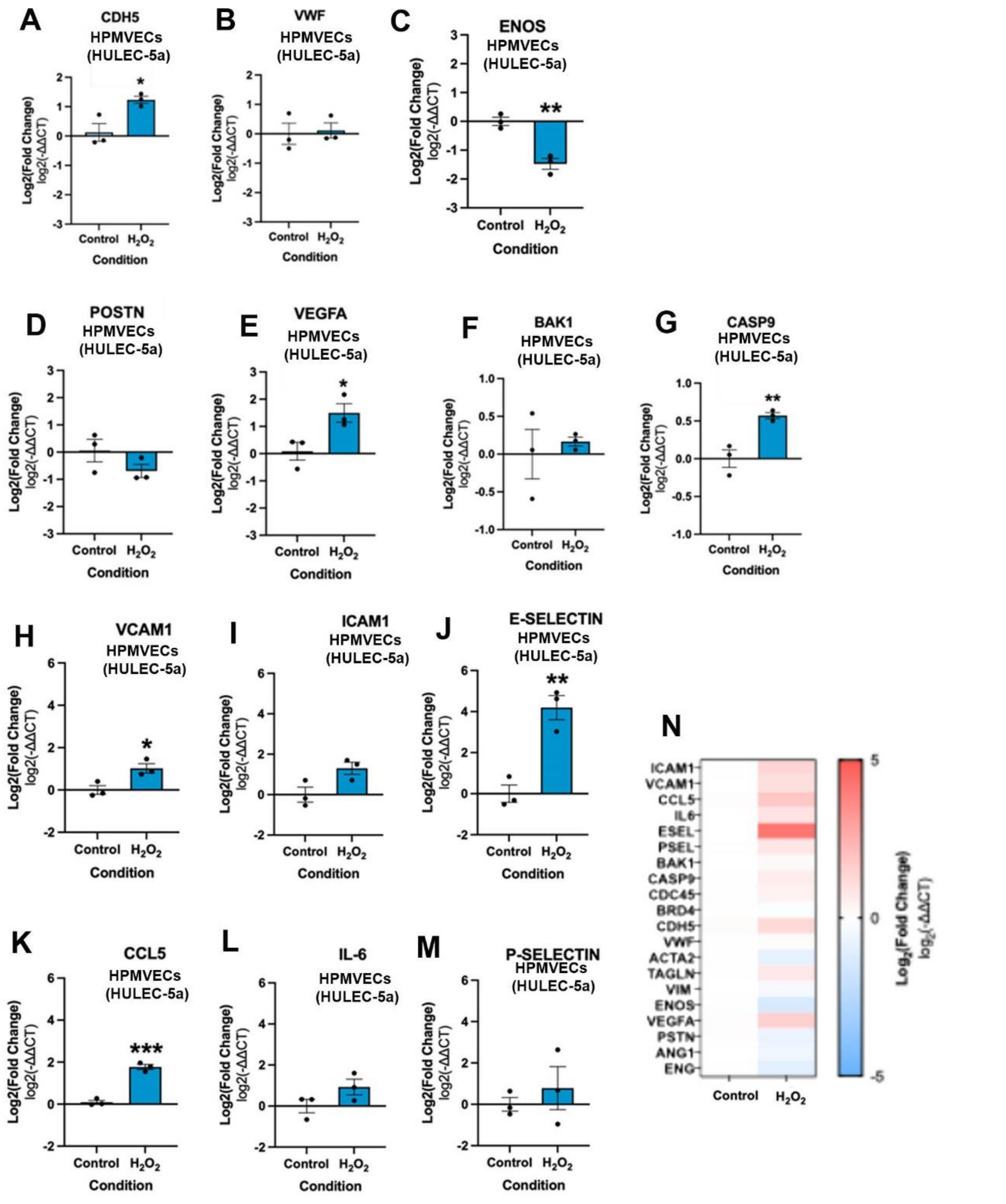

**Figure 11: $H_2O_2$-induced gene expression changes expression in HPMVEC cell line (HULEC-5a).** HPMVECs (HULEC-5a) were co-cultured with HsAEpCs in the respiratory chip. 1h following $H_2O_2$-treatment of HsAEpCs, the respiratory and vascular chips were connected and 48 hours later marker gene expression in HPMVECs was measured by qPCR. (**A, B, C**) Expression of epithelial cell identity markers *CDH5*, *VWF* and *NOS3*; **(D, E)** Expression of angiogenesis markers *POSTN* and *VEGFA*; **(F, G)** Expression of apoptosis markers *BAK1* and *CASP9*; **(H-M)** Expression of inflammatory marker genes *VCAM1*, *ICAM1*, *ESEL*, *CCL5*, *IL-6* and *PSEL* in HMVCs cultured in the respiratory chip under basal and $H_2O_2$-treated conditions. Relative gene expression was assessed using the comparative $\Delta CT/2^{\wedge-\Delta\Delta CT}$ method and normalised to *YHWAZ* expression. Data are presented as mean ± SEM; $n \geq 3$.

*P < 0.05 **P < 0.01, unpaired t-test, comparison with untreated controls. (**N**) heatmap summarising gene expression changes shown in (A-M).

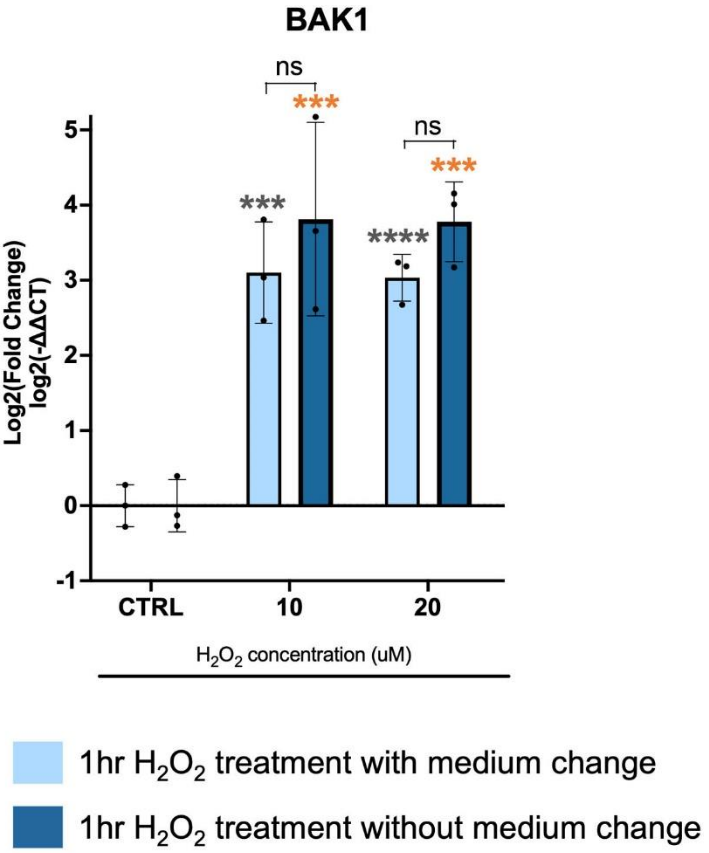

**Figure 12. BAK1 expression in HsAEpCs following HsAEpCs $H_2O_2$ exposure with or without media change.** HsAEpCs cultured in the upper chambers of 6-well Transwell plates were treated with $H_2O_2$ (10 or 20 µM) for 1 hour. They were then co-cultured with HPAECs in the lower chambers either without a media change (dark blue bars) or after washing the HsAEpCs with fresh medium prior to co-culture (light blue bars). Expression of apoptotic marker (*BAK1*) was measured by qPCR using the comparative $\Delta CT/2^{\wedge -\Delta\Delta CT}$ method and normalised to *YHWAZ*. Error bars indicate mean ± SEM. Statistical analysis was performed using a one-way ANOVA with a Šidák multiple comparisons test; ***P < 0.001, ****P < 0.0001, comparison between control and treatment groups; ns-not significant, comparisons, as indicated; n=3.

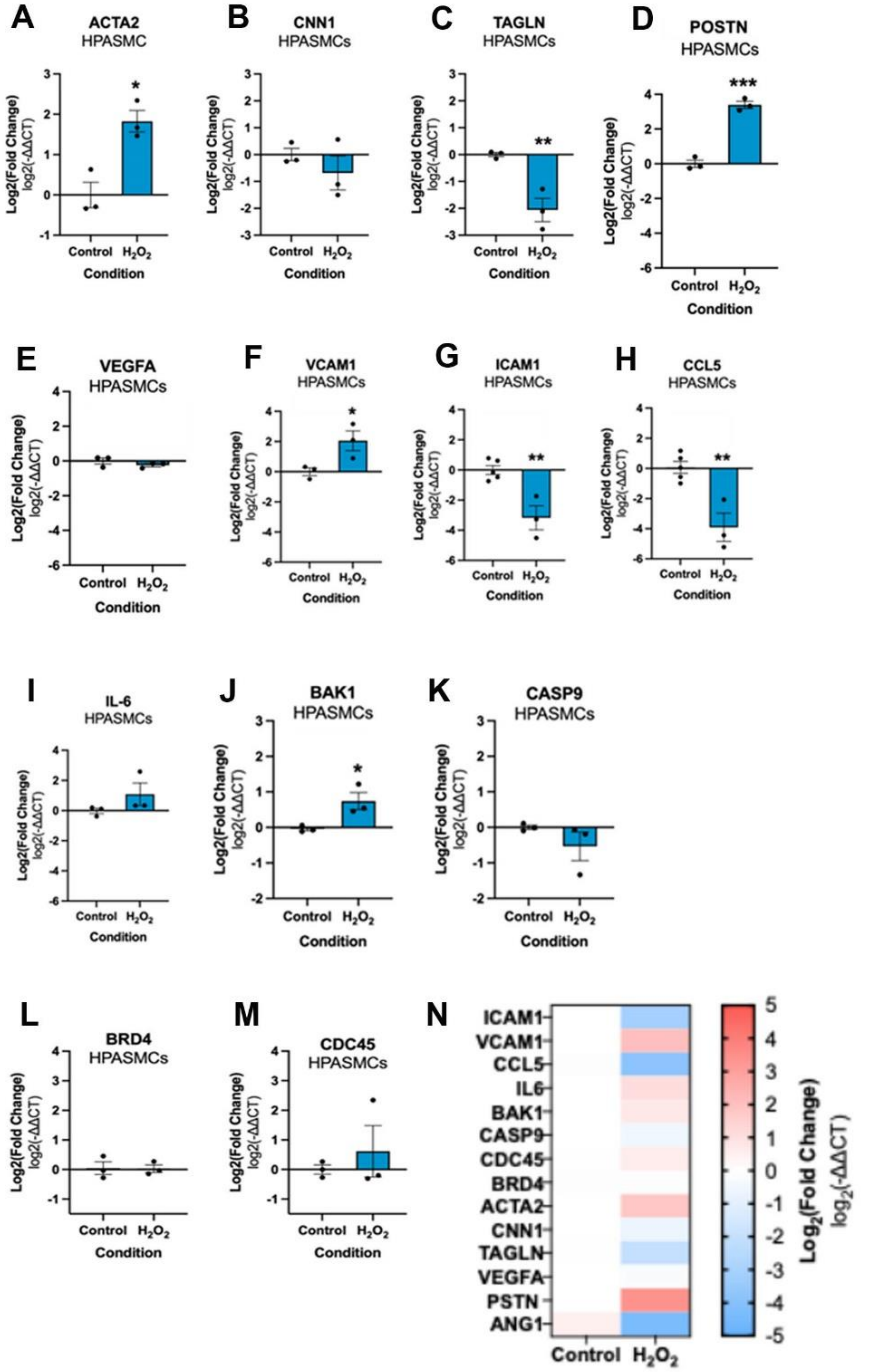

**Figure 13. Changes in the expression of marker genes in HPASMCs cultured in the REVAS circuit under $H_2O_2$-treated conditions** HPASMCs were cultured in the bottom channel of the vascular chip connected with respiratory chip. HsAEpCs were exposed to 25 µM $H_2O_2$ for 1 hour prior to integration of vascular and respiratory chips under flow. **(A, B, C)** SMC differentiation marker gene expression (***ACTA2***, ***CNN1***, ***TAGLN***); **(D, E)** Angiogenic marker gene expression (***POSTN***, ***VEGFA***); **(F-I)** Inflammatory marker gene expression (***VCAM1***, ***ICAM1***, ***CCL5***, ***IL-6***); **(J, K)** Apoptosis marker gene expression (***BAK1***, ***CASP9***); **(L, M)** Proliferation marker gene expression (***BRD4***, ***CDC45***), as indicated. Relative gene expression of marker genes was assessed using the comparative $\Delta CT/2^{-\Delta\Delta CT}$ method and normalised to ***YHWAZ*** expression. Data are represented as mean ± SEM; n=3. Statistical analysis was performed using an unpaired t-test; * $P < 0.05$, ** $P < 0.01$, *** $P < 0.001$, comparison between cells cultured in in REVAS with and without $H_2O_2$. **(N)** Heatmap summarising changes in gene expression shown in (A-M).

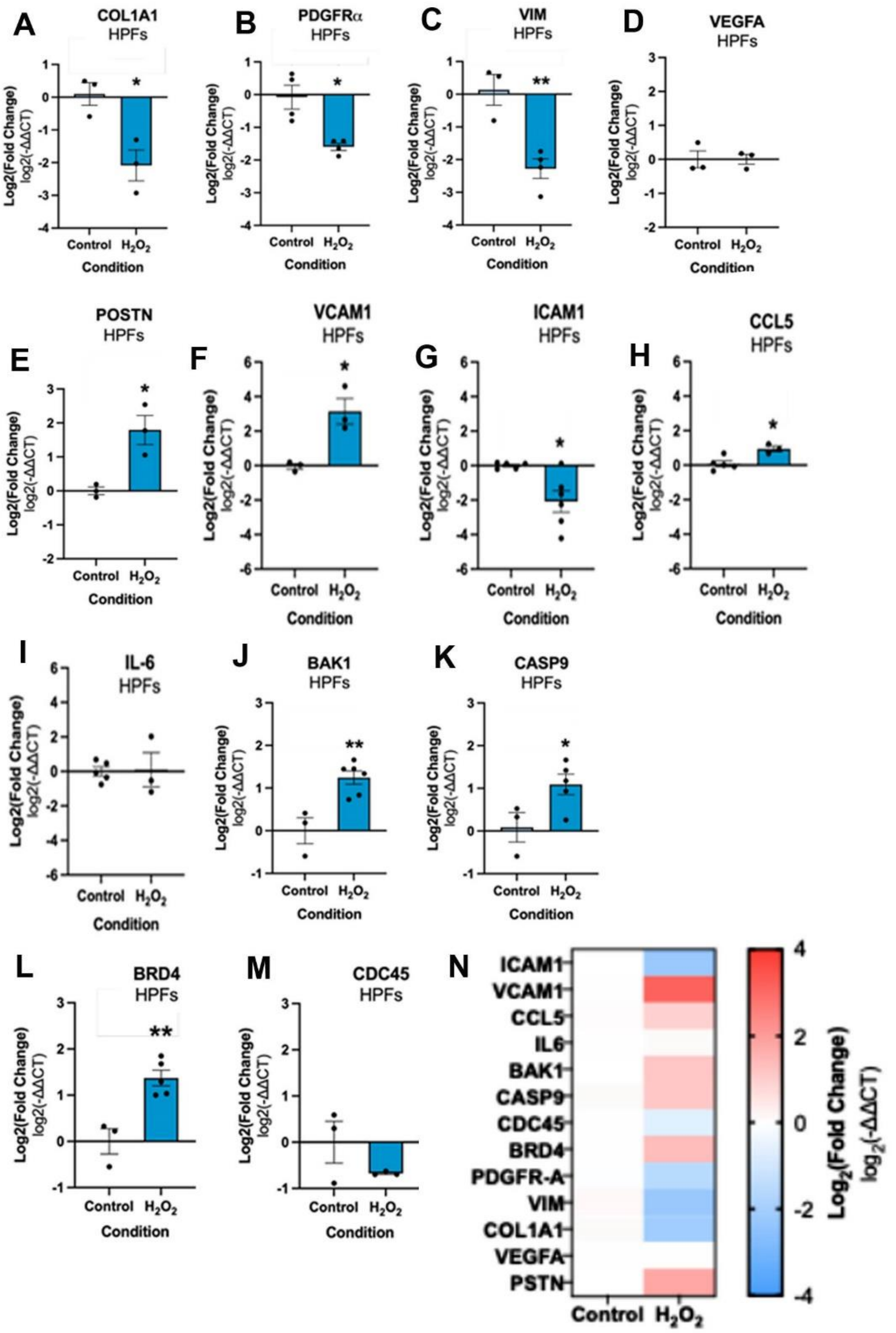

**Figure 14: Cell identity marker gene expression in HPFs cultured in the REVAS circuit under $H_2O_2$-induced oxidative stress.**HPFs were cultured in the bottom channel of the vascular chip connected with respiratory chip. HsAEpCs were exposed to 25 µM $H_2O_2$ for 1 hour prior to integration of vascular and respiratory chips under flow. (**A-C**) HFs differentiation marker gene expression (***COL1A1***, ***PDGF-A***, ***VIM***); (**D, E)** Angiogenic marker gene expression (***POSTN***, ***VEGFA***); **(F-I)** Inflammatory marker gene expression (***VCAM1***, ***ICAM1***, ***CCL5***, ***IL-6***); **(J, K**) Apoptosis marker gene expression (***BAK1***, ***CASP9***); (**L, M)** Proliferation marker gene expression (***BRD4***, ***CDC45***), as indicated. Relative gene expression of gene markers was assessed using the comparative ΔCT/2^−ΔΔCT method and normalised to ***YHWAZ*** expression. Data are represented as mean ± SEM; n=3. Statistical analysis was performed using an unpaired t-test; *P < 0.05, **P < 0.01, comparison between cells cultured in REVAS with, or without $H_2O_{2.2}$. **(N)** Heatmap summarising changes in gene expression shown in (A-M).

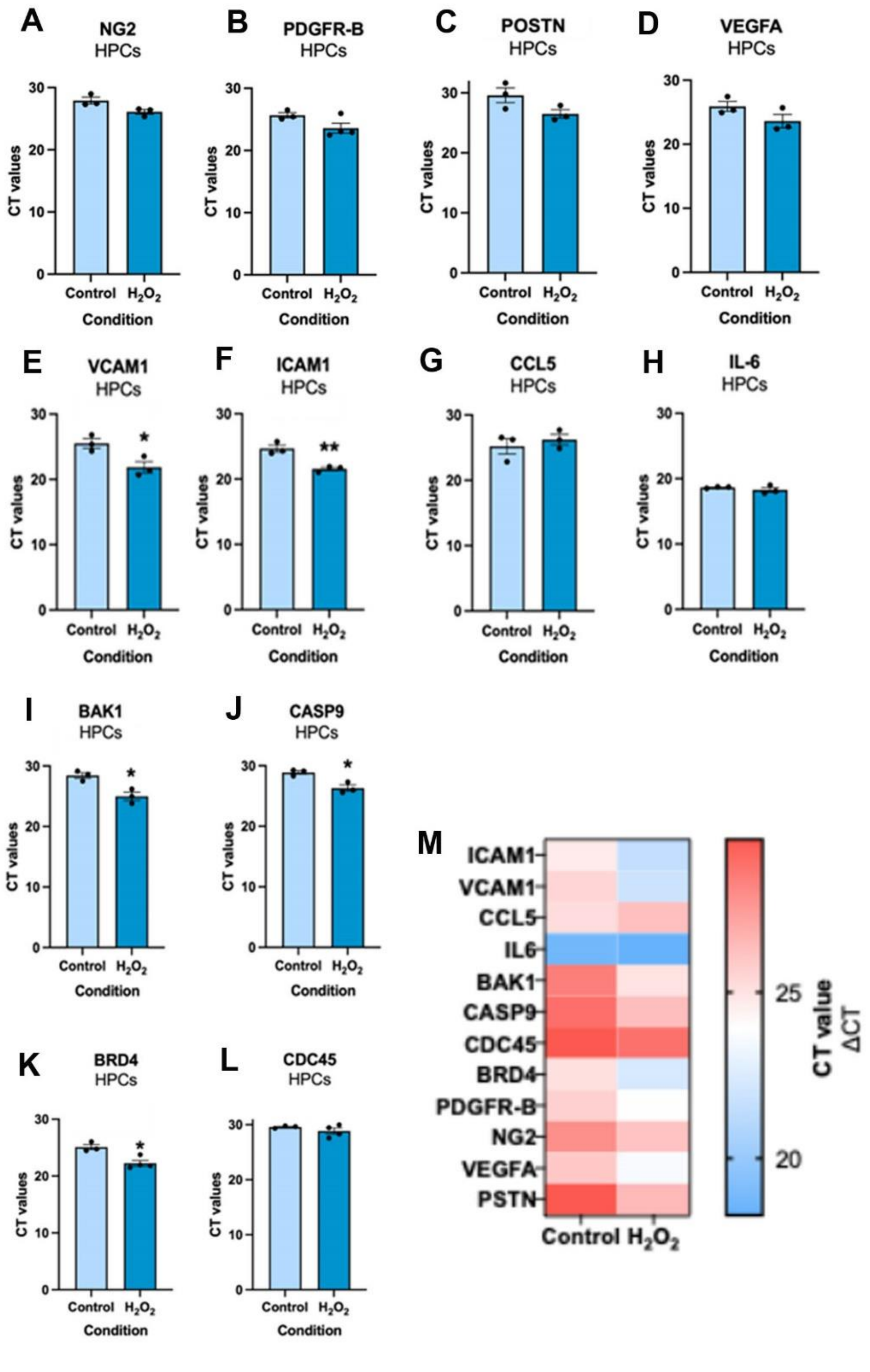

**Figure 15. Changes in marker gene expression in HPCs cultured in the REVAS circuit under $H_2O_2$-treated conditions.** HPCs were cultured in the bottom channel of the vascular chip connected with respiratory chip. HsAEpCs were exposed to 25 µM $H_2O_2$ for 1 hour prior to integration of vascular and respiratory chips under flow. **(A, B)** HPCs differentiation marker gene expression (*NG2*, *PDGFR-B*); (**C, D**) Angiogenic marker gene expression (*POSTN*, *VEGFA*); (**E-H**) Inflammatory marker gene expression (*VCAM1*, *ICAM1*, *CCL5*, *IL-6*); (**I, J**) Proliferation marker gene expression (*BRD4*, *CDC45*); **(K, L)** Apoptosis marker gene expression (*BAK1*, *CASP9*), as indicated. Due to variability in the housekeeping gene expression in HPCs, raw CT values were used to assess relative gene expression of proliferative markers. Data are represented as mean ± SEM; n≥3. Statistical analysis was performed using an unpaired t-test; $*P < 0.05$, comparison between cells cultured in REVAS with, or without $H_2O_2$. **(M)** Heatmap summarising changes in gene expression shown in (A-L).

### SUPPLEMENTARY RESULTS TABLES

**Supplementary Results Table 1: Dimensions of a single top and bottom channel in the vascular chip.**

| | Top channel | Bottom channel |
|---|---|---|
| **Length** | 27,000 μm | 15,200 μm |
| **Width** | 1,000 μm | 1,000 μm |
| **Height** | 200 μm | 200 μm |
| **Membrane Surface Area** | 27.0 $mm^2$ | 15.20 $mm^2$ |
| **Volume** | 5.4 μl | 3.04 μl |

**Supplementary Results Table 2: Dimensions of an overlapping area between the channels in the vascular chip.**

| Dimensions | |
|---|---|
| **Length** | 9,600 μm |
| **Width** | 1,000 μm |
| **Height** | 200 μm |
| **Membrane Surface Area** | 9.6 $mm^2$ |

**Supplementary Results Table 3: Size of HPAECs, HPASMCs, HPFs, HPCs.** Sizes were taken from microscopy images.

| | | Min [μm] | Max [μm] | Average [μm] | No. cells in channel | No. cells in overlap |
|---|---|---|---|---|---|---|
| **HPAEC** | **Length** | 60 | 90 | 75 | 9,474 | 3,368 |
| | **Width** | 25 | 50 | 38 | | |
| | **Area/cell [$\mu m^2$]** | | | 2,850 | | |
| **HPASMC** | **Length** | 120 | 240 | 180 | 2,815 | 1,778 |
| | **Width** | 15 | 45 | 30 | | |
| | **Area/cell [$\mu m^2$]** | | | 5,400 | | |
| **HPF** | **Length** | 140 | 130 | 135 | 5,362 | 3,386 |
| | **Width** | 15 | 27 | 21 | | |
| | **Area/cell [$\mu m^2$]** | | | 2,835 | | |
| **HPC** | **Length** | 110 | 70 | 90 | 8,042 | 5,079 |

| | | | | |
|---|---|---|---|---|
| | Width | 25 | 17 | 21 |
| | Area/cell [µm²] | | | 1,890 |

**Supplementary Results Table 4: Dimensions of a single top and bottom channel in the respiratory chip.**

| | Top channel | Bottom channel |
|---|---|---|
| **Length** | 27,000 µm | 28,600 µm |
| **Width** | 1,000 µm | 1,000 µm |
| **Height** | 200 µm | 200 µm |
| **Membrane Surface Area** | 27.0 $mm^2$ | 28.60 $mm^2$ |
| **Volume** | 5.4 µl | 5.72 µl |

**Supplementary Results Table 5: Dimensions of an overlapping area between the channels in the respiratory chip.**

| | Dimensions |
|---|---|
| **Length** | 2,300 µm |
| **Width** | 1,000 µm |
| **Height** | 200 µm |
| **Membrane Surface Area** | 23.0 $mm^2$ |

**Supplementary Results Table 6: Size of a HsAEpC and HPMVEC.** Sizes were taken from microscopy images.

| | | Min [µm] | Max [µm] | Average [µm] | No. cells in channel | No. cells in overlap |
|---|---|---|---|---|---|---|
| **HsAEpC** | **Length** | 35 | 52 | 43.5 | 20,350 | 17,336 |
| | **Width** | 27 | 34 | 30.5 | | |
| | **Area/cell [µm²]** | | | 1,327 | | |
| **HPMVEC** | **Length** | 52 | 76 | 64 | 13,965 | 11,230 |
| | **Width** | 26 | 38 | 32 | | |
| | **Area/cell [µm²]** | | | 2,048 | | |

**Supplementary Results Table 7: Top 15 GO pathway enrichment pathways with lists of linked upregulated DEPs in HPAECs cultured with vascular mural cells versus HPAECs cultured alone in a vascular chip.**

| Description | Hits |
|---|---|
| regulation of cellular catabolic process | SLC25A5\|APP\|ATP6V1C1\|PIK3C2A\|PSAP\|SNX4\|SNX5\|ATP6V1D\|RPTOR |
| regulation of autophagy | SLC25A5\|ATP6V1C1\|PIK3C2A\|PSAP\|SNX4\|SNX5\|ATP6V1D\|RPTOR |
| macrophage activation | APP\|IFNGR1\|NMI\|IL33 |
| localization within membrane | NACA\|PSAP\|SPTBN1\|SNX4\|ASAP1\|VPS37B\|FRMD8\|SH3PXD2B\|TOMM5 |
| stress granule assembly | FMR1\|ATXN2L\|YTHDF2 |
| post-Golgi vesicle-mediated transport | SPTBN1\|SCAMP1\|ASAP1\|CORO7 |
| positive regulation of catabolic process | SLC25A5\|APP\|FMR1\|PIK3C2A\|SNX4\|YTHDF2\|RPTOR\|IL33 |
| positive regulation of cellular catabolic process | SLC25A5\|APP\|PIK3C2A\|SNX4\|RPTOR |
| regulation of mRNA processing | FMR1\|SAFB\|SAFB2\|HDAC7 |
| establishment of protein localization to membrane | NACA\|SPTBN1\|ASAP1\|VPS37B\|TOMM5 |
| neuron cellular homeostasis | APP\|ATP6V1C1\|ATP6V1D |
| proton transmembrane transport | SLC25A5\|ATP6V1C1\|ATP6V1D\|SLC25A22 |
| positive regulation of inflammatory response | APP\|SNX4\|NMI\|IL33 |
| extracellular matrix organization | APP\|LOX\|MYO1E\|TGFBI\|SH3PXD2B |
| extracellular structure organization | APP\|LOX\|MYO1E\|TGFBI\|SH3PXD2B |

**Supplementary Results Table 8: Top 15 GO pathway enrichment pathways with lists of linked downregulated DEPs in HPAECs cultured with vascular mural cells versus HPAECs cultured alone in a vascular chip.**

| Description | Hits |
|---|---|
| membrane organization | SLC25A4\|SLC25A5\|SLC25A6\|ANXA2\|RHOA\|CAV1\|CAV2\|CDC42\|HK2\|HSPA8\|LETM1\|OXA1L\|PPT1\|PRKCA\|RAC1\|SURF4\|VAMP2\|VDAC2\|DEGS1\|SNAP23\|VAMP3\|HUWE1\|TOMM40\|AFG3L2\|TMED2\|EMC1\|EPB41L3\|EXOC7\|VPS8\|MTCH1\|MTCH2\|SAMM50\|MYO |

| | |
|---|---|
| | F\|GHITM\|SEC61A1\|EHD2\|NDUFA13\|EMC4\|RAB14\|RAB4B\|EXOC2\|NCLN\|TOMM22\|PLSCR3\|CHMP1B\|TMEM43\|RHOT2\|TOMM5 |
| generation of precursor metabolites and energy | ACADM\|ALDOA\|SLC25A4\|COX5B\|COX7A2\|CS\|CYC1\|DLAT\|FDXR\|FECH\|NIPSNAP2\|GBE1\|HK1\|HK2\|IDH2\|COX2\|NDUFA7\|NDUFB10\|NDUFS1\|NDUFS8\|OXA1L\|UQCRC1\|UQCRC2\|UQCRFS1\|UQCRH\|SLC25A12\|SUCLA2\|ATP5MF\|SLC25A13\|ATP5PD\|NDUFA13\|NDUFS7 |
| cellular respiration | COX5B\|COX7A2\|CS\|CYC1\|DLAT\|NIPSNAP2\|IDH2\|COX2\|NDUFA7\|NDUFB10\|NDUFS1\|NDUFS8\|OXA1L\|UQCRC1\|UQCRC2\|UQCRFS1\|UQCRH\|SLC25A12\|SUCLA2\|ATP5MF\|SLC25A13\|ATP5PD\|NDUFA13\|NDUFS7 |
| aerobic respiration | COX5B\|COX7A2\|CS\|CYC1\|DLAT\|NIPSNAP2\|IDH2\|COX2\|NDUFA7\|NDUFB10\|NDUFS1\|NDUFS8\|OXA1L\|UQCRC1\|UQCRC2\|UQCRFS1\|UQCRH\|SUCLA2\|ATP5MF\|ATP5PD\|NDUFA13\|NDUFS7 |
| energy derivation by oxidation of organic compounds | ACADM\|COX5B\|COX7A2\|CS\|CYC1\|DLAT\|NIPSNAP2\|GBE1\|IDH2\|COX2\|NDUFA7\|NDUFB10\|NDUFS1\|NDUFS8\|OXA1L\|UQCRC1\|UQCRC2\|UQCRFS1\|UQCRH\|SLC25A12\|SUCLA2\|ATP5MF\|SLC25A13\|ATP5PD\|NDUFA13\|NDUFS7 |
| mitochondrial transport | SLC25A4\|SLC25A5\|SLC25A6\|HK2\|LETM1\|SLC25A3\|VDAC1\|VDAC2\|SLC25A12\|SLC25A13\|TOMM40\|AFG3L2\|MTCH1\|MTCH2\|SAMM50\|GHITM\|SLC25A24\|NDUFA13\|TOMM22\|RHOT2\|TOMM5 |
| localization within membrane | ARF6\|CAV1\|GOLGA4\|HSPA5\|NACA\|OXA1L\|RAC1\|RALA\|RAP1A\|VAMP2\|VAMP3\|NHERF2\|TM9SF4\|TOMM40\|RAB10\|TMED2\|RAB31\|EMC1\|EPB41L3\|MTCH1\|MTCH2\|SAMM50\|SEC61A1\|EHD4\|EHD2\|NDUFA13\|EMC4\|RAB14\|EXOC2\|TM9SF3\|NCLN\|TOMM22\|TOMM5 |
| mitochondrion organization | SLC25A4\|SLC25A5\|SLC25A6\|CAV2\|COX7A2\|NIPSNAP2\|HK2\|LETM1\|NDUFB10\|NDUFS1\|NDUFS8\|NOS3\|OXA1L\|TFAM\|UQCRFS1\|VDAC2\|LONP1\|TOMM40\|AFG3L2\|MTCH1\|MTCH2\|SAMM50\|GHITM\|ACAD9\|NDUFA13\|TOMM22\|RHOT2\|NDUFS7\|TOMM5 |
| mitochondrial membrane organization | SLC25A4\|SLC25A5\|SLC25A6\|HK2\|LETM1\|OXA1L\|VDAC2\|TOMM40\|AFG3L2\|MTCH1\|MTCH2\|SAMM50\|GHITM\|NDUFA13\|TOMM22\|RHOT2\|TOMM5 |
| oxidative phosphorylation | COX5B\|COX7A2\|CYC1\|NIPSNAP2\|COX2\|NDUFA7\|NDUFB10\|NDUFS1\|NDUFS8\|UQCRC1\|UQCRC2\|UQCRFS1\|UQCRH\|ATP5MF\|ATP5PD\|NDUFA13\|NDUFS7 |
| protein localization to membrane | ARF6\|CAV1\|GOLGA4\|HSPA5\|NACA\|OXA1L\|RAP1A\|VAMP2\|VAMP3\|NHERF2\|TM9SF4\|TOMM40\|RAB10\|TMED2\|RAB31\|EMC1\|EPB41L3\|MTCH1\|MTCH2\|SAMM50\|SEC61A1\|EHD4\|EHD2\|NDUFA13\|EMC4\|TM9SF3\|NCLN\|TOMM22\|TOMM5 |
| proton transmembrane transport | SLC25A4\|SLC25A5\|ATP1A1\|COX5B\|CYC1\|LETM1\|COX2\|SLC25A3\|UQCRC1\|UQCRFS1\|UQCRH\|SLC25A12\|ATP5MF\|SLC25A13\|TCIRG1\|ATP5PD\|GHITM\|NDUFS7 |
| respiratory electron transport chain | COX5B\|COX7A2\|CYC1\|COX2\|NDUFA7\|NDUFB10\|NDUFS1\|NDUFS8\|UQCRC1\|UQCRC2\|UQCRFS1\|UQCRH\|SLC25A12\|SLC25A13\|NDUFS7 |
| purine nucleotide metabolic process | ADK\|ALDOA\|DLAT\|FDXR\|GMPR\|GNAI3\|HK1\|HK2\|HSPA8\|IDH2\|NDUFA7\|NDUFB10\|NDUFS1\|NDUFS8\|PCK2\|PPT1\|SUCLA2\|ATP5MF\|SLC25A13\|ATP5PD\|DNPH1\|NDUFA13\|HSD17B12\|GMPR2\|FAR1\|NDUFS7 |

| | |
|---|---|
| electron transport chain | COX5B\|COX7A2\|CYC1\|COX2\|NDUFA7\|NDUFB10\|NDUFS1\|NDUFS8\|UQCRC1\|UQCRC2\|UQCRFS1\|UQCRH\|SLC25A12\|SLC25A13\|NDUFS7 |

**Supplementary Results Table 9: Top 25 GO enriched pathways associated with upregulated DEPs in HPAECs, induced by connection of the vascular with the respiratory chip under basal conditions.**

| Description | Hits |
|---|---|
| cellular respiration | ACO2\|ATP5F1A\|ATP5F1B\|ATP5F1C\|ATP5PB\|ATP5ME\|ATP5PO\|COX4I1\|COX5B\|COX7A2\|COX7C\|CS\|CYC1\|DLAT\|DLD\|DLST\|ETFA\|ETFB\|FH\|NIPSNAP2\|GPD2\|IDH2\|IDH3A\|IDH3G\|ATP6\|COX2\|NDUFA9\|NDUFA10\|NDUFB3\|NDUFB4\|NDUFB10\|NDUFS1\|NDUFS2\|NDUFV1\|OGDH\|OXA1L\|PDHA1\|PDHB\|SDHA\|SDHB\|UQCRB\|UQCRC1\|UQCRC2\|UQCRFS1\|UQCRH\|SLC25A11\|SLC25A12\|SUCLG2\|SUCLA2\|COX5A\|ATP5MF\|SLC25A13\|ATP5PD\|ATP5MG\|NNT\|NDUFA13\|CYCS\|NDUFB11\|KGD4 |
| generation of precursor metabolites and energy | ACADM\|ACADVL\|ACAT1\|ACO2\|SLC25A4\|ATP5F1A\|ATP5F1B\|ATP5F1C\|ATP5PB\|ATP5ME\|ATP5PO\|COX4I1\|COX5B\|COX7A2\|COX7C\|CS\|CYC1\|DLAT\|DLD\|DLST\|ETFA\|ETFB\|FDXR\|FH\|GAA\|NIPSNAP2\|GPD2\|HK1\|IDH2\|IDH3A\|IDH3G\|ATP6\|COX2\|NDUFA9\|NDUFA10\|NDUFB3\|NDUFB4\|NDUFB10\|NDUFS1\|NDUFS2\|NDUFV1\|OGDH\|OXA1L\|OXCT1\|PDHA1\|PDHB\|PGD\|POR\|SDHA\|SDHB\|TKT\|UQCRB\|UQCRC1\|UQCRC2\|UQCRFS1\|UQCRH\|SLC25A11\|SLC25A12\|SUCLG2\|SUCLA2\|COX5A\|ATP5MF\|SLC25A13\|ATP5PD\|ATP5MG\|NNT\|NDUFA13\|CYCS\|NDUFB11\|KGD4 |
| aerobic respiration | ACO2\|ATP5F1A\|ATP5F1B\|ATP5F1C\|ATP5PB\|ATP5ME\|ATP5PO\|COX4I1\|COX5B\|COX7A2\|COX7C\|CS\|CYC1\|DLAT\|DLD\|DLST\|FH\|NIPSNAP2\|IDH2\|IDH3A\|IDH3G\|ATP6\|COX2\|NDUFA9\|NDUFA10\|NDUFB3\|NDUFB4\|NDUFB10\|NDUFS1\|NDUFS2\|NDUFV1\|OGDH\|OXA1L\|PDHA1\|PDHB\|SDHA\|SDHB\|UQCRB\|UQCRC1\|UQCRC2\|UQCRFS1\|UQCRH\|SUCLG2\|SUCLA2\|COX5A\|ATP5MF\|ATP5PD\|ATP5MG\|NNT\|NDUFA13\|CYCS\|NDUFB11\|KGD4 |
| energy derivation by oxidation of organic compounds | ACADM\|ACADVL\|ACO2\|ATP5F1A\|ATP5F1B\|ATP5F1C\|ATP5PB\|ATP5ME\|ATP5PO\|COX4I1\|COX5B\|COX7A2\|COX7C\|CS\|CYC1\|DLAT\|DLD\|DLST\|ETFA\|ETFB\|FH\|GAA\|NIPSNAP2\|GPD2\|IDH2\|IDH3A\|IDH3G\|ATP6\|COX2\|NDUFA9\|NDUFA10\|NDUFB3\|NDUFB4\|NDUFB10\|NDUFS1\|NDUFS2\|NDUFV1\|OGDH\|OXA1L\|PDHA1\|PDHB\|SDHA\|SDHB\|UQCRB\|UQCRC1\|UQCRC2\|UQCRFS1\|UQCRH\|SLC25A11\|SLC25A12\|SUCLG2\|SUCLA2\|COX5A\|ATP5MF\|SLC25A13\|ATP5PD\|ATP5MG\|NNT\|NDUFA13\|CYCS\|NDUFB11\|KGD4 |
| oxidative phosphorylation | ATP5F1A\|ATP5F1B\|ATP5F1C\|ATP5PB\|ATP5ME\|ATP5PO\|COX4I1\|COX5B\|COX7A2\|COX7C\|CYC1\|DLD\|NIPSNAP2\|ATP6\|COX2\|NDUFA9\|NDUFA10\|NDUFB3\|NDUFB4\|NDUFB10\|NDUFS1\|NDUFS2\|NDUFV1\|SDHA\|SDHB\|UQCRB\|UQCRC1\|UQCRC2\|UQCRFS1\|UQCRH\|COX5A\|ATP5MF\|ATP5PD\|ATP5MG\|NDUFA13\|CYCS\|NDUFB11 |
| electron transport chain | COX4I1\|COX5B\|COX7A2\|COX7C\|CYC1\|DLD\|ETFA\|ETFB\|GPD2\|COX2\|NDUFA9\|NDUFA10\|NDUFB3\|NDUFB4\|NDUFB10\|NDUFS1\|NDUF |

| | |
|---|---|
| | S2\|NDUFV1\|POR\|SDHA\|SDHB\|UQCRB\|UQCRC1\|UQCRC2\|UQCRFS1\|UQCRH\|SLC25A11\|SLC25A12\|COX5A\|SLC25A13\|CYCS |
| respiratory electron transport chain | COX4I1\|COX5B\|COX7A2\|COX7C\|CYC1\|DLD\|ETFA\|ETFB\|GPD2\|COX2\|NDUFA9\|NDUFA10\|NDUFB3\|NDUFB4\|NDUFB10\|NDUFS1\|NDUFS2\|NDUFV1\|SDHA\|SDHB\|UQCRB\|UQCRC1\|UQCRC2\|UQCRFS1\|UQCRH\|SLC25A11\|SLC25A12\|COX5A\|SLC25A13\|CYCS |
| proton transmembrane transport | SLC25A4\|SLC25A5\|ATP5F1A\|ATP5F1B\|ATP5F1C\|ATP5PB\|ATP5ME\|ATP5PO\|COX4I1\|COX5B\|CYC1\|LETM1\|ATP6\|COX2\|NDUFA9\|NDUFA10\|NDUFB3\|NDUFB4\|NDUFB10\|NDUFS1\|NDUFS2\|NDUFV1\|SLC25A3\|UQCRC1\|UQCRFS1\|UQCRH\|SLC25A12\|COX5A\|ATP5MF\|SLC25A13\|ATP5PD\|ATP5MG\|NNT\|GHITM |
| purine-containing compound metabolic process | ACAT1\|AK4\|ATP5F1A\|ATP5F1B\|ATP5F1C\|ATP5PB\|ATP5ME\|ATP5PO\|DLAT\|DLD\|DLST\|FDXR\|GPD2\|HK1\|IDH2\|ME1\|ATP6\|NDUFA9\|NDUFA10\|NDUFB3\|NDUFB4\|NDUFB10\|NDUFS1\|NDUFS2\|NDUFV1\|NT5E\|OGDH\|PDHA1\|PDHB\|PGD\|SDHA\|SDHB\|TKT\|SUCLG2\|SUCLA2\|ATP5MF\|NAMPT\|SLC25A13\|ATP5PD\|ATP5MG\|NNT\|ACOT9\|SAMHD1\|MACROD1\|AK3\|NDUFA13\|NDUFB11\|PARP9\|NADK2 |
| nucleobase-containing small molecule metabolic process | ACAT1\|AK4\|ATP5F1A\|ATP5F1B\|ATP5F1C\|ATP5PB\|ATP5ME\|ATP5PO\|CTPS1\|DLAT\|DLD\|DLST\|FDXR\|GPD2\|HK1\|IDH2\|ME1\|ALDH6A1\|ATP6\|NDUFA9\|NDUFA10\|NDUFB3\|NDUFB4\|NDUFB10\|NDUFS1\|NDUFS2\|NDUFV1\|NT5E\|OGDH\|PDHA1\|PDHB\|PGD\|SDHA\|SDHB\|TKT\|SUCLG2\|SUCLA2\|ATP5MF\|NAMPT\|SLC25A13\|ATP5PD\|ATP5MG\|NNT\|ACOT9\|SAMHD1\|MACROD1\|AK3\|NDUFA13\|NDUFB11\|CMAS\|PARP9\|NADK2 |
| nucleoside phosphate metabolic process | ACAT1\|AK4\|ATP5F1A\|ATP5F1B\|ATP5F1C\|ATP5PB\|ATP5ME\|ATP5PO\|CTPS1\|DLAT\|DLD\|DLST\|FDXR\|GPD2\|HK1\|IDH2\|ME1\|ATP6\|NDUFA9\|NDUFA10\|NDUFB3\|NDUFB4\|NDUFB10\|NDUFS1\|NDUFS2\|NDUFV1\|NT5E\|OGDH\|PDHA1\|PDHB\|PGD\|SDHA\|SDHB\|TKT\|SUCLG2\|SUCLA2\|ATP5MF\|NAMPT\|SLC25A13\|ATP5PD\|ATP5MG\|NNT\|ACOT9\|SAMHD1\|AK3\|NDUFA13\|NDUFB11\|CMAS\|PARP9\|NADK2 |
| carboxylic acid metabolic process | ACADM\|ACADVL\|ACAT1\|ACAT2\|ACO2\|BCAT2\|CPT1A\|CRAT\|CS\|DECR1\|DLAT\|DLD\|DLST\|ECH1\|ECHS1\|ETFA\|ETFB\|FH\|GLS\|GLUD1\|HADHA\|HADHB\|HK1\|IDH2\|IDH3A\|IDH3G\|ME1\|MGST3\|ALDH6A1\|OAT\|OGDH\|PDHA1\|PDHB\|PGD\|PTGS1\|SDHA\|SHMT2\|TST\|ALDH4A1\|SUCLG2\|SUCLA2\|LYPLA1\|ILVBL\|HIBADH\|PTGR1\|MTHFD1L\|ABHD12\|ACAD9\|PYCARD\|PYCR2\|FAHD2A\|HACD3\|CMAS\|MCCC1\|KGD4 |
| nucleoside phosphate biosynthetic process | ACAT1\|AK4\|ATP5F1A\|ATP5F1B\|ATP5F1C\|ATP5PB\|ATP5ME\|ATP5PO\|CTPS1\|DLAT\|DLD\|HK1\|ME1\|ATP6\|NDUFA9\|NDUFA10\|NDUFB3\|NDUFB4\|NDUFB10\|NDUFS1\|NDUFS2\|NDUFV1\|PDHA1\|PDHB\|SDHA\|SDHB\|ATP5MF\|NAMPT\|SLC25A13\|ATP5PD\|ATP5MG\|AK3\|NDUFA13\|NDUFB11\|CMAS\|PARP9\|NADK2 |
| purine-containing compound biosynthetic process | ACAT1\|AK4\|ATP5F1A\|ATP5F1B\|ATP5F1C\|ATP5PB\|ATP5ME\|ATP5PO\|DLAT\|DLD\|ATP6\|NDUFA9\|NDUFA10\|NDUFB3\|NDUFB4\|NDUFB10\|NDUFS1\|NDUFS2\|NDUFV1\|NT5E\|PDHA1\|PDHB\|SDHA\|SDHB\|ATP5MF\|NAMPT\|SLC25A13\|ATP5PD\|ATP5MG\|AK3\|NDUFA13\|NDUFB11\|PARP9\|NADK2 |
| mitochondrion organization | SLC25A4\|SLC25A5\|SLC25A6\|COX7A2\|NIPSNAP2\|HSPA9\|HSPD1\|LETM1\|NDUFA9\|NDUFA10\|NDUFB3\|NDUFB4\|NDUFB10\|NDUFS1\|NDUFS2\|OXA1L\|TFAM\|UQCRFS1\|VDAC2\|AIFM1\|LONP1\|AFG3L2\|PHB2\|MTCH1\|MTCH2\|SAMM50\|GHITM\|ACAD9\|NDUFA13\|CHCHD2\|TA |

| | |
|---|---|
| | CO1\|NDUFB11\|ATAD3A\|DNAJC11\|AGK\|TOMM22\|SLIRP\|PNPT1\|NDUFAF2\|COX20\|TOMM5 |
| nucleoside triphosphate biosynthetic process | AK4\|ATP5F1A\|ATP5F1B\|ATP5F1C\|ATP5PB\|ATP5ME\|ATP5PO\|CTPS1\|ATP6\|NDUFA9\|NDUFA10\|NDUFB3\|NDUFB4\|NDUFB10\|NDUFS1\|NDUFS2\|NDUFV1\|SDHA\|SDHB\|ATP5MF\|SLC25A13\|ATP5PD\|ATP5MG\|AK3\|NDUFA13\|NDUFB11 |
| proton motive force-driven mitochondrial ATP synthesis | ATP5F1A\|ATP5F1B\|ATP5F1C\|ATP5PB\|ATP5ME\|ATP5PO\|ATP6\|NDUFA9\|NDUFA10\|NDUFB3\|NDUFB4\|NDUFB10\|NDUFS1\|NDUFS2\|NDUFV1\|SDHA\|SDHB\|ATP5MF\|ATP5PD\|ATP5MG\|NDUFA13\|NDUFB11 |
| aerobic electron transport chain | COX4I1\|COX5B\|COX7A2\|COX7C\|CYC1\|DLD\|COX2\|NDUFA9\|NDUFA10\|NDUFB3\|NDUFB4\|NDUFB10\|NDUFS1\|NDUFS2\|NDUFV1\|SDHA\|SDHB\|UQCRB\|UQCRC1\|UQCRC2\|UQCRFS1\|UQCRH\|COX5A\|CYCS |
| ATP synthesis coupled electron transport | COX4I1\|COX5B\|COX7A2\|COX7C\|CYC1\|DLD\|COX2\|NDUFA9\|NDUFA10\|NDUFB3\|NDUFB4\|NDUFB10\|NDUFS1\|NDUFS2\|NDUFV1\|SDHA\|SDHB\|UQCRB\|UQCRC1\|UQCRC2\|UQCRFS1\|UQCRH\|COX5A\|CYCS |
| mitochondrial ATP synthesis coupled electron transport | COX4I1\|COX5B\|COX7A2\|COX7C\|CYC1\|DLD\|COX2\|NDUFA9\|NDUFA10\|NDUFB3\|NDUFB4\|NDUFB10\|NDUFS1\|NDUFS2\|NDUFV1\|SDHA\|SDHB\|UQCRB\|UQCRC1\|UQCRC2\|UQCRFS1\|UQCRH\|COX5A\|CYCS |
| purine nucleotide metabolic process | AK4\|ATP5F1A\|ATP5F1B\|ATP5F1C\|ATP5PB\|ATP5ME\|ATP5PO\|FDXR\|GPD2\|HK1\|IDH2\|ME1\|ATP6\|NDUFA9\|NDUFA10\|NDUFB3\|NDUFB4\|NDUFB10\|NDUFS1\|NDUFS2\|NDUFV1\|NT5E\|OGDH\|PGD\|SDHA\|SDHB\|TKT\|ATP5MF\|NAMPT\|SLC25A13\|ATP5PD\|ATP5MG\|NNT\|SAMHD1\|AK3\|NDUFA13\|NDUFB11\|PARP9\|NADK2 |
| ATP biosynthetic process | ATP5F1A\|ATP5F1B\|ATP5F1C\|ATP5PB\|ATP5ME\|ATP5PO\|ATP6\|NDUFA9\|NDUFA10\|NDUFB3\|NDUFB4\|NDUFB10\|NDUFS1\|NDUFS2\|NDUFV1\|SDHA\|SDHB\|ATP5MF\|SLC25A13\|ATP5PD\|ATP5MG\|NDUFA13\|NDUFB11 |
| proton motive force-driven ATP synthesis | ATP5F1A\|ATP5F1B\|ATP5F1C\|ATP5PB\|ATP5ME\|ATP5PO\|ATP6\|NDUFA9\|NDUFA10\|NDUFB3\|NDUFB4\|NDUFB10\|NDUFS1\|NDUFS2\|NDUFV1\|SDHA\|SDHB\|ATP5MF\|ATP5PD\|ATP5MG\|NDUFA13\|NDUFB11 |
| mitochondrial transport | SLC25A4\|SLC25A5\|SLC25A6\|CPT1A\|HSPA9\|HSPD1\|LETM1\|SLC25A3\|SLC25A1\|TST\|VDAC2\|SLC25A11\|SLC25A12\|AIFM1\|PMPCB\|SLC25A13\|TIMM44\|AFG3L2\|MTCH1\|MTCH2\|SAMM50\|GHITM\|SLC25A24\|NDUFA13\|AGK\|TOMM22\|SFXN3\|PNPT1\|SFXN1\|TOMM5 |
| ribonucleoside triphosphate biosynthetic process | ATP5F1A\|ATP5F1B\|ATP5F1C\|ATP5PB\|ATP5ME\|ATP5PO\|CTPS1\|ATP6\|NDUFA9\|NDUFA10\|NDUFB3\|NDUFB4\|NDUFB10\|NDUFS1\|NDUFS2\|NDUFV1\|SDHA\|SDHB\|ATP5MF\|SLC25A13\|ATP5PD\|ATP5MG\|NDUFA13\|NDUFB11 |

**Supplementary Results Table 10: Top 25 GO enriched pathways associated with downregulated DEPs in HPAECs, induced by connection of the vascular with the respiratory chip under basal conditions.**

| Description | Hits |
|---|---|
| extracellular matrix organization | SERPINH1\|COL5A2\|LOX\|MMP14\|P4HA1\|PLG\|SERPINF2\|PTX3\|TGFBI\|VTN\|ADAM15\|FLOT1\|CRTAP\|POSTN\|FKBP10\|COLGALT1\|SH3PXD2B |
| extracellular structure organization | SERPINH1\|COL5A2\|LOX\|MMP14\|P4HA1\|PLG\|SERPINF2\|PTX3\|TGFBI\|VTN\|ADAM15\|FLOT1\|CRTAP\|POSTN\|FKBP10\|COLGALT1\|SH3PXD2B |
| external encapsulating structure organization | SERPINH1\|COL5A2\|LOX\|MMP14\|P4HA1\|PLG\|SERPINF2\|PTX3\|TGFBI\|VTN\|ADAM15\|FLOT1\|CRTAP\|POSTN\|FKBP10\|COLGALT1\|SH3PXD2B |
| negative regulation of protein metabolic process | ACO1\|SERPINH1\|EIF4G1\|IPO5\|CAPRIN1\|NPM1\|SERPINF2\|PTX3\|THBS1\|UCHL1\|VTN\|SQSTM1\|CNOT9\|ITM2B\|CRTAP\|GTPBP4\|PDCD4\|RTRAF\|TBC1D24\|P3H1\|DNAJC1\|VPS25 |
| collagen fibril organization | SERPINH1\|COL5A2\|LOX\|P4HA1\|SERPINF2\|CRTAP\|FKBP10\|COLGALT1 |
| tube morphogenesis | CCN2\|TYMP\|FN1\|GJA1\|LOX\|MMP14\|MTHFD1\|SERPINF2\|TGFBI\|TGFBR2\|THBS1\|ADAM15\|NRP1\|BCL10\|PLXNB2\|PARVA\|FKBP10\|HHIP\|CTHRC1\|CLEC14A |
| regulation of cell-substrate adhesion | CRK\|FN1\|LIMS1\|MMP14\|PLG\|THBS1\|GFUS\|VTN\|ADAM15\|NRP1\|POSTN\|FERMT2 |
| regulation of hydrolase activity | SERPINH1\|CRK\|GSN\|STMN1\|LIMS1\|NPM1\|PCNA\|SERPINF2\|PTX3\|SOD1\|THBS1\|VTN\|NCSTN\|PLXNB2\|RCN3\|AGRN |
| supramolecular fiber organization | CAPG\|SERPINH1\|CFL2\|COL5A2\|DPYSL3\|GSN\|KRT2\|KRT10\|STMN1\|LOX\|P4HA1\|SERPINF2\|PRKAR1A\|PPFIA1\|CRTAP\|ARHGAP17\|FKBP10\|COLGALT1 |
| blood vessel morphogenesis | CCN2\|TYMP\|FN1\|GJA1\|LOX\|MMP14\|SERPINF2\|TGFBI\|TGFBR2\|THBS1\|ADAM15\|NRP1\|PARVA\|FKBP10\|CLEC14A |
| cell-substrate adhesion | CDH11\|CCN2\|FN1\|RAB1A\|TRIP6\|VTN\|PPFIA1\|ADAM15\|FERMT2\|PARVA |
| regulation of cytoskeleton organization | CAPG\|CFL2\|CRK\|CCN2\|GSN\|STMN1\|NUBP1\|NPM1\|SERPINF2\|PPFIA1\|NRP1\|FERMT2\|SYNPO\|PHLDB1\|GIT1\|ARHGAP17 |
| ossification | CAT\|CDH11\|COL5A2\|CCN2\|FASN\|FHL2\|GJA1\|LOX\|MMP14\|GTPBP4\|GIT1\|CTHRC1 |
| regulation of phosphorylation | EIF4G1\|FN1\|IPO5\|NPM1\|SDCBP\|THBS1\|UCHL1\|VTN\|TNFRSF10B\|NRP1\|BCL10\|CNOT9\|FLOT1\|GTPBP4\|PLXNB2\|PDCD4\|RTRAF\|VPS25 |
| negative regulation of protein modification process | EIF4G1\|IPO5\|NPM1\|UCHL1\|SQSTM1\|CRTAP\|GTPBP4\|PDCD4\|RTRAF\|TBC1D24\|P3H1\|VPS25 |

| | |
|---|---|
| negative regulation of fibrinolysis | PLG\|SERPINF2\|THBS1\|VTN |
| regulation of cell-matrix adhesion | LIMS1\|MMP14\|THBS1\|GFUS\|ADAM15\|NRP1\|POSTN\|FERMT2 |
| blood vessel development | CCN2\|TYMP\|FN1\|GJA1\|LOX\|MMP14\|SERPINF2\|TGFBI\|TGFBR2\|THBS1\|ADAM15\|NRP1\|PARVA\|FKBP10\|CLEC14A |
| regulation of actin filament-based process | CAPG\|CFL2\|CRK\|CCN2\|GSN\|STMN1\|SERPINF2\|PPFIA1\|NRP1\|FERMT2\|SYNPO\|ARHGAP17\|ABRACL |
| positive regulation of supramolecular fiber organization | CFL2\|CCN2\|GSN\|SERPINF2\|NRP1\|FERMT2\|SYNPO\|GIT1\|COLGALT1 |
| negative regulation of catalytic activity | SERPINH1\|EIF4A2\|IPO5\|NPM1\|SERPINF2\|PTX3\|THBS1\|UCHL1\|VTN\|PDCD4\|RTRAF\|VPS25 |
| vasculature development | CCN2\|TYMP\|FN1\|GJA1\|LOX\|MMP14\|SERPINF2\|TGFBI\|TGFBR2\|THBS1\|ADAM15\|NRP1\|PARVA\|FKBP10\|CLEC14A |
| regulation of stress fiber assembly | CCN2\|STMN1\|SERPINF2\|PPFIA1\|NRP1\|FERMT2\|SYNPO |
| regulation of actin cytoskeleton organization | CAPG\|CFL2\|CRK\|CCN2\|GSN\|STMN1\|SERPINF2\|PPFIA1\|NRP1\|FERMT2\|SYNPO\|ARHGAP17 |
| cell-matrix adhesion | CDH11\|CCN2\|FN1\|TRIP6\|VTN\|PPFIA1\|ADAM15\|FERMT2 |

**Supplementary Results Table 11. Top 25 enriched GO pathways with associated upregulated DEPs in HPAECs following $H_2O_2$-induced epithelial injury in REVAS.**

| Description | Hits |
|---|---|
| blood vessel development | APOB\|BAK1\|COL1A1\|COL1A2\|COL5A1\|CCN2\|TYMP\|ENG\|GJA1\|IGFBP7\|MMP14\|MYH9\|SERPINE1\|SERPINF2\|TGFBI\|TGFBR2\|THBS1\|WARS1\|RNF213 |
| vasculature development | APOB\|BAK1\|COL1A1\|COL1A2\|COL5A1\|CCN2\|TYMP\|ENG\|GJA1\|IGFBP7\|MMP14\|MYH9\|SERPINE1\|SERPINF2\|TGFBI\|TGFBR2\|THBS1\|WARS1\|RNF213 |
| response to type I interferon | IFI27\|IFIT1\|MX1\|OAS1\|OAS2\|OAS3\|STAT1\|ISG15\|IFIH1 |
| antiviral innate immune response | IFIT2\|IFIT1\|IFIT3\|MX1\|OAS1\|OAS2\|OAS3\|RIGI\|IFIH1 |

| | |
|---|---|
| extracellular matrix organization | COL1A1\|COL1A2\|COL5A1\|COL6A1\|ENG\|FBLN1\|FMOD\|MMP14\|MYH11\|PLG\|SERPINF2\|PLOD2\|TGFBI\|POSTN |
| extracellular structure organization | COL1A1\|COL1A2\|COL5A1\|COL6A1\|ENG\|FBLN1\|FMOD\|MMP14\|MYH11\|PLG\|SERPINF2\|PLOD2\|TGFBI\|POSTN |
| external encapsulating structure organization | COL1A1\|COL1A2\|COL5A1\|COL6A1\|ENG\|FBLN1\|FMOD\|MMP14\|MYH11\|PLG\|SERPINF2\|PLOD2\|TGFBI\|POSTN |
| defense response to virus | IFI27\|IFIT2\|IFIT1\|IFIT3\|MX1\|MX2\|OAS1\|OAS2\|OAS3\|STAT1\|ISG15\|RIGI\|IFIH1\|PARP9 |
| negative regulation of viral process | FBLN1\|GSN\|IFIT1\|MX1\|OAS1\|OAS2\|OAS3\|STAT1\|ISG15\|IFIH1 |
| blood vessel morphogenesis | APOB\|BAK1\|CCN2\|TYMP\|ENG\|GJA1\|IGFBP7\|MMP14\|MYH9\|SERPINE1\|SERPINF2\|TGFBI\|TGFBR2\|THBS1\|WARS1\|RNF213 |
| response to virus | APOB\|IFI27\|IFIT2\|IFIT1\|IFIT3\|MX1\|MX2\|OAS1\|OAS2\|OAS3\|STAT1\|ISG15\|RIGI\|IFIH1\|PARP9 |
| tube morphogenesis | APOB\|BAK1\|CCN2\|TYMP\|ENG\|GJA1\|IGFBP7\|MMP14\|MYH9\|SERPINE1\|SERPINF2\|TGFBI\|TGFBR2\|THBS1\|WARS1\|RNF213\|HHIP\|CTHRC1 |
| interleukin-27-mediated signaling pathway | MX1\|OAS1\|OAS2\|OAS3\|STAT1 |
| negative regulation of cell-substrate adhesion | CDKN2A\|COL1A1\|FBLN1\|MMP14\|SERPINE1\|PLG\|THBS1\|POSTN |
| innate immune response | GSN\|HLA-A\|CFI\|IFI27\|IFIT2\|IFIT1\|IFIT3\|MX1\|MX2\|OAS1\|OAS2\|OAS3\|STAT1\|ISG15\|RIGI\|PARP14\|IFIH1\|PARP9 |
| regulation of viral process | FBLN1\|GSN\|IFIT1\|MX1\|OAS1\|OAS2\|OAS3\|STAT1\|ISG15\|IFIH1 |
| response to mechanical stimulus | FAS\|BAK1\|CNN2\|COL1A1\|COL6A1\|ENG\|GSN\|MMP14\|STAT1\|THBS1\|POSTN |
| regulation of body fluid levels | SERPINC1\|F5\|FBLN1\|GJA1\|MYH9\|OAS2\|SERPINE1\|PLG\|SERPINF2\|THBS1\|VCL\|PAPSS2\|MYL9 |
| negative regulation of viral genome replication | IFIT1\|MX1\|OAS1\|OAS2\|OAS3\|ISG15\|IFIH1 |
| cellular response to type I interferon | IFI27\|IFIT1\|OAS1\|OAS2\|OAS3\|STAT1\|IFIH1 |

| | |
|---|---|
| angiogenesis | CCN2\|TYMP\|ENG\|IGFBP7\|MMP14\|MYH9\|SERPINE1\|TGFBI\|TGFBR2\|THBS1\|WARS1\|RNF213 |
| collagen fibril organization | COL1A1\|COL1A2\|COL5A1\|COL6A1\|FMOD\|SERPINF2\|PLOD2 |
| response to wounding | SERPINC1\|CNN2\|COL5A1\|COL6A1\|CCN2\|ENG\|F5\|FBLN1\|MYH9\|PLG\|VCL\|PAPSS2\|MYL9 |
| positive regulation of interferon-beta production | OAS1\|OAS2\|OAS3\|ISG15\|RIGI\|IFIH1 |
| positive regulation of type I interferon production | OAS1\|OAS2\|OAS3\|STAT1\|ISG15\|RIGI\|IFIH1 |

**Supplementary Results Table 12. Top 25 enriched GO pathways with associated downregulated DEPs in HPAECs following $H_2O_2$-induced epithelial injury in REVAS.**

| **Description** | **Hits** |
|---|---|
| Metabolism of RNA | DHX15\|FBL\|HNRNPA2B1\|HNRNPD\|HNRNPU\|NCL\|PSMB5\|RAN\|RPL8\|RPL28\|RPS10\|RPS27A\|SRSF1\|SRSF3\|SRSF7\|EFTUD2\|SNRNP40\|HNRNPR\|NUDT21\|LSM4\|PRPF19 |
| Mitochondrial protein degradation | ACO2\|ALDH2\|DLD\|FH\|IDH2\|OXCT1\|PDHB\|SHMT2\|SSBP1 |
| mRNA Splicing - Major Pathway | DHX15\|HNRNPA2B1\|HNRNPU\|SRSF1\|SRSF3\|SRSF7\|EFTUD2\|SNRNP40\|HNRNPR\|LSM4\|PRPF19 |
| Metabolism of lipids | ACADM\|ACADVL\|AKR1B1\|DECR1\|FABP5\|FDXR\|GLA\|GLB1\|HEXB\|ME1\|OXCT1\|CTSA\|LGMN\|RAN\|GNPAT\|CPNE3\|THRAP3\|PTGR1 |
| mRNA Splicing | DHX15\|HNRNPA2B1\|HNRNPU\|SRSF1\|SRSF3\|SRSF7\|EFTUD2\|SNRNP40\|HNRNPR\|LSM4\|PRPF19 |
| Processing of Capped Intron-Containing Pre-mRNA | DHX15\|HNRNPA2B1\|HNRNPU\|SRSF1\|SRSF3\|SRSF7\|EFTUD2\|SNRNP40\|HNRNPR\|NUDT21\|LSM4\|PRPF19 |
| Neutrophil degranulation | ANXA2\|CTSB\|CTSD\|FABP5\|GAA\|GLA\|GLB1\|GNS\|HEXB\|CTSA\|PRCP\|SPTAN1\|XRCC5\|CPNE3 |
| Viral Infection Pathways | PARP1\|RCC1\|CTSL\|NCL\|PSMB5\|RAN\|RPL8\|RPL28\|RPS10\|RPS27A\|TUFM\|XRCC5\|TAF15\|TRIM28\|H2AC12\|H2BC26 |
| Cellular responses to stimuli | ACADVL\|ANXA2\|NQO1\|H2AX\|HSPE1\|LMNB1\|ME1\|PGD\|PSMB5\|RING1\|RPL8\|RPL28\|RPS10\|RPS27A\|TKT\|H2BC26 |
| Cellular responses to stress | ACADVL\|NQO1\|H2AX\|HSPE1\|LMNB1\|ME1\|PGD\|PSMB5\|RING1\|RPL8\|RPL28\|RPS10\|RPS27A\|TKT\|H2BC26 |

| | |
|---|---|
| Citric acid cycle (TCA cycle) | ACO2\|DLD\|FH\|IDH2\|SDHA |
| Nuclear events mediated by NFE2L2 | NQO1\|ME1\|PGD\|PSMB5\|RPS27A\|TKT |
| Aerobic respiration and respiratory electron transport | ACO2\|DLD\|FH\|IDH2\|ME1\|PDHB\|RPS27A\|SDHA\|NDUFAF2 |
| KEAP1-NFE2L2 pathway | NQO1\|ME1\|PGD\|PSMB5\|RPS27A\|TKT |
| Diseases of metabolism | DLD\|FDXR\|GAA\|GLB1\|GNS\|HEXB\|CTSA\|RPS27A |
| Diseases of carbohydrate metabolism | GAA\|GLB1\|GNS\|RPS27A |
| Regulation of pyruvate metabolism | DLD\|ME1\|PDHB\|RPS27A |
| Meiotic synapsis | H2AX\|LMNA\|LMNB1\|SMC3\|H2BC26 |
| Mitotic Anaphase | RCC1\|LMNA\|LMNB1\|PSMB5\|RAN\|RPS27A\|SMC3 |
| Mitotic Metaphase and Anaphase | RCC1\|LMNA\|LMNB1\|PSMB5\|RAN\|RPS27A\|SMC3 |
| M Phase | RCC1\|H2AX\|LMNA\|LMNB1\|PSMB5\|RAN\|RPS27A\|SMC3\|H2BC26 |
| Metabolism of carbohydrates | AKR1B1\|GAA\|GLB1\|GNS\|HEXB\|PGD\|RPS27A\|TKT |
| Glycosphingolipid catabolism | GLA\|GLB1\|HEXB\|CTSA |
| Trafficking and processing of endosomal TLR | CTSB\|CTSL\|LGMN |
| Keratan sulfate degradation | GLB1\|GNS\|HEXB |

**Supplementary Results Table 13. Top 25 enriched GO pathways with associated upregulated DEPs in HPAECs following $H_2O_2$-induced epithelial injury in REVAS, cultured with or without vascular mural cells.**

| Description | Hits |
|---|---|

| | |
|---|---|
| actin cytoskeleton organization | ACTA1\|ACTN4\|ACTN1\|ANXA1\|CFL2\|CNN2\|CSRP1\|DBN1\|DPYSL3\|FLNB\|MYH9\|MYH10\|MYH11\|PAWR\|PLEC\|SRC\|TPM1\|TPM3\|TPM4\|VASP\|TAGLN2\|PDLIM4\|PDLIM1\|PDLIM7\|TRIP10\|ACTR3\|ARPC2\|MYL9\|CAP1\|ARPC1A\|PDLIM5\|PALLD\|EPB41L3\|PDLIM3\|FHOD1\|INF2 |
| actin filament-based process | ACTA1\|ACTN4\|ACTN1\|ANXA1\|CFL2\|CNN2\|CSRP1\|DBN1\|DPYSL3\|FLNB\|MYH9\|MYH10\|MYH11\|PAWR\|PLEC\|SRC\|TPM1\|TPM3\|TPM4\|VASP\|TAGLN2\|PDLIM4\|PDLIM1\|PDLIM7\|TRIP10\|ACTR3\|ARPC2\|MYL9\|CAP1\|ARPC1A\|PDLIM5\|PALLD\|EPB41L3\|PDLIM3\|FHOD1\|INF2 |
| supramolecular fiber organization | ACTA1\|ACTN1\|CDKN2A\|CFL2\|CNN2\|CSRP1\|DBN1\|DPYSL3\|KRT7\|MAP4\|MYH11\|PAWR\|PLEC\|PLOD1\|PLOD2\|SRC\|TPM1\|TPM3\|TPM4\|VASP\|TAGLN2\|PDLIM1\|ACTR3\|ARPC2\|MYL9\|BASP1\|CAP1\|ARPC1A\|INF2\|COLGALT1\|EPPK1 |
| actin filament organization | ACTA1\|ACTN1\|CFL2\|CNN2\|DBN1\|DPYSL3\|PAWR\|PLEC\|SRC\|TPM1\|TPM3\|TPM4\|VASP\|TAGLN2\|PDLIM1\|ACTR3\|ARPC2\|MYL9\|CAP1\|ARPC1A\|INF2 |
| actomyosin structure organization | ACTA1\|CFL2\|CNN2\|CSRP1\|MYH9\|MYH10\|MYH11\|PLEC\|SRC\|TPM1\|PDLIM1\|MYL9\|EPB41L3 |
| muscle structure development | ACTA1\|ACTN4\|ACTN1\|ATP2A2\|CFL2\|CSRP1\|FHL1\|FLNB\|LGALS1\|MYH9\|MYH11\|PLEC\|TAGLN\|TPM1\|PDLIM4\|PDLIM1\|PDLIM7\|MYL9\|BASP1\|PDLIM5\|PDLIM3 |
| response to wounding | CNN2\|CSRP1\|CCN2\|DPYSL3\|FN1\|ITGB4\|MYH9\|PLEC\|SOD2\|SRC\|TPM1\|VCL\|PAPSS2\|MYL9\|MYOF\|AK3\|EPPK1 |
| cell-substrate junction assembly | ACTN1\|FN1\|ITGB4\|PLEC\|SRC\|TNS1\|FERMT2 |
| cell-substrate junction organization | ACTN1\|FN1\|ITGB4\|PLEC\|SRC\|TNS1\|FERMT2 |
| muscle cell development | ACTA1\|ACTN4\|ACTN1\|ATP2A2\|CFL2\|CSRP1\|MYH11\|PLEC\|TPM1\|MYL9\|PDLIM5 |
| regulation of cell-substrate adhesion | ACTN4\|CDKN2A\|COL8A1\|DUSP3\|FN1\|GBP1\|LIMS1\|SRC\|VCL\|ARPC2\|FERMT2\|MYADM |
| cellular component assembly involved in morphogenesis | ACTA1\|CFL2\|CSRP1\|ITGB4\|MYH11\|PLEC\|TPM1\|MYL9\|EPB41L3 |
| cellular anatomical entity morphogenesis | ACTA1\|CFL2\|CSRP1\|ITGB4\|MYH11\|PLEC\|TPM1\|MYL9\|EPB41L3 |
| myofibril assembly | ACTA1\|CFL2\|CSRP1\|MYH11\|PLEC\|TPM1\|MYL9 |
| wound healing | CNN2\|CSRP1\|FN1\|MYH9\|PLEC\|SRC\|TPM1\|VCL\|PAPSS2\|MYL9\|MYOF\|AK3\|EPPK1 |
| muscle cell differentiation | ACTA1\|ACTN4\|ACTN1\|ATP2A2\|CFL2\|CSRP1\|MYH9\|MYH11\|PLEC\|TPM1\|MYL9\|PDLIM5 |

| | |
|---|---|
| membrane organization | ATP2A2\|BAK1\|MYH9\|PLEC\|RAB7A\|NDRG1\|YKT6\|TMED10\|RER1\|EMC1\|EPB41L3\|MYOF\|SEC61A1\|ATL2\|TMEM43\|MYADM\|BROX\|RILPL1 |
| regulation of substrate adhesion-dependent cell spreading | ACTN4\|GBP1\|LIMS1\|ARPC2\|FERMT2\|MYADM |
| cell-cell adhesion | ANXA1\|CSRP1\|ICAM2\|ITGB4\|KRT18\|LGALS1\|LIMS1\|MYH9\|SRC\|VCL\|PDLIM1\|MYL9\|PDLIM5\|PALLD\|NEXN |
| actin filament bundle assembly | ACTN1\|DPYSL3\|PAWR\|SRC\|PDLIM1\|MYL9 |
| striated muscle cell development | ACTA1\|CFL2\|CSRP1\|MYH11\|PLEC\|TPM1\|MYL9\|PDLIM5 |
| actin filament bundle organization | ACTN1\|DPYSL3\|PAWR\|SRC\|PDLIM1\|MYL9 |
| response to xenobiotic stimulus | ACAA1\|BAK1\|EPHX1\|ACSL1\|NNMT\|POR\|ABCD3\|SOD2\|SRC\|STAT1\|SULT1A1\|SULT1B1\|RAB6C |
| positive regulation of cellular component biogenesis | BAK1\|CCN2\|DPYSL3\|LIMS1\|MMP1\|SRC\|TPM1\|VASP\|WARS1\|RAB7A\|ACTR3\|ARPC2\|FERMT2\|FHOD1 |
| regulation of actin filament-based process | ATP2A2\|BST2\|CFL2\|CNN2\|CCN2\|MYH9\|TPM1\|VASP\|ARPC2\|FERMT2\|FHOD1\|MYADM |

**Supplementary Results Table 14. Top 25 enriched GO pathways with associated downregulated DEPs in HPAECs following $H_2O_2$-induced epithelial injury in REVAS, cultured with or without vascular mural cells.**

| Description | Hits |
|---|---|
| mRNA metabolic process | CDC5L\|CSTF1\|CSTF2\|DDX5\|DHX9\|DHX15\|ETF1\|HNRNPA1\|HNRNPA2B1\|HNRNPAB\|HNRNPF\|HNRNPK\|HNRNPU\|EIF3E\|HNRNPM\|NONO\|NOVA2\|PCBP2\|SFPQ\|SNRNP70\|SNRPD3\|SSB\|STAT3\|SF1\|KHSRP\|EFTUD2\|SNRNP40\|EIF4A3\|HNRNPR\|SAP18\|SYNCRIP\|DDX17\|PRPF8\|NUDT21\|LSM6\|U2AF2\|SNRNP200\|TARDBP\|LSM4\|PRPF19\|LUC7L2\|RBM25\|CPSF7\|PHF5A\|TNKS1BP1\|HNRNPUL2 |
| mRNA processing | CDC5L\|CSTF1\|CSTF2\|DDX5\|DHX9\|DHX15\|HNRNPA1\|HNRNPA2B1\|HNRNPF\|HNRNPK\|HNRNPU\|HNRNPM\|NONO\|NOVA2\|SFPQ\|SNRNP70\|SNRPD3\|SF1\|KHSRP\|EFTUD2\|SNRNP40\|EIF4A3\|HNRNPR\|SAP18\|SYNCRIP\|DDX17\|PRPF8\|NUDT21\|LSM6\|U2AF2\|SNRNP200\|TARDBP\|LSM4\|PRPF19\|LUC7L2\|RBM25\|CPSF7\|PHF5A\|HNRNPUL2 |

| | |
|---|---|
| RNA splicing | CDC5L\|DDX5\|DHX9\|DHX15\|HNRNPA1\|HNRNPA2B1\|HNRNPF\|HNRNPK\|HNRNPU\|HNRNPM\|NONO\|NOVA2\|SFPQ\|SNRNP70\|SNRPD3\|SF1\|TAF15\|KHSRP\|EFTUD2\|SNRNP40\|EIF4A3\|HNRNPR\|SAP18\|SYNCRIP\|DDX17\|PRPF8\|LSM6\|U2AF2\|SNRNP200\|TARDBP\|LSM4\|PRPF19\|LUC7L2\|RBM25\|PHF5A\|HNRNPUL2 |
| RNA splicing, via transesterification reactions | CDC5L\|DDX5\|DHX9\|DHX15\|HNRNPA1\|HNRNPA2B1\|HNRNPF\|HNRNPK\|HNRNPU\|HNRNPM\|NOVA2\|SFPQ\|SNRNP70\|SNRPD3\|SF1\|KHSRP\|EFTUD2\|SNRNP40\|EIF4A3\|HNRNPR\|SYNCRIP\|DDX17\|PRPF8\|LSM6\|U2AF2\|SNRNP200\|LSM4\|PRPF19\|LUC7L2\|PHF5A\|HNRNPUL2 |
| RNA splicing, via transesterification reactions with bulged adenosine as nucleophile | CDC5L\|DDX5\|DHX9\|DHX15\|HNRNPA1\|HNRNPA2B1\|HNRNPF\|HNRNPK\|HNRNPU\|HNRNPM\|NOVA2\|SFPQ\|SNRNP70\|SNRPD3\|SF1\|EFTUD2\|SNRNP40\|EIF4A3\|HNRNPR\|SYNCRIP\|DDX17\|PRPF8\|LSM6\|U2AF2\|SNRNP200\|LSM4\|PRPF19\|LUC7L2\|PHF5A\|HNRNPUL2 |
| mRNA splicing, via spliceosome | CDC5L\|DDX5\|DHX9\|DHX15\|HNRNPA1\|HNRNPA2B1\|HNRNPF\|HNRNPK\|HNRNPU\|HNRNPM\|NOVA2\|SFPQ\|SNRNP70\|SNRPD3\|SF1\|EFTUD2\|SNRNP40\|EIF4A3\|HNRNPR\|SYNCRIP\|DDX17\|PRPF8\|LSM6\|U2AF2\|SNRNP200\|LSM4\|PRPF19\|LUC7L2\|PHF5A\|HNRNPUL2 |
| regulation of mRNA metabolic process | DDX5\|DHX9\|ELAVL1\|HNRNPA1\|HNRNPAB\|HNRNPD\|HNRNPK\|HNRNPU\|NCL\|NOVA2\|NPM1\|SNRNP70\|SF1\|TAF15\|KHSRP\|EIF4A3\|SAP18\|SYNCRIP\|DDX17\|IGF2BP2\|U2AF2\|TARDBP\|DAZAP1\|PRPF19\|RBM25\|TNKS1BP1 |
| regulation of DNA metabolic process | PARP1\|CTNNB1\|DHX9\|H1-0\|H1-2\|H1-3\|H1-4\|H1-5\|HNRNPA1\|HNRNPA2B1\|HNRNPD\|HNRNPU\|KPNA2\|LMNA\|MCM3\|NPM1\|PCNA\|PRKDC\|RPA2\|XRCC5\|DEK\|RUVBL1\|RUVBL2\|CCT5\|OTUB1\|SLFN11 |
| DNA metabolic process | PARP1\|CDC5L\|DDB1\|DHX9\|XRCC6\|HMGB2\|HMGA1\|HNRNPAB\|KPNA2\|MCM3\|NASP\|NME1\|NONO\|NPM1\|PCNA\|PRKDC\|RBBP4\|RBBP7\|RECQL\|RPA1\|RPA2\|SFPQ\|XRCC5\|RUVBL1\|SYNCRIP\|RUVBL2\|SWAP70\|PRPF19\|OTUB1\|TNKS1BP1\|SLFN11 |
| regulation of RNA splicing | DDX5\|HNRNPA1\|HNRNPF\|HNRNPK\|HNRNPU\|NCL\|NOVA2\|SNRNP70\|SF1\|RBM12\|SAP18\|DDX17\|U2AF2\|DAZAP1\|PRPF19\|RBM25\|AHNAK |
| regulation of mRNA processing | DDX5\|DHX9\|HNRNPA1\|HNRNPK\|HNRNPU\|NCL\|NOVA2\|SNRNP70\|SF1\|SAP18\|DDX17\|U2AF2\|DAZAP1\|PRPF19\|RBM25 |
| regulation of mRNA splicing, via spliceosome | DDX5\|HNRNPA1\|HNRNPK\|HNRNPU\|NCL\|NOVA2\|SNRNP70\|SF1\|SAP18\|DDX17\|U2AF2\|DAZAP1\|PRPF19\|RBM25 |
| protein-RNA complex assembly | DHX9\|EIF3E\|PRKDC\|SNRPD3\|XRCC5\|SF1\|RUVBL1\|EIF3F\|EIF3I\|PRPF8\|RUVBL2\|U2AF2\|SNRNP200\|LSM4\|PRPF19\|LUC7L2\|PHF5A |
| chromosome organization | PARP1\|DHX9\|XRCC6\|H1-0\|H1-2\|H1-3\|H1-4\|H1-5\|HMGB2\|HMGA1\|HNRNPA2B1\|KPNB1\|MCM3\|NASP\|PCNA\|PRKDC\|RECQL\|RPA1\|RPA2\|XRCC5\|RUVBL1\|NUDC\|RUVBL2\|TNKS1BP1 |
| protein-RNA complex organization | DHX9\|EIF3E\|PRKDC\|SNRPD3\|XRCC5\|SF1\|RUVBL1\|EIF3F\|EIF3I\|PRPF8\|RUVBL2\|U2AF2\|SNRNP200\|LSM4\|PRPF19\|LUC7L2\|PHF5A |

| | |
|---|---|
| ribonucleoprotein complex biogenesis | DHX9\|FBL\|EIF3E\|NPM1\|PRKDC\|SNRPD3\|XRCC5\|SF1\|RUVBL1\|EIF3F\|EIF3I\|EIF4A3\|DDX17\|PRPF8\|RUVBL2\|LSM6\|U2AF2\|SNRNP200\|LSM4\|PRPF19\|LUC7L2\|LAS1L\|PHF5A |
| negative regulation of mRNA metabolic process | DHX9\|ELAVL1\|HNRNPAB\|HNRNPD\|HNRNPK\|HNRNPU\|TAF15\|SAP18\|SYNCRIP\|IGF2BP2\|U2AF2\|TARDBP |
| chromatin looping | DDX5\|DHX9\|DHX15\|EIF4A1\|MCM3\|NSF\|PCBP2\|RECQL\|RUVBL1\|EIF4A3\|DDX17\|RUVBL2\|SNRNP200\|RNF213 |
| DNA geometric change | DHX9\|XRCC6\|HMGB2\|HMGA1\|HNRNPA2B1\|MCM3\|RECQL\|RPA1\|XRCC5\|RUVBL1\|RUVBL2 |
| regulation of innate immune response | PARP1\|CRK\|DHX9\|XRCC6\|GRB2\|GRN\|HMGB2\|HPX\|IFI16\|NONO\|PCBP2\|PRKDC\|SFPQ\|UFD1\|XRCC5\|YWHAE\|MATR3\|RBM14\|ERAP1\|PSPC1\|NPLOC4 |
| alternative mRNA splicing, via spliceosome | DDX5\|DHX9\|HNRNPU\|HNRNPM\|SFPQ\|DDX17\|HNRNPUL2 |
| DNA conformation change | DHX9\|XRCC6\|HMGB2\|HMGA1\|HNRNPA2B1\|MCM3\|RECQL\|RPA1\|XRCC5\|RUVBL1\|RUVBL2 |
| positive regulation of translation | DHX9\|ELAVL1\|HNRNPD\|HNRNPU\|EIF3E\|ITGA2\|KRT17\|NPM1\|PRKDC\|SSB\|EIF4A3\|SYNCRIP |
| regulation of telomere maintenance | PARP1\|CTNNB1\|HNRNPA1\|HNRNPA2B1\|HNRNPD\|HNRNPU\|LMNA\|XRCC5\|RUVBL1\|RUVBL2\|CCT5 |
| DNA repair | PARP1\|CDC5L\|DDB1\|XRCC6\|HMGB2\|HMGA1\|MCM3\|NONO\|NPM1\|PCNA\|PRKDC\|RECQL\|RPA1\|RPA2\|SFPQ\|XRCC5\|RUVBL1\|RUVBL2\|PRPF19\|OTUB1\|TNKS1BP1 |

**Supplementary Results Table 15. List of DEPs/DEGs shared between REVAS HPAEC proteomic dataset and proteomic endothelial COPD dataset[1] and two endothelial single-cell RNA seq datasets from COPD lung tissues[2,3]. Datasets were filtered using an adjusted p-value threshold to ensure statistical stringency; log2fc**

| DEG/DEP | CHIP | COPD |
|---|---|---|
| TYMP | 2.2 | -2.42702 |
| TINAGL1 | 1.95 | -0.93006 |
| **POSTN** | 1.68 | 0.400386 |
| ITM2B | 1.66 | -0.61233 |
| **PAWR** | 1.49 | 0.627124 |
| SYNGR2 | 1.38 | -0.7052 |
| SRPX | 1.31 | -1.01227 |
| HLA-A | 1.12 | -0.84254 |
| **ENG** | 1.05 | 0.385867 |

| GENE | CHIP | COPD |
|---|---|---|
| **NAMPT** | -0.49 | -0.69394 |
| AHNAK | -0.51 | 0.786834 |
| **ANXA2** | -0.51 | -0.94808 |
| **SRSF1** | -0.51 | -2.50903 |
| **AKR1B1** | -0.52 | -1.47247 |
| **RPL28** | -0.55 | -0.70764 |
| **RPL8** | -0.57 | -0.65008 |
| **FBL** | -0.61 | -0.66904 |
| **HSPE1** | -0.63 | -1.35175 |

| | | |
|---|---|---|
| **STAT1** | 1.01 | 1.03111 |
| **TGFBR2** | 0.97 | 0.983508 |
| **PKN1** | 0.94 | 1.089946 |
| **CSNK1A1** | 0.93 | 0.633234 |
| **DDX58** | 0.92 | 1.26529 |
| SDCBP | 0.9 | -1.05 |
| SERPINE1 | 0.89 | -0.84985 |
| **PARP14** | 0.83 | 0.54855 |
| SLC3A2 | 0.78 | -1.39449 |
| GSN | 0.69 | -0.85522 |
| ICAM1 | 0.67 | -2.73308 |
| SNCG | 0.57 | -1.34834 |
| **PAPSS2** | 0.44 | 0.497184 |
| **VCL** | 0.4 | 1.08822 |
| **MYH9** | 0.39 | 0.572995 |
| **DYNC1H1** | 0.18 | 1.06719 |
| MACF1 | -0.27 | 0.630522 |
| RACK1 | -0.32 | 1.15382 |
| **HNRNPA2B1** | -0.38 | -0.57948 |
| DDX17 | -0.39 | 0.4414 |
| **NCL** | -0.39 | -0.30298 |
| SPTAN1 | -0.4 | 1.1024 |
| **XRCC5** | -0.4 | -0.36056 |
| **HNRNPA1** | -0.46 | -5.86943 |
| THRAP3 | -0.46 | 0.568904 |
| DHX15 | -0.49 | 0.872977 |

| | | |
|---|---|---|
| **RAN** | -0.64 | -1.13763 |
| **CTSD** | -0.66 | -4.24394 |
| **LMNA** | -0.68 | -0.27096 |
| **RPS10** | -0.68 | -5.71828 |
| **ACADVL** | -0.69 | -0.90936 |
| **SRSF3** | -0.73 | -0.37045 |
| **RPS27A** | -0.77 | -0.64986 |
| **PSMB5** | -0.78 | -1.41957 |
| PRCP | -0.81 | 0.540785 |
| **ALDH2** | -0.82 | -1.01538 |
| NOVA2 | -0.82 | 0.631599 |
| **ADIRF** | -0.83 | -3.53103 |
| PPFIBP1 | -0.84 | 0.714224 |
| TAF15 | -0.86 | 1.19208 |
| **SRSF7** | -0.9 | -0.90944 |
| RCC1 | -0.93 | 2.5316 |
| **DEK** | -0.98 | -0.2909 |
| LUC7L2 | -1.02 | 0.769753 |
| HNRNPR | -1.11 | 0.336549 |
| PPM1F | -1.15 | 0.679908 |
| ITGA6 | -1.47 | 0.739669 |
| **CTSB** | -0.72 | -0.699 |
| MX1 | 1.27 | -0.31907 |
| **PGRMC2** | 1.34 | 0.376312 |
| **SIPA1** | 1.42 | 0.545246 |
| **DYSF** | -0.72 | -0.52518 |
| **IFI27** | 2.47 | 0.486152 |

### MATERIALS AND METHODS

#### Design of microfluidic device.

The microfluidic device was designed using the Computer Aided Design (CAD) software, AutoCAD 2020 (AutoDesk Inc, Ca, USA). All designs were drawn in 2D and consisted of top and bottom rectangular parts with a length of 32.0 mm and a width of 20.0 mm. Extra rectangles were added to what would be the final outline of the chip as a “safety zone” to reduce margin of error during the excision and cutting to size processes. The top part was designed with linear microchannels to enable continuous flow and with guideline markers for punching the access holes. The bottom part comprising multiple channels was designed to maximise the area of contact between cells grown in the top and bottom part of the device.

#### Chip fabrication

The photomasks were printed on high resolution photomask films (MicroLitho, UK). The process of chip manufacture is described in[4]. In short, PDMS was cast over the pre-prepared patterned silicon wafers, peeled off and top and bottom part of the REVAS were excised. Top and matching bottom parts of respiratory or vascular chips were aligned in conformal contact with a PET membrane placed in between.

To enable continuous perfusion with culture media, the channel inlets were connected to a flow circuit operated by a peristaltic pump.

#### Device cell count

For determining approximate cell count inside the device to evaluate how many cells from different cell types can potentially be hosted within the channels, confocal microscopy images were analysed and scaled using the image processing program ImageJ 2020 (NIH, University of Wisconsin, USA).

#### Flow simulation

The REVAS circuit was modelled in COMSOL as follows: two connected macrovascular endothelial channels (VAS-chip), linked to four microvascular endothelial channels (RE-chips). The detail of the model can be found in the **Supplemental Methods Table 1**. Flow simulations were performed to assess three elements: determine the flow rate at the inlet allowing the recapitulation of physiologically relevant wall shear stress (WSS), characterise the flow throughout the circuit, and verify the flow distribution after the branching.

**Supplemental Methods Table 1: Parameters of the PDMS wall model.** Simulation settings, including abbreviations and descriptions, were configured in COMSOL. Channels refer to macrovascular endothelial (VAS-chip) and microvascular endothelial (RE-chip) channels.

| Description | Parameter | Expression |
|---|---|---|
| Channel length | L | 27[mm] |
| Channel width | W | 1[mm] |
| Channel height | H | 0.2[mm] |
| Inlet/outlet diameter | $D_{i/o}$ | 1[mm] |
| Inlet/outlet height | $H_{i/o}$ | 1.5[mm] |
| Tube diameter | $D_{tube}$ | 1[mm] |

**Parameter setting**

The following parameters were chosen for the simulation. Domain (inside of the channel) was defined as water at 20 °C (dynamic viscosity $\mu = 1$ mPa·s; density $\rho = 997$ kg/m³), considered as an approximation of cell culture medium. As for the boundaries, PDMS and a PETwere selected according to their respective position in the device. Additionally, boundaries were defined with no slip condition.

Using these parameters, we calculated Reynolds numbers, using the Reynolds number formula for rectangular channels (utilising the hydraulic diameter). The Reynolds numbers were all under 2000, justifying the use of a laminar flow simulation.

Another parameter to input is inlet velocity, which was calculated based on the volumetric flow rate outputted by the pump ($Flow_{pump}$) and reported to the inlet of the REVAS circuit ($Flow_i$) and the surface area at the inlet, using **Equation 4.1** and **Equation 4.2**. The equation of the velocity is displayed in **Equation 4.3** and **Equation 4.4**.

$$Flow_i\left[\frac{m^3}{s}\right] = \frac{Flow_{pump}}{3.6 * 10^9} \quad \textbf{(4.1)} \qquad \text{and} \qquad Surface_{i/o}[m^2] = \frac{D^2{}_{i/o}}{4} \quad \textbf{(4.2)}$$

$$Velocity\left[\frac{m}{s}\right] = \frac{Flow_i}{Surface_{\frac{i}{o}}} \quad \textbf{(4.3)} \qquad \text{and} \qquad Velocity\left[\frac{m}{s}\right] = \frac{4Flow_i}{\pi D_{i/o}^2} \quad \textbf{(4.4)}$$

The final parameter to calculate is the pressure at the outlet ($P_{out}$), using the Hagen-Poiseuille equation (**Equation 4.5**). To improve the accuracy of pressure drop calculations, a tube extension (1 mm diameter, 25 cm length) was added at the outlet. This extension minimises the influence of the outlet boundary on the simulation. For this reason, $L_{tube}$ is 25 cm and $D_{tube}$ is 1 mm.

$$P_{out}[Pa] = \frac{128 * \mu * Flow_i * L_{tube}}{\pi * D_{tube}{}^4} \qquad (\mathbf{4.5})$$

**Meshing and numerical stability**

Mesh convergence studies were performed to ensure numerical accuracy while maintaining computational efficiency. Mesh convergence analysis was performed using five levels of refinement (coarse to extremely fine). An extra-fine mesh (~17 million elements) was selected, as the average wall shear stress (WSS) varied by less than 1 % with further refinement. The corresponding mesh is shown in **Supplemental Figure 1**.

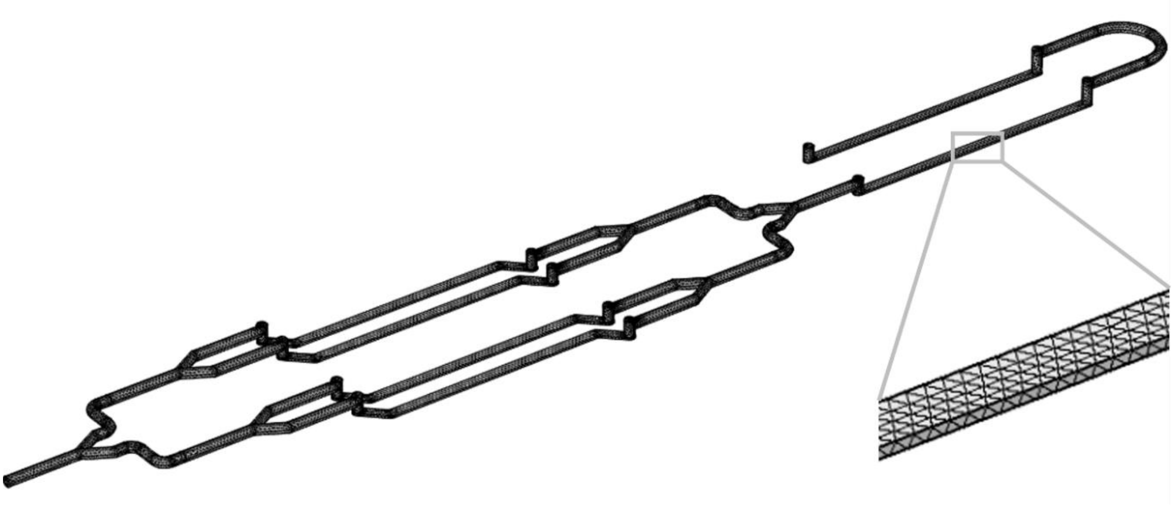

**Supplemental Methods Figure 1: Model of macrovascular end microvascular endothelial channels connected with tubing forming the REVAS circuit.** The model was constructed in COMSOL and a extra fine mesh was generated, composed of four distinct element types: tetrahedra, hexahedra, triangular prisms, and pyramids.

**Post-processing**

Wall shear stress (WSS) was calculated as:

WSS = viscosity of the fluid x shear rate

WSS = spf.mu*spf.sr (using the built-in COMSOL parameters)

WSS distributions were visualised along channel walls. For precise WSS plots, a 3D line was drawn along the PET membrane surface and 2D graph of the WSS along this line were plotted.

Velocity streamlines were plotted using 30 evenly spaced data points from the inlet to assess flow continuity and identify potential regions of disturbance or recirculation.

Cross-sectional WSS profiling showed a plateau-like distribution across the channel centre, where ~ 60 % of channel width was exposed to ≥ 95 % of the target WSS and ~80 % to ≥ 85 %, confirming spatially uniform shear exposure over most of the culture surface (**Supplementary Methods Figure 2**).

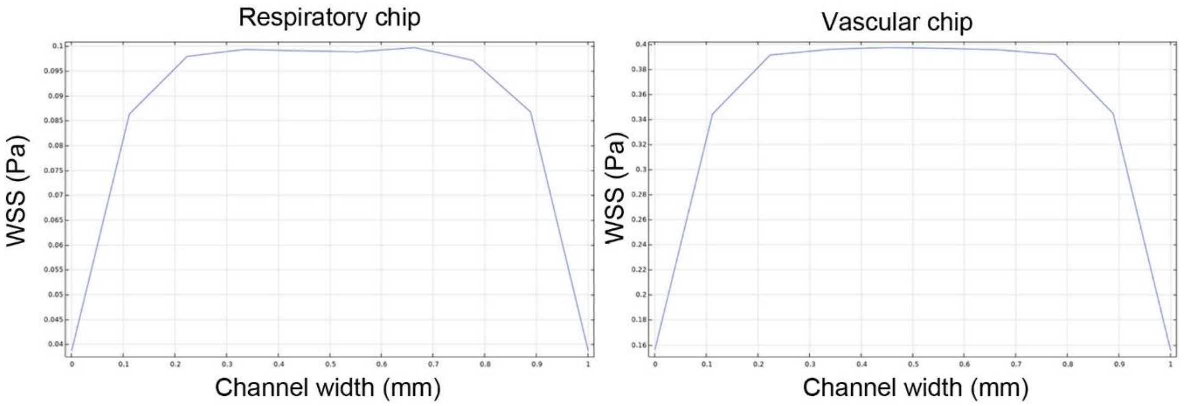

**Supplemental Methods Figure 2: Cross-sectional wall shear stress (WSS) distribution across REVAS microfluidic channels. (A)** Simulated WSS profile across the width of the respiratory channel demonstrating a distribution centred around the target shear stress of 1 dyne/cm² (0.1 Pa). **(B)** Simulated WSS profile across the width of the vascular channel demonstrating a distribution centred around the target shear stress of 4 dynes/cm² (0.4 Pa).

#### Characterisation of flow parameters

To confirm that the flow rates defined by in silico modelling were accurately delivered within the microfluidic system, flow rate measurements were performed using a Sensirion flow sensor (SLF3S-1300F).

After priming the system with distilled water, the vascular chip was linked to the reservoirs through tubing and perfused with distilled water for 1 minute at a flow rate of 160 µl/min, corresponding to a WSS of 4 dynes/cm². The flow sensor, connected via USB to a laptop, was positioned between the channel outlet and the reservoir, where waveform measurements were recorded. Data was collected and analysed using the Sensirion Data Viewer software (Sensirion AG, Switzerland).

**Cell culture**

All cells were maintained in T75 cell culture flasks (Sarstedt, Germany, Cat. No. 833911002) coated with 0.2 % porcine gelatine (Sigma, Cat. No. G1890) in a humidified incubator (37 °C, 5 % CO2). All media were sterile filtered using a SteriCup filter unit (Merck, USA, Cat No. SCGPU05RE) prior to use. After reaching confluence, cells were passaged at the 1:3 ratio using 0.05 % Trypsin-EDTA solution (Gibco, UK). Cells were used between passages 4 – 8 upon receipt from commercial suppliers. Culture media were changed every two days, unless stated otherwise. All donors are listed in **Supplementary Methods Table 2**.

Primary HPAEC culture. Human pulmonary artery endothelial cells (HPAEC; PromoCell, Germany, Cat. No. C-12241) from 3 different biological donors were cultured in Endothelial Growth Medium 2 (EGM-2, PromoCell, Germany C-22211), supplemented with 2 % Foetal Calf Serum (FCS) and growth supplement (PromoCell, Cat. No. C-22111) containing Epidermal Growth Factor (EGF, 5 ng/mL), Basic Fibroblast Growth Factor (FGF, 10 ng/mL), Insulin-like Growth Factor (IGF, 20 ng/L), Vascular Endothelial Growth Factor (VEGF, 0.5 ng/mL), Ascorbic Acid (1 μg/mL), Heparin (22.5 μg/mL), Hydrocortisone (0.2 μg/mL) and antibiotics: 1 % Streptomycin/Penicillin (100 μg/mL, Gibco; Cat. No. 15140-122), and 1 % MycoZap™ (Lonza, Cat. No. VZA-2032).

Primary HPASMC culture. Human pulmonary artery smooth muscle cells (HPASMC; Lonza, UK, Cat. No. CC-2581) from 3 different biological donors were cultured in Smooth Muscle Cell Growth Medium 2 (SmGM-2, PromoCell, Cat. No. C-22062), supplemented with 5 % FCS and growth supplement (PromoCell, Cat. No. C-39267) containing EGF (0.5 ng/mL), Basic FGF (2 ng/mL), insulin (5 μg/mL), 1 % Streptomycin/Penicillin (100 μg/mL) and 1 % MycoZap™.

Primary HPF culture. Human pulmonary fibroblasts (HPF; PromoCell, Cat. No. 12360) from 3 different biological donors were cultured in T75 tissue culture flasks in Fibroblast Growth Medium 2 (PromoCell, Cat. No. C-23020), supplemented with growth supplement (PromoCell, Cat. No. C-39325) containing FCS (0.02 mL/mL), Basic FGF (1 ng/mL), insulin (5 μg/mL), 1 % Streptomycin/Penicillin (100 μg/mL) and 1 % MycoZap™.

Primary HPC culture. Human pericytes (HPC; PromoCell, Cat. No. C-12980) from 3 different biological donors were cultured in Pericyte Growth Medium 2 (PromoCell, Cat. No. C-28041), with growth supplement (PromoCell, Cat. No. C-39267) containing FCS (0.02 mL/mL), Basic

FGF (1 ng/mL), insulin (5 µg/mL), 1 % Streptomycin/Penicillin (100 µg/mL) and 1 % MycoZap™.

Primary HPMVEC culture. Human pulmonary microvascular endothelial cells (HPMVEC; PromoCell, Cat. No. C-12281) from 3 different biological donors were cultured in T75 tissue culture flasks in Endothelial Cell Growth Medium MV2 (PromoCell, Cat. No. C-22221) supplemented with growth supplement mix (PromoCell, Cat. No. C-39226) containing FCS (0.05 mL/mL), EGF (5 ng/mL), FGF (10 ng/mL), (IGF, 20 ng/L), (VEGF, 0.5 ng/mL), Ascorbic Acid (1 µg/mL) and Hydrocortisone (0.2 µg/mL), with added antibiotics: 1 % Streptomycin/Penicillin (100 µg/mL) and 1 % MycoZap™.

Primary HsAEpC culture. Human small airway epithelial cells (HsAEpC; Lonza, Cat. No. CC-2547) from 3 different biological donors were cultured in Small Airway Epithelial Cell Growth Medium (Lonza, Cat. No. CC-3118), with added growth supplement (Lonza, Cat. No. C-4124) containing Bovine Pituitary Extract (BPE, 4 µl/mL), hydrocortisone (1 µl/mL), hEGF (1µl/mL), Retinoic Acid (1 µl/mL), triiodothyronine (1 µl/mL), Gentamicin sulface-Amphotericin (GA, 1 µl/mL), Fatty Acid Free – Bovine Serum Albumin (BSA-FAF, 10 µl/mL) and antibiotics: 1 % Streptomycin/Penicillin (100 µg/mL), and 1 % MycoZap™.

HULEC-5a cell culture. HULEC-5a is a lung microvascular endothelial cell line (HMEC; ATCC, Cat. No. CRL-3244). Cells were cultured in MCDB131 medium (Merck, Cat. No. M8537) substituted with FCS (0.1 mL/mL), L-Glutamine (0.01 mmol/mL; Merck, Cat. No. G5792), Hydrocortisone (Stemcell, Cat. No. 07926; 1 µg/mL), EGF (10ng/mL) and antibiotics: 1 % Streptomycin/Penicillin (100 µg/mL) and 1 % MycoZap™.

**Supplementary Methods Table 2. Human pulmonary artery endothelial, smooth muscle cells, fibroblasts, pericytes, microvascular endothelial, small airway epithelial cells used in experiments.** The table shows donor's age and gender. All donors were non-smokers.

| Donor number | Age | Gender |
|---|---|---|
| HPAECs | | |
| Donor 1 | 53 | Male |
| Donor 2 | 51 | Female |
| Donor 3 | 51 | Female |
| HPASMCs | | |
| Donor 1 | 51 | Female |

| | | |
|---|---|---|
| Donor 2 | 64 | Male |
| Donor 3 | 52 | Female |
| HPFs | | |
| Donor 1 | 57 | Female |
| Donor 2 | 58 | Male |
| Donor 3 | 62 | Female |
| HPCs | | |
| Donor 1 | 0 | Female |
| Donor 2 | 0 | Female |
| Donor 3 | 0 | Female |
| HPMVECs | | |
| Donor 1 | 57 | Female |
| Donor 2 | 62 | Male |
| Donor 3 | 57 | Female |
| HsAEpCs | | |
| Donor 1 | 45 | Male |
| Donor 2 | 38 | Female |
| Donor 3 | 32 | Male |

Co-culture medium in REVAS. The co-culture medium, previously used in pulmonary-artery-on-a-chip to support endothelial-smooth muscle co-culture [5], consisted of a basal endothelial cell medium supplemented with 10 % FBS, 2.5 ng/mL epidermal growth factor, 10 ng/mL fibroblast growth factor, 20 ng/mL insulin-like growth factor, 0.5 ng/mL vascular endothelial growth factor, 1 % Streptomycin/Penicillin (100 µg/mL), and 1 % MycoZap™. To prevent excessive proliferation of HPFs within the REVAS system, the FBS was reduced to 5 % as suggested by Lopez-Martinez et al.[6]. The following EGM2 supplements were excluded to avoid dysregulated proliferation of HPASMCs, HPFs and HPCs: VEGF, heparin, ascorbic acid, and hydrocortisone [7-10].

**Air-liquid interface**

The protocol of cell culture at an air-liquid interface requires two sets of media: one for the seeding/submerged phase and one for the air-exposed part. Reagents are listed in **Supplementary Methods Table 3 and 4,** respectively.

For 500 mL of media for submerged cultures, 250 mL of Airway Epithelial Growth Medium was mixed with 250 mL of DMEM and supplemented with HEPES (5.96 mg/mL), BSA (1.5 µg/mL), BPE (4 µl/mL), rhEGF (10 ng/mL), Hydrocortisone (0.5 µg/mL), Epinephrine (0.5 µg/mL), rhTransferrin (10 µg/mL), Retinoic Acid (0.1 ng/mL), 1 % Streptomycin/Penicillin (100 µg/mL) and 1 % MycoZap™.

**Supplementary Methods Table 3. DMEM ALI Mix.** Needed for the seeding/submerged phase of transwell inserts.

| Reagent | Manufacturer | Cat. No. |
|---|---|---|
| Airway Epithelial Cell Growth Medium – Growth Medium Kit | PromoCell | C-21160 |
| DMEM, high glucose, GlutaMAX™ Supplement, pyruvate | Gibco / Thermo Fisher | 31966047 |
| Bovine serum albumin (BSA) | Sigma | A7030-100G |
| HEPES solution 1 M, pH 7.0-7.6 | Sigma | H0887-100ML |
| Penicillin-Streptomycin (lab stock) | - | - |

For 500 mL of media for air-exposed cultures, 450 mL of PneumaCult™-ALI Basal Medium was supplemented with 50 mL of PneumaCult™-ALI 10X Supplement, 5 vials of PneumaCult™-ALI Maintenance Supplement, Heparin (4 µg/mL), Hydrocortisone (0.48 µg/mL), 1 % Streptomycin/Penicillin (100 µg/mL) and 1 % MycoZap™.

**Supplementary Methods Table 4. PneumaCult ALI complete.** Needed for air-exposed transwell inserts.

| Reagent | Manufacturer | Cat. No. |
|---|---|---|
| PneumaCult™-ALI Medium | Stemcell | #05001 |
| Hydrocortisone Stock Solution | Stemcell | #07926 |
| Heparin Solution | Stemcell | #07980 |
| Penicillin-Streptomycin (lab stock) | - | - |

To set up an air-liquid interface (ALI), 6.5 mm Transwell inserts with 0.4 µm pore size polyester membranes (Corning, USA, Cat. No. 3470) were coated with sterile 0.2 % gelatin for

1 hour. HsAEpCs (30,000 cells) were suspended in 200 µl of the pre-mixed airway epithelial growth medium and seeded onto the PET membrane in each insert. The basal compartment was filled with 500 µl of medium, and the medium was changed every two days until the cells reached confluence. Once confluence was achieved, the medium was removed from the apical compartment to expose the cells to air. Pre-mixed PneumaCult-ALI medium was added to the basal compartment only, and the medium was changed every two days to maintain the cells at the air-liquid interface until full differentiated.

48 hours before the experiment's completion, the Transwell inserts were carefully inverted, and 30,000 HPMVECs suspended in 200 µl of "co-culture medium" were seeded onto the opposite side of the PET membrane. After a 2-hour incubation at 37°C, the inserts were reverted and placed back for further culture.

**qPCR**

RNA isolation. Prior to RNA isolation, all surfaces and pipettes were treated with RNase-Zap RNase Decontamination Solution (Cat: AM9780; Invitrogen, UK) and sprayed with 70 % ethanol. Cells were washed once with PBS and treated with trypsin (Cat. No. 25200072; Gibco, UK) for 5-7 mins at 37 °C to facilitate cell detachment. Once the cells rounded up and detached, trypsin was neutralised with DMEM containing 10 % FBS, and the resulting cell suspension was collected in RNase-free Low Bind 1.5 mL microcentrifuge tubes (Cat. No. Z666548-250EA; Merck Life Science, UK). Cells were centrifuged at 5000xg for 5 minutes at room temperature, and the residual medium was carefully aspirated without disturbing the cell pellet. The tubes were then either stored at -80 °C or used immediately for RNA isolation. Total RNA was extracted using either Monarch Total RNA Miniprep Kit (Cat. No. T2010S; New England Biolabs, MA, USA) or RNeasy Plus Micro Kit (Cat. No. 74034; Qiagen, Germany), following the manufacturers' protocols, with a modification of reducing the recommended centrifugation speed by half in all steps except for the final wash and elution in RNase-free water. Double elutions were performed to maximise RNA recovery from the spin columns. Total RNA levels were quantified using a NanoDrop™ ND2000 spectrophotometer (Thermo Scientific, UK). The RNA purity was evaluated by the absorbance ratio at 260 nm and 280 nm (A260/A280), with values above 1.8 considered acceptable. The RNA was then stored at -80 °C for further experiments.

Reverse transcription. 50 ng of RNA extracted from cells grown in chips or 100 ng of RNA extracted from any other samples was reverse transcribed into cDNA using the LunaScript Reverse Transcription SuperMix Kit (Cat. No. E3010L; New England Biolabs, MA, USA). Samples were kept on ice for room temperature in PCR-certified 96-well plates, with a total reaction volume of 20 µl. The resulting cDNA samples were diluted with RNase-free water to achieve a cDNA concentration of 1 ng/µl. 30 µl was added to the chip cDNA samples, while 70 µl was added for all other samples. The cDNA solutions were either stored at -20 °C or were used immediately for qPCR. Components of the reverse transcription reaction mix, and parameters of reverse transcription cycle are outlined in **Supplemental Methods Table 5** and **Table 6** below.

**Supplemental Methods Table 5. Reverse Transcription reaction mix.**Negative controls using the No-Reverse Transcription Control Mix (5x) were included in each plate to check for DNA/RNA contamination. cDNA synthesis was carried out using a SimpliAmp™ Thermal Cycler (Applied Biosystems, UK).

| Component | 20 µl Reaction | Final Concentration |
|---|---|---|
| LunaScript Reverse Transcription SuperMix (5x) | 4 µl | 1x |
| RNA sample | variable | 50/100 ng |
| Nuclease-free water | Remainder up to 20 µl | - |

**Supplemental Methods Table 6. Reverse Transcription Cycling Conditions.**

| Cycle Step | Temperature | Time | Cycles |
|---|---|---|---|
| Primer Annealing | 25 °C | 2 minutes | 1 |
| cDNA Synthesis | 55 °C | 10 minutes | 1 |
| Heat Inactivation | 95 °C | 1 minute | 1 |

qPCR primer design. All primer sequences were designed using FASTA sequences from PubMed, NCBI. Briefly, coding sequences (CDS) were copied and pasted into the Primer3web website (https://primer3.ut.ee). Primer pairs of identical lengths were preferably selected, and their sequences were input into Primer-BLAST (NCBI) to verify gene specificity. Primer pairs were retained only if they specifically matched the gene of interest. Selected primers (DNA oligos) were ordered from Sigma-Aldrich, reconstituted in RNase-free water conferring to the manufacturer's instructions, and stored at -20 °C. A list of primers used for the different cell types can be found in **Supplemental Methods Table 7** below.

**Supplemental Methods Table 7. List of qPCR primers sequences used for each gene of interest for different cell types**.

| **Gene**<br>Cell type | **Forward Primer** | **Reverse Primer** |
|---|---|---|
| ***ACTA2***<br>HPAEC, SMC | TCCCAGGGCTGTTTTCCCAT | TCTGTGCTTCGTCACCCACG |
| ***BAK1***<br>ALL | GCCACCAGCCTGTTTGAGAGT | GACCATTGCCCAAGTTCAGGG |
| ***BRD4***<br>ALL | CGTCAAGCTGAACCTCCCTGATTA | ATGATCTCGGTTTCTTCTGTGGGT |
| ***CASP9***<br>ALL | GCTCTTCCTTTGTTCATCTCCTGC | TCTGCATTTCCCCTCAAACTCTCA |
| ***CCL5***<br>ALL | CTCGCTGTCATCCTCATTGCTACT | AGTTGATGTACTCCCGAACCCATT |
| ***CDC45***<br>ALL | CTGAAGCAGGTGAAGCAGAAGTTC | AGAGCCTGGATGAAGTGATCTGTC |
| ***CDH5***<br>HPAEC, HPMVEC | CCTCCGATACATGAGCCCTCC | TCGGAAGAACTGGCCCTTGT |
| ***CNN1***<br>SMC | CAGAAGTATGACCACCAGCG | ATTGATCTTCTTCACGGAGCC |
| ***COL1A1***<br>HPF | TGGCGAGAGAGGTGAACAAG | TCACCCTTAGCACCATCGTT |
| ***eNOS***<br>HMEC | ATCCCCCGGAGAATGGAGAG | AGCGGATCTTATAACTCTTGTGCT |
| ***E-Selectin***<br>HMEC | CTGGGCTCCAGGTGAACCCAAC | CCGTGGCCACTGCAGGATGTA |
| ***FOXJ1***<br>HsAEpC | GAGAGTCCCCAGGTACAACAAAT | ATTTTGGTCTCCTCTCTCTGCCC |
| ***GAPDH***<br>housekeeping | CGGATTTGGTCGTATTGGGCG | GCCTTCTCCATGGTGGTGAAGAC |
| ***ICAM1***<br>ALL | GGCCAGCTTATACACAAGAACCAG | GTGCCATCCTTTAGACACTTGAGC |
| ***IL6***<br>ALL | TGAGAGTAGTGAGGAACAAGCCA | TTGTGGTTGGGTCAGGGGTG |

| | | |
|---|---|---|
| ***KLF2***<br>HPAEC | GTGAGAAGCCCTACCACTGCAACT | CCGGTTCTCTGGGTCCAATAAATA |
| ***KRT5***<br>HsAEpC | CTCAAGGATGCCAGGAACAAGCTG | TCCACTGAGTCTGCATTCCTCG |
| ***MUC5AC***<br>HsAEpC | CAGTCTGGGGTCCTCATTCAGC | TCAGCTTGGTGTTGTGGGAGAG |
| ***NG2***<br>HPC | ACTCAGCAGTCAGACCCTCAG | AATATTOCCAGOGTAGACCTCTGC |
| ***PDGFRα***<br>HPF | TTACCCTGGAGAAGTGAAAGGCAA | GGCTGAAGGTGGGTTTGATTTCAA |
| ***PDGFRβ***<br>HPC | TGTGGGAACGGATGTCCCAG | ATTAGGGAGGAAGCCCACGG |
| ***PECAM**-1*<br>HPAEC | TGGAAGGAGTGCCCAGTCCCA | CGGAAGGATAAAACGCGGTCCTG |
| ***POSTN***<br>ALL | CCTAATTCCTGATTCTGCCAAACA | GCAGAATTAATTTAAGGAGGCGCT |
| ***P-Selectin***<br>HMEC | GGCATAGCATCACTTCCTACTCCA | TCCTATGAGTGTGAATCCAGCGTT |
| ***SCGB1A1***<br>HsAEpC | CTTTCAGCGTGTCATCGAAACCC | ATGATGCTTTCTCTGGGCTTTTGG |
| ***TAGLN***<br>HPAEC, SMC | AACAGCCTGTACCCTGATGGC | GATCTCCACGGTAGTGCCCAT |
| ***TJP1***<br>HsAEpC | GCTCAAGAGGAAGCTGTGGGTAAC | GAAGCGTCACTGTATGTTGTTCCC |
| ***VCAM1***<br>ALL | AAAATCGAGACCACCCCAGAATCT | ACAGGTAAGAGTGTTCGTTCCCAA |
| ***VEGFA***<br>ALL | AGGGAAAGGGGCAAAAACGAAAG | ACAAATGCTTTCTCCGCTCTG |
| ***VIM***<br>HPAEC, HPF | ACTCGGTGGACTTCTCGCTG | TCCCGCATCTCCTCCTCGTA |
| ***VWF***<br>HPAEC, HPMVEC | ATGTGGCGTTCGTCCTGGAA | TTGGACTGTGCCTCGCTGAA |
| ***YHWAZ***<br>housekeeping | CCGAGCCAGCAGAACATCCA | GAGACGACCCTCCAAGATGACCTA |

qPCR. Samples were prepared on ice. 9 µl of Luna Universal qPCR Master Mix (New England Biolabs, MA, USA; Cat. No. M3003X) and 1 µl of 1 ng/µl cDNA (prepared as delineated in Chapter 2.4.2) were added to each well of a 384-well skirted PCR plate (StarLab, UK, Cat. No.

E1042-3840). For each gene of interest, a negative control was included by adding 1 µl of nuclease-free water instead of cDNA, along with 9 µl of the appropriate qPCR master mix. Details of the reagents used in the reactions are described in **Supplemental Methods Table 8.**

**Supplemental Methods Table 8. qPCR master mix components**.

| Component | 10 µl Reaction | Final Concentration |
|---|---|---|
| Luna Universal qPCR Master Mix | 5µl | 1x |
| 10 µM forward primer | 0.5 µl | 0.25 µM |
| 10 µM reverse primer | 0.5 µl | 0.25 µM |
| Nuclease-free water | 3 µl | - |
| cDNA products | 1 µl | 1 ng |

The plates were sealed with MicroAmp Optical Adhesive Film (Thermo Fisher Scientific, Cat. No. 4311971), inverted to mix the samples, and centrifuged at 1000xg for 1 minute. qPCR was conducted using a QuantStudio 12K Flex Real-Time PCR System (Applied Biosystems, USA). Amplification was performed over 40 cycles, as outlined in **Supplemental Methods Table 9.**

**Supplemental Methods Table 9.** qPCR cycling conditions. qPCR amplifications were performed with a QuantStudio 12K Flex Real-Time PCR System (Applied Biosystems, USA).

| Cycle Stage | Temperature | Time | Cycles |
|---|---|---|---|
| **Hold Stage** | 50 °C | 2 minutes | 1 |
| | 95 °C | 5 minutes | |
| **PCR Stage** | 95 °C | 15 seconds | 40 |
| | 62 °C | 30 seconds | |
| **Melt Curve Stage** | 95 °C | 15 seconds | 1 |
| | 60 °C | 1 minute | |
| | 95 °C | 15 seconds | |

qPCR Data analysis. The relative gene expression was determined by the comparative ΔCT method/ 2-ΔΔCT method [283]. Genes were normalised to Tyrosine 3-Monooxygenase/Tryptophan 5-Monooxygenase Activation Protein Zeta (*YHWAZ*) expression. The fold change relative to the control group for each sample was plotted with the data presented as individualised data points. Gene expression between two different groups was compared using unpaired t-tests. For comparisons involving more than two groups, one- or two-way ANOVA was used as appropriate, followed by post hoc testing (Tukey's or Sidak's test) to correct for multiple comparisons. Data analyses were performed using PRISM 9 (GraphPad 2023), and statistical significance was set at $P < 0.05$.

#### TaqMan qPCR

Gene expression analysis of differentiation markers in cells from monocultures and co-cultures was conducted using qPCR. Cells were collected from PET membranes by trypsinisation, and RNA was extracted using the Monarch Total RNA Miniprep Kit (New England BioLabs, UK, Cat. No. T2010S) as per manufacturers' instructions. cDNA was synthesised using the LunaScript RT SuperMix Kit (NEB, Cat. No. E3010S) on the SimpliAmp™ Thermal Cycler (ThermoFisher, Cat. No. A24811). The cDNA synthesis protocol included a primer annealing step at 25 °C for 2 minutes, followed by reverse transcription at 55 °C for 10 minutes, and heat inactivation at 95 °C for 1 minute.

qPCR was performed using a ViiA 7 Real-Time PCR System (ThermoFisher). Primer mixes were prepared by combining 12 µl of TaqMan primers (ThermoFisher) as listed in **Supplementary Methods Table 10**, 12 µl of dd$H_2O$, and 120 µl of TaqMan Mastermix (ThermoFisher, Cat. No. 4305719). For each reaction, 3.6 µl of this primer mix was combined with 2.4 µl of cDNA, normalised to 1 ng/µl. The qPCR was run for a total of 45 cycles. The cycling conditions included a hold stage at 50 °C for 2 minutes, followed by 20 seconds at 95 °C. The PCR stage consisted of 1 second at 95 °C and 20 seconds at 60 °C for extension.

The relative gene expression was determined by the comparative ΔCT method/ $2^{-\Delta\Delta CT}$ method. Genes were normalised to Tyrosine 3-Monooxygenase/Tryptophan 5-Monooxygenase Activation Protein Zeta (*YHWAZ*) expression.

**Supplementary Methods Table 10: TaqMan primer list.**

| Gene | Assay ID | Gene | Assay ID |
|---|---|---|---|
| *YWHAZ* | Hs01122445_g1 | *MUC5B* | Hs00861595_m1 |
| *FOXJ1* | Hs00230964_m1 | *TJP1* | Hs01551861_m1 |

| | | | |
|---|---|---|---|
| *KLF2* | Hs00360439_g1 | *TJP2* | Hs00910543_m1 |
| *KRT5* | Hs00361185_m1 | *TJP3* | Hs00274276_m1 |
| *MUC5AC* | Hs01365616_m1 | *TP63* | Hs00978340_m1 |

**Immunocytochemistry (ICC)**

Cells were fixed in 4 % paraformaldehyde (Sigma Aldrich, UK, Cat. No. 1004968350) solution in PBS for 20 min, washed three times in PBS, permeabilised in 0.1 % Triton-X-100 (Sigma Aldrich, UK, Cat. No. X100-5ML) solution in PBS for 5 min, washed thrice in PBS and incubated in 2 % BSA (Sigma Aldrich, UK, Cat. No. A9418) blocking solution in PBS for 30 min. Cells were then incubated for 2 hours at room temperature or overnight at 4 °C with primary antibodies (Table 2.6) at appropriate dilutions in 0.1 % BSA. Following the incubation with primary antibodies, cells were washed 3 times in PBS and then incubated with fluorescently labelled secondary antibodies, as appropriate for 2 hours at room temperature in the dark (**Supplementary Methods Tables 11**). Following another three washes with PBS, samples were mounted in VECTASHIELD® Antifade Mounting Medium with DAPI (Vector Laboratories, Cat No. H-1200). Hoechst 33342 (Invitrogen, Cat No. H3570) was also added at a final concentration of 10 µg/mL to enhance nuclear staining. Images were taken under a STELLARIS 8 Confocal Microscope (Leica Microsystems).

**Supplementary Methods Tables 11. Primary and secondary antibodies.**

| Antibody Name | Species | Company | Cat No | Final Conc. |
|---|---|---|---|---|
| Anti-NG2 | Rabbit | Abcam | ab129051 | 5 µg/mL |
| Anti-TAGLN | Rabbit | Abcam | ab14106 | 2 µg/mL |
| Calponin-1 | Rabbit | Sigma-Aldrich | ABT129 | 2 µg/mL |
| FOXJ1 | Goat | R&D Systems, Inc | AF3619 | 10 µg/mL |
| Human alpha-Smooth Muscle Actin (Alexa Fluor™ 594 Conjugated) | Mouse | R&D Systems, Inc | IC1420T | 1 µg/mL |
| KRT5 (Alexa Fluor® 647) | Rabbit | Abcam | ab193895 | 5 µg/mL |
| Mucin 5AC antibody (Alexa Fluor® 555) | Rabbit | Abcam | ab218714 | 10 µg/mL |

| | | | | |
|---|---|---|---|---|
| ZO-1 (Alexa Fluor™ 488) | Mouse | Thermo Fisher Scientific | ZO1-1A12 | 10 µg/mL |
| Phalloidin-TRITC | Amanita phalloides (Death Cap Mushroom) | Tocris Bioscience | 5783 | 6.6 µM |
| VE-cadherin (Alexa Fluor™ 488 Conjugated) | Mouse | eBioscience | 53-1449-42 | 10 µg/mL |
| Alexa fluor-546 (Anti-Rabbit) | Goat | Life Technologies | A11010 | 10 µg/mL |
| Alexa Fluor Plus 594 (Anti-Mouse) | Goat | Life Technologies | A32742 | 3.33 µg/mL |
| Alexa Fluor Plus 594 (Anti-Rabbit) | Goat | Life Technologies | A32740 | 3.33 µg/mL |
| anti-Rabbit IgG (H+L) Highly Cross-Adsorbed Secondary Antibody (Alexa Fluor™ Plus 647) | Goat | Invitrogen | A32733 | 1 µg/mL |

**H&E staining**

To perform H&E (Hematoxylin & Eosin) staining on HsAEpCs and HPMVECs, the Transwell inserts were fixed in 200 µl of 4 % paraformaldehyde (PFA) for 20 minutes and then washed with 200 µl of PBS. The membranes were carefully cut out and placed apical side up into a 24-well plate containing 4 % low melting point agarose (ThermoFisher, Cat. No. R0801) in PBS, pre-heated to 37 °C. The plate was placed on ice until the agarose solidified. The agarose-embedded samples were then sectioned into 4 µm slices and stained with H&E by the Research Histology Facility at the South Kensington Campus. Images were captured using the Aperio Versa 8 (Leica Biosystems, Germany).

**Permeability assays**

Endothelial and epithelial barrier function was evaluated by measuring passage of fluorescent dextran through cell monolayer growing on top of the porous PET membrane in Transwell filters or in microfluidic chips. In microfluidic chips, a 1 mg/mL solution of 40 kDa FITC-Dextran (Sigma Aldrich, Dorset, UK, Cat. No. FD40S) was introduced into the co-culture medium and perfused through the upper (endothelial or epithelial) channel at shear stress of 4 dynes/cm² for vascular and 0.0001 dynes/cm² for respiratory channels, respectively. After 1 hour, the medium from the bottom channel was collected by flushing the channel with 250 µl of medium, which was collected at the channel outlet. For permeability experiments involving thrombin, 1 U/mL thrombin (Sigma Aldrich, Dorset, UK, Cat. No. T7513) was added to the co-culture medium containing 1 mg/mL FITC-Dextran, and the experiment was conducted following the same protocol. Measurements of the amount of FITC-dextran that passed through the membrane were performed using a GLOMAX spectrophotometer (Promega, USA) with excitation/emission wavelengths of 490/525 nm. The apparent permeability (Papp [cm/s]) was determined using the formula described in[11]

**Analysis of alignment**

Cell alignment relative to flow was quantified using the "Draw Max/Min Ferets Tool" macro from the ROI toolbox developed by Stephen Rothery (FILM, Imperial College London). This tool measures the Feret angle of cells within a selected region of interest. In brief, cell nuclei from images of HPAECs stained with DAPI were marked using the "threshold" and "watershed" functions. Regions of interest (ROIs) were generated using the "analyse particle" function, with 20/30 cell nuclei captured in each image. The "Draw Max/Min Ferets Tool" was then applied, and the measurements were exported to Excel for further analysis. Cells with a Feret angle of less than 30° relative to the horizontal axis, were considered aligned with the flow. For consistency, all images were oriented so that the bottom edge aligned with the flow direction.

**Cytokine profiling of REVAS on culture medium**

Cytokine profiling was performed on culture medium collected from the REVAS system after 48 hours of co-culture, with a total of n=5 per experimental group. This included comparison between untreated REVAS controls and $H_2O_2$-treated REVAS systems, both in the presence of all vascular and respiratory cell types. Samples were sent to Eve Technologies (Canada) for quantification using their 48-plex Human Cytokine Assay platform. For each sample, 100 µl of culture medium was provided in 0.5 mL snap cap vials. Two pilot samples were included, one expected to exhibit low cytokine levels and another with predicted high cytokine content, for initial quality assurance.

The fold change was calculated relative to the REVAS control group and plotted in bar-charts. For the heatmap, z-score of the concentration was plotted. Some cytokines had concentrations below the limit of detection (OOR; out of range), and these were assigned a concentration of 0 pg/mL according to the provider's guidelines. Statistical comparisons were conducted using two-tailed unpaired t-tests. Where unequal variances were identified between groups, Welch's correction was applied.

**Comparative analysis of REVAS and COPD datasets**

Pulmonary endothelial transcriptomic data was isolated from four publicly available scRNA-seq datasets [12-15]. Data were filtered to include only age-matched donors (50-76 years old). Three main endothelial cell populations were identified: general capillary cells, aerocytes, and arterial endothelial cells. Analysis was performed using the Seurat pipeline, with differential gene expression calculated using the Wilcoxon two-sample test and Bonferroni correction. Differentially expressed genes (DEGs) between control and COPD samples were selected based on adjusted P-value < 0.05 and a $\log_2$ fold change > 0.6 (corresponding to >1.5-fold change) [16].

These DEG lists were then compared with significantly dysregulated proteins (adj P < 0.05) identified in the in vitro HPAEC proteomic dataset following REVAS experimentation. The differentially expressed genes for the endothelial cell subtypes were combined into one dataset. Where the same gene was differentially expressed in multiple endothelial cell subtypes, the average $\log_2$ fold change was calculated. The direction of fold change stayed the same. The datasets were overlapped to find genes/proteins differentially expressed in both datasets. The shared genes and their respective $\log_2$ fold changes are shown in a heatmap. Overlaps and unique features were visualised using a Venn diagram to highlight shared and cell-type-specific responses.

**Statistical analysis and data presentation**

All experiments were performed at least in triplicate, with three biological repeats, unless otherwise stated. The number of replicates for each experimental type (e.g., gene expression, proteomics, permeability) is specified in respective figure legends. Graphs were generated using GraphPad Prism 9 software (GraphPad Software Inc., CA, USA), with statistical tests performed, as appropriate. Error bars in bar graphs represent the standard error of the mean (SEM). Data normality was assessed using the Shapiro-Wilk test. For comparisons between two groups, normally distributed data were analysed using unpaired Student's t-tests. For comparisons involving more than two groups, one-way or two-way ANOVA was used as appropriate, followed by post hoc testing (Tukey's or Sidak's test) to correct for multiple comparisons. A significance threshold was set at $P < 0.05$.

Gene expression between two groups (e.g., chip vs transwell) was analysed using unpaired t-tests unless stated otherwise. For functional studies comparing effects of stimuli such as thrombin or $H_2O_2$, or comparing monoculture vs co-culture conditions, group comparisons were made using ANOVA frameworks to account for multiple experimental groups.
